# simPIC: flexible simulation of single-cell ATAC-seq paired-insertion counts from individuals to populations

**DOI:** 10.1101/2025.09.21.676689

**Authors:** Sagrika Chugh, Heejung Shim, Davis J. McCarthy

**Affiliations:** Bioinformatics and Cellular Genomics, St. Vincent’s Institute of Medical Research, 9 Princes Street, Fitzroy, 3065, Victoria, Australia; School of Mathematics and Statistics, Faculty of Science, University of Melbourne, Grattan Street, Parkville, 3010, Victoria, Australia; Faculty of Medicine, Dentistry and Health Sciences, University of Melbourne, Grattan Street, Parkville, 3010, Victoria, Australia

**Author notes:** Contributing authors.

**Keywords:** single-cell ATAC-seq, single-cell genomics, software, simulation

## Abstract

Single-cell Assay for Transposase Accessible Chromatin (scATAC-seq) is increasingly used at population scale to study how genetic variation shapes chromatin accessibility. Method development is limited by the lack of flexible simulation tools with known ground truth. Here, we present simPIC, a fast, memory-efficient framework for simulating realistic single-cell ATAC-seq count data across individuals and populations. simPIC models cell groups, batch effects, and genotype-dependent accessibility variation, enabling controlled evaluation of population-scale methods, including chromatin accessibility quantitative traits locus (QTL) mapping. Across multiple datasets and cell types, simPIC closely matches real data distributions while scaling to cohort sizes impractical for current tools.

## 1 Background

The epigenetic template governing lineage-specific programs is shaped by a complex landscape of cis-regulatory elements. Single-cell Assay for Transposase-Accessible Chromatin sequencing (scATAC-seq) has become foundational for mapping chromatin accessibility at single-cell resolution, enabling the study of regulatory interactions and transcriptional control [1]. By using Tn5 transposase to preferentially insert sequencing adapters into accessible genomic regions, scATAC-seq provides detailed maps of cell-specific regulatory landscapes. This approach provides critical insights into the interplay between chromatin structure and gene expression [1, 2]. However, extracting robust biological insights from scATAC-seq remains computationally challenging; the data is characterised by extreme sparsity and high-dimensional feature spaces. Consequently, most analytical strategies to date have focused on clustering cells into discrete groups or aggregating accessibility signals across genomic peaks. While effective in many contexts, these approaches may obscure the biological variability present at the single-cell level. New data analysis approaches are needed, but the computational infrastructure for scATAC-seq data analysis remains underdeveloped relative to rapid advances in experimental techniques [3]. Notably, flexible and biologically realistic simulation frameworks are lacking, which constrains rigorous evaluation and benchmarking of novel computational methods.

As experimental designs transition toward larger cohorts and multi-condition studies, the requirement for rigorous methodological validation has become acute. In the absence of comprehensive experimental “gold standards” where regulatory events are validated *a priori* simulation provides a practical solution by generating synthetic datasets with known ground truth under controlled settings. Such frameworks enable objective assessment of whether a method can faithfully recapitulate empirical data distributions, preserve latent biological signals, and maintain computational scalability, which are core criteria used in systematic benchmarking studies [4]. While the transcriptomics field utilizes mature, parametric simulators such as Splatter [5], and its population extension splatPop [6], which provide transparent parametric models calibrated to real data, analogous tools for scATAC-seq remain fragmented across several distinct methodological approaches.

Existing scATAC-seq simulators fall into two broad categories, each with distinct strengths and limitations. The first comprises tools purpose-built for scATAC-seq. SCAN-ATAC-Sim [7] derives synthetic profiles by downsampling bulk chromatin accessibility data, masking cell-type-specific heterogeneity. simATAC [8] generates bin-by-cell count matrices rather than peak-by-cell matrices, produces pseudo-bulk profiles, and does not adequately capture the extreme sparsity characteristic of singlecell data [9]. EpiAnno is constrained by a strict dependency on prior knowledge of highly accessible peaks, restricting its use to datasets with established reference annotations [9]. simCAS [10] uses a deep-learning embedding strategy that reproduces the high-level properties of real data but requires extensive parameter calibration and utilises a mechanistically opaque architecture, which limits the user’s ability to precisely specify biological variables. scReadSim [11] simulates reads rather than count matrices directly. It learns local genomic distributions from aligned reads and can generate realistic count matrices; however, it requires raw BAM files as input, which are not readily available. Critically, none of these purpose-built tools models genetic effects such as chromatin accessibility quantitative trait loci (caQTLs), limiting their applicability in population-scale studies where inter-individual genetic variation substantially shapes chromatin accessibility landscapes.

The second category comprises general-purpose multi-modal simulators that include scATAC-seq as one of several supported data types. scDesign3 [12] uses a copula-based statistical framework to model multivariate single-cell data across RNA, ATAC, and spatial modalities; it offers broad distributional flexibility but is not tailored to the discrete count structure of paired insertion counts (PIC) quantified scATAC-seq data [13] and does not support population-scale simulation with genotype effects. scMultiSim [14] is a mechanistic simulator guided by gene regulatory networks (GRN) and cell-cell interactions, whose primary strength is benchmarking multi-omics integration and GRN inference methods; it similarly lacks support for inter-individual genetic variation. DiTSim [15] uses diffusion-transformer architectures to learn and reproduce the global distributional structure of real scATAC-seq datasets and demonstrates strong marginal fidelity; however, as a deep-generative model, it does not expose an interpretable parametric structure and does not directly support user-specified experimental designs such as batch structure or caQTL.

Across existing scATAC-seq simulation frameworks, methods typically do not simultaneously provide an interpretable and calibrated generative model of count data, flexible experimental design, and population-scale simulation with explicit genetic effects. This limitation is increasingly consequential as functional genomics shifts toward population-scale studies that require modelling of inter-individual variation in chromatin accessibility. As such, including caQTL effects is essential for evaluating methods that aim to identify genetic drivers of regulatory variation and complex traits. To address the need for realistic and efficient simulation of single-cell chromatin accessibility data, we developed simPIC, an R/Bioconductor package that generates peak-by-cell count matrices based on PIC, a recently introduced standardized approach for quantifying scATAC-seq signal [16]. PIC quantifies scATAC-seq signal by counting the number of Tn5 insertion events within each peak per cell, yielding a count structure constrained by the underlying diploid biology. The resulting counts are well suited to probabilistic modelling of sparsity and accessibility. simPIC learns key data properties from a reference dataset and uses these calibrated parameters to generate synthetic count matrices under user-specified experimental designs. The simPIC framework supports multiple cell types, experimental batch effects, and through integration with the splatPop framework [6] enables population-scale simulation with caQTL effects, a capability that is not supported in existing scATAC-seq simulation frameworks. This integration is crucial for mimicking the multidimensional heterogeneity present in real single-cell studies, where technical artifacts, diverse cellular identities, and inter-individual genetic variation all shape the observed chromatin landscape [17]. simPIC is well documented and freely available through Bioconductor, providing researchers with a flexible, scalable, and biologically grounded tool for simulating realistic scATAC-seq datasets (Figure1).

**Fig. 1:**
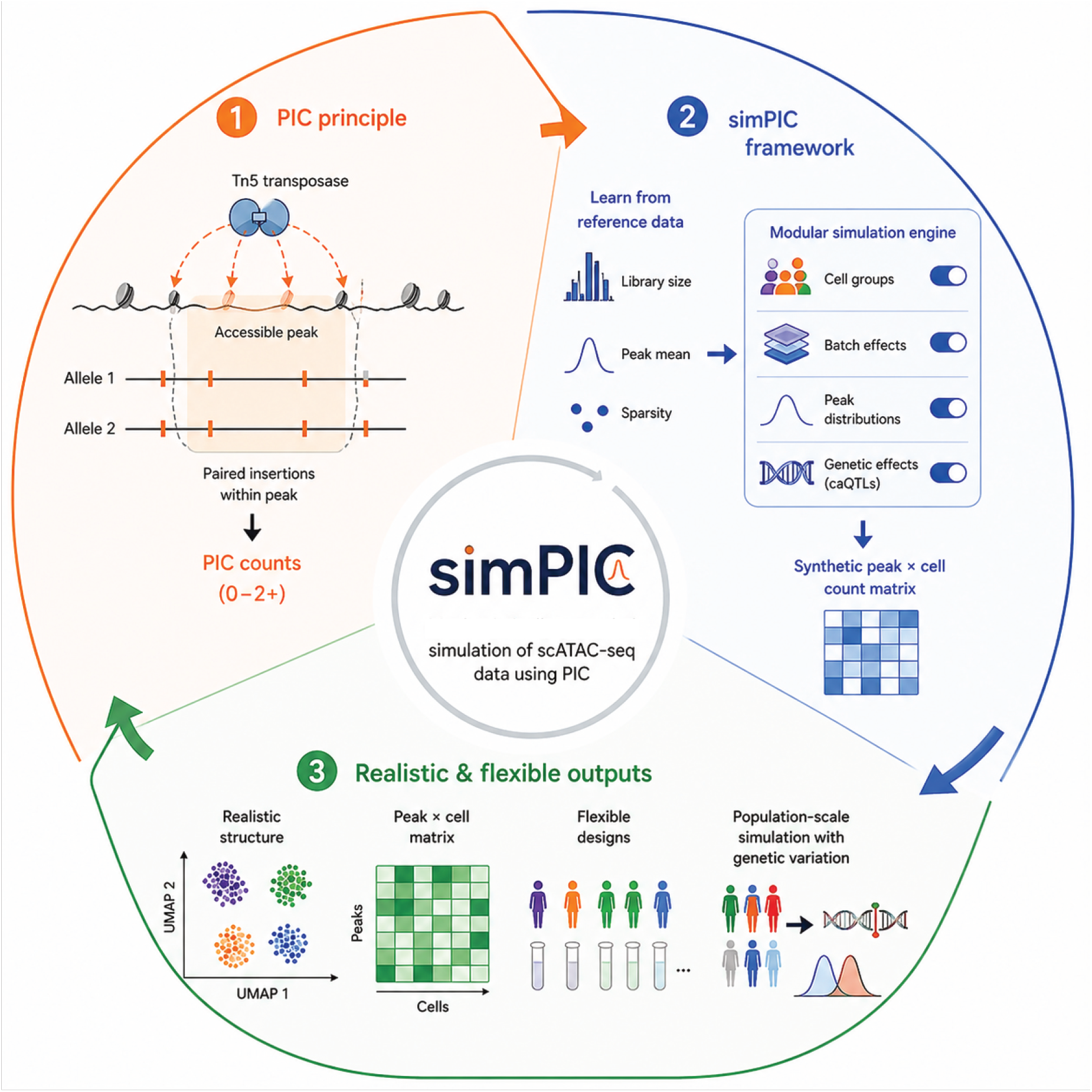
Schematic representation of simPIC. simPIC simulates scATAC-seq data using paired insertion counts (PIC), which quantify Tn5 insertion events within accessible peaks. The framework learns key properties from reference data, including library size, peak accessibility, and sparsity, and uses a modular simulation engine to generate synthetic peak-by-cell count matrices with configurable cell groups, batch effects, peak distributions, and genetic effects. The resulting datasets preserve realistic accessibility structure while supporting flexible single-sample and population-scale simulations.

## 2 Results

### 2.1 The simPIC model framework

simPIC is a modular R/Bioconductor package for simulating synthetic scATAC-seq datasets that capture the key biological and technical features observed in real data. The simPIC framework supports two main simulation modes: (a) simulating one or more cell types within a single sample, optionally with multiple experimental batches; and (b) simulating population-scale data with inter-individual genetic variation via integration with splatPop [6] (Figure 2, Supplementary Table S1). All output is returned as a SingleCellExperiment object, enabling easy integration with downstream tools. The simPICcompare() function, provides diagnostic plots comparing real and simulated data across key metrics: library size, peak accessibility, and sparsity, enabling rapid model calibration. Together, these features support user-defined scATAC-seq simulations for method evaluation and hypothesis testing under explicit generative assumptions.

**Fig. 2:**
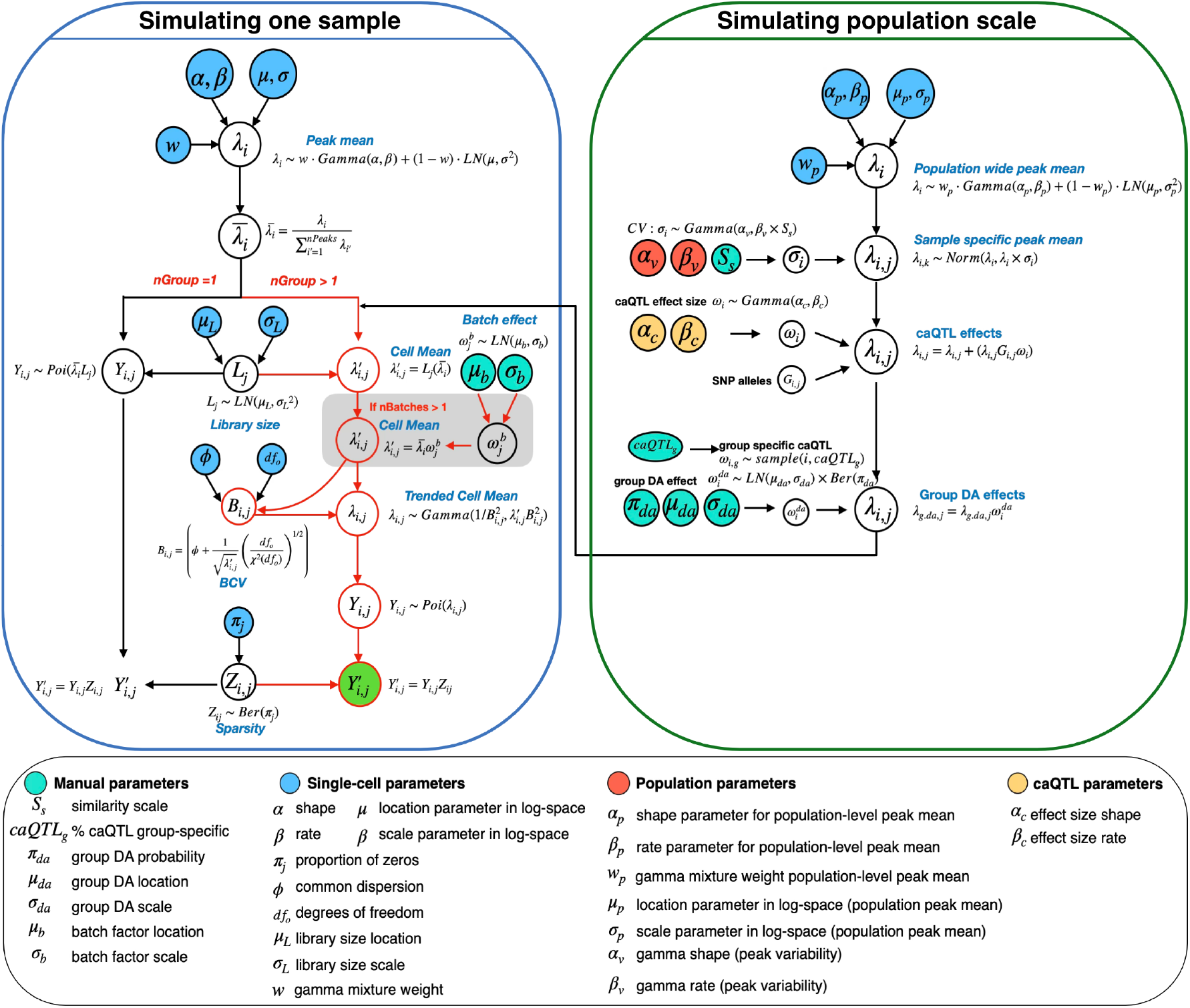
Graphical model representation of the simPIC simulation framework. Input parameters estimated from real data are indicated in blue, population-level parameters are in orange, manually specified parameters are in cyan. caQTL parameters are in yellow. The green circle represents the final output, the simulated peak-by-cell count matrix. Population-scale simulation is achieved through integration with splat-Pop. Detailed list of parameters is in Supplementary Table S1. Abbreviations : (*LN*)-lognormal,(*Ber*)-Bernoulli, (*Poi*)-Poisson distribution.

simPIC offers four peak-mean models—gamma, Weibull, lognormal-gamma (ln-gamma), and Pareto; which differ in the shape of the marginal peak accessibility distribution they impose. Across a benchmark of 25 cell types from six datasets (Table 1) spanning multiple scATAC-seq technologies and species, simPIC with ln-gamma peak means consistently achieved the best distributional accuracy and is therefore the recommended default. All four variants complete simulations in under five minutes for the cell types evaluated here; runtime differences between variants are small relative to differences between simPIC and competing tools (Section 2.5).

**Table 1:** Datasets used in this study.

| Dataset:Year | Organism | Platform | Sequencing | Accession No |
| --- | --- | --- | --- | --- |
| PBMC5k:2021 | *Homo sapiens* | 10x<br>Chromium-<br>(snATAC) | Illumina<br>Novaseq 6000 | NA |
| PBMC10k:2022 | *Homo sapiens* | 10x<br>Chromium-<br>(snATAC) | Illumina<br>Novaseq 6000 | NA |
| Sathpathy:2019 | *Homo sapiens* | 10x<br>Chromium-<br>(scATAC) | Illumina HiSeq<br>2000 | GEO:<br>GSE129785 |
| Buquicchio:2022 | *Mus musculus* | 10x<br>Chromium-<br>(scATAC) | Illumina HiSeq<br>4000 | GEO:<br>GSE199799 |
| Cusanovich:2018 | *Mus musculus* | sciATAC | Illumina HiSeq<br>2500 | GEO:<br>GSE111586 |
| Fly Brain:2022 | *Drosophila*<br>*melanogaster* | scATAC | NextSeq500<br>and<br>NovaSeq6000 | GEO:<br>GSE163697 |

To contextualise simPIC within the broader simulation landscape, we compared it with four tools capable of producing count-level scATAC-seq data: simCAS [10], scDesign3 [12], scMultiSim [14], and DiTSim [15]. These tools differ substantially in their required inputs, modelling frameworks, and computational demands (Supplementary Tables S2-S3). Briefly, scDesign3 is a multi-modal simulator using Generalized Additive Models for Location, Scale, and Shape (GAMLSS) and copula models that requires cell-type annotations; scMultiSim is a mechanistic simulator guided by GRN and cell-cell interactions, targeting RNA and ATAC jointly; and DiTSim is a diffusiontransformer-based deep learning simulator requiring cell-type labels and GPU access. Unlike these tools, simPIC requires only a count matrix as input, is the only simulator in this comparison that supports population-scale and caQTL simulation, and provides built-in diagnostic functions.

### 2.2. simPIC accurately reproduces single-cell accessibility profiles across cell types

We evaluated simPIC’s ability to reproduce distributional properties of scATAC-seq data across 25 cell types from six publicly available datasets: PBMC5k, PBMC10k, Satpathy et al. [18], Buquicchio et al. (liver) [19], Cusanovich et al. [20], and Janessens et al. (Fly brain) [21]. These datasets span multiple scATAC-seq technologies (10x Chromium, sci-ATAC-seq, bulk-indexed) and three species (human, mouse and *Drosophila melanogaster*), providing a broad evaluation regime (Table 1). For each cell type, simulations matched the number of cells in the corresponding real dataset (Supplementary Table S4). All four simPIC variants were compared alongside scDesign3, simCAS, DiTSim, and scMultiSim, on three core distributional properties: log library size, log peak mean, and cell sparsity. For each statistic we report Kolmogorov– Smirnov (KS) distances between simulated and real distributions, with lower values indicating closer agreement.

We show a representative cell type from Cusanovich dataset: Kidney Podocytes. All four simPIC variants closely reproduced the log library size distribution of real data (KS = 0.06–0.09), while scDesign3 showed moderate deviation (KS = 0.38), simCAS performed similarly (KS = 0.04), and DiTSim deviated more substantially (KS = 0.39). scMultiSim showed the largest discrepancy (KS = 1.00), reflecting a systematic mismatch in both scale and shape (Figure 3a). A similar pattern was observed for log peak mean: simPIC variants maintained low KS distances (0.08–0.18), with simPIC default ln-gamma being lowest (KS = 0.08), comparable to scDesign3 (KS = 0.09) and DiTSim (KS = 0.09), while simCAS deviated more substantially (KS = 0.38), and scMultiSim again showed maximal divergence (KS = 1.00) (Figure 3b). For cell sparsity, a defining feature of scATAC-seq data, simPIC consistently achieved the lowest KS values (0.05–0.08), with simCAS performing comparably (KS = 0.05) and scDesign3 showing moderate deviation (KS = 0.37). In contrast, DiTSim and scMultiSim both showed KS = 1.00, indicating a failure to capture the characteristic high-sparsity structure of real data (Figure 3c).

**Fig. 3:**
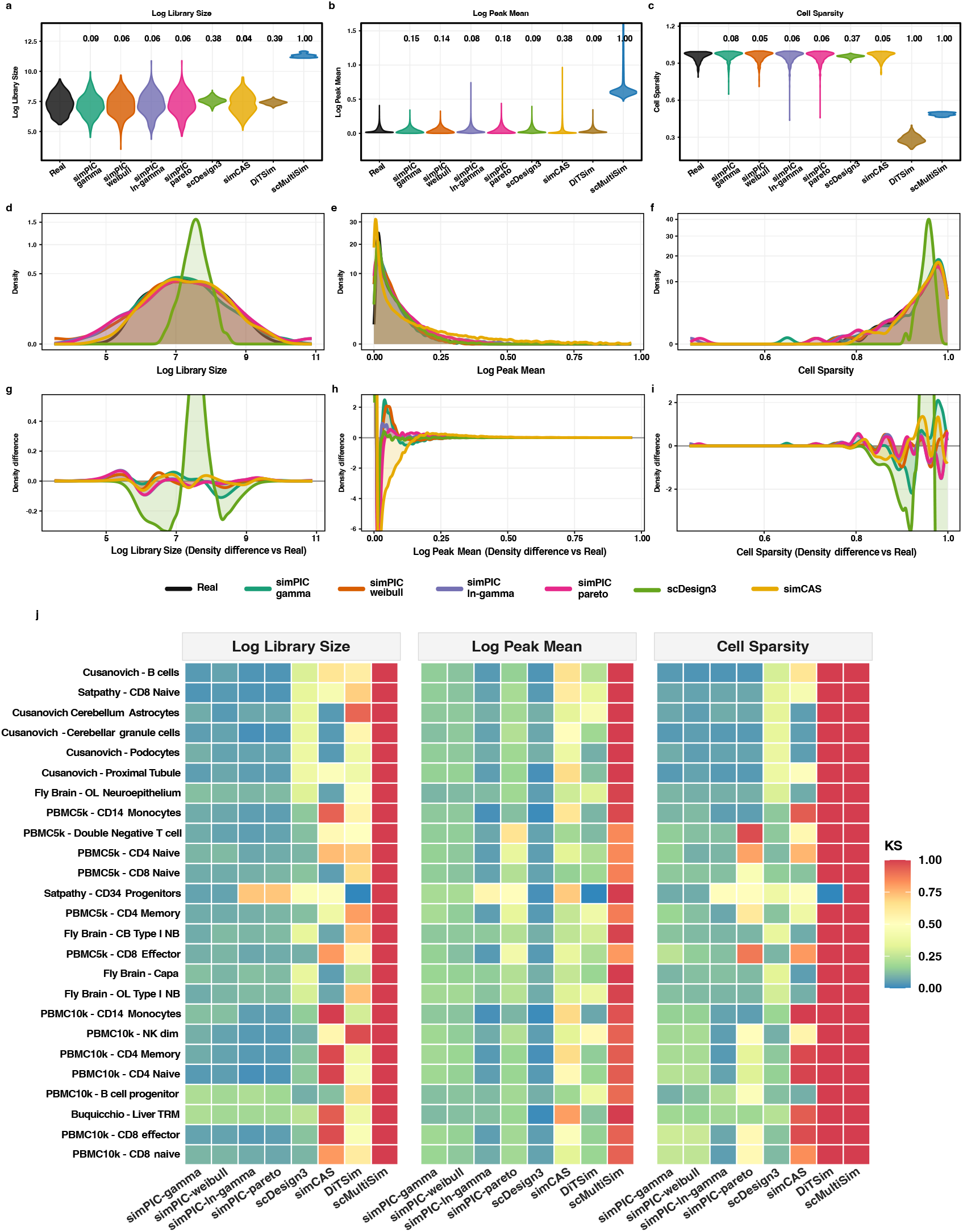
Distributional comparison of simPIC and competing simulators across 25 cell types from six datasets spanning multiple technologies and species. (a-c) Violin plots of (a) log library size, (b) log peak mean, and (c) cell sparsity for Cusanovich: Kidney Podocyte cell type between each simulator and real data. Annotated values are median Kolmogorov–Smirnov (KS) statistics per tool; lower values indicate closer distributional agreement with real data. (d-f) Density plots for the same cell type, showing (d) log library size, (e) log peak mean, and (f) cell sparsity for real data (black) and each simulator. (g-i) Density difference plots (simulated minus real) for the same cell type; values close to zero indicate distributional agreement. (j) Heatmap of KS statistics across all 25 cell types and three properties; blue indicates close agreement with real data.

Density-based comparisons reinforced these findings.DiTSim and scMultiSim are not shown in the density plots because they produced degenerate or near-degenerate discrepancies for key metrics in this benchmark setting: scMultiSim had a KS distance of 1 for peak means, and DiTSim had a KS distance of 1 for cell sparsity. We therefore omitted these two methods from the density and density-difference panels to improve visual resolution among the remaining competing methods. For log library size, simPIC variants broadly overlapped the real-data distribution, whereas scDesign3 produced a narrower distribution centred near the middle of the observed range. The corresponding density-difference plot showed a positive deviation near the distribution centre and negative deviations in the shoulders, consistent with under-representation of library-size heterogeneity (Figure 3d and 3g). For log peak mean, simPIC closely followed the real density across most of the distribution, with modest deviations near zero; simCAS showed a stronger negative deviation at the lowest peak means and a broader positive deviation at intermediate values (Figure 3e and 3h). We also note that simPIC simulates peaks conditionally independently given peak-wise and cellwise parameters, it does not explicitly model peak–peak covariance (co-accessibility). Consistent with this design choice, we observe differences in the tails of the peak–peak correlation distribution between simulated and real data (Supplementary Figure S2 and Supplementary Table S5). For cell sparsity, simPIC reproduced the high-sparsity mode characteristic of real scATAC-seq data, with residual discrepancies concentrated in the extreme high-sparsity tail; scDesign3 produced a sharper and shifted sparsity profile, while simCAS showed larger departures from the real distribution (Figure 3f and 3i). Beyond these three properties, we examined peak sparsity, log peak variance, the mean-variance relationship, and the associations of peak mean with non-zero proportion, and peak sparsity for this cell type (Supplementary Figure S3 (page 7-8)). These additional diagnostics further support the distributional agreement between simPIC-simulated and real data.

To assess whether these patterns generalised across datasets, we computed KS statistics between each simulated and real data across all 25 cell types. Consistent with the above summaries, the per-cell-type KS heatmap showed that all simPIC variants had uniformly low KS values for log library size, across cell types, reflecting stable recovery of library size distributions. In contrast, scDesign3 and simCAS exhibited increased variability, with several cell types showing moderate deviations, while DiTSim and scMultiSim consistently showed high KS values, indicating poor agreement with real data. For log peak mean, simPIC again maintained low and consistent KS values across nearly all cell types, demonstrating reliable modelling of peak-level accessibility. scDesign3 showed competitive performance for some cell types but with greater variability overall. simCAS and DiTSim showed higher KS values across multiple cell types, while scMultiSim consistently failed to recover the peak mean distribution, as reflected by uniformly high KS values. For cell sparsity, simPIC variants closely matched real data across the majority of cell types, with low KS values and minimal variability. scDesign3 showed moderate but consistent deviations (KS = 0.37), whereas simCAS exhibited cell-type-dependent performance with both moderate and large deviations. DiTSim and scMultiSim consistently showed maximal KS values (KS = 1), indicating a systematic failure to capture the sparsity structure across all cell types (Figure 3j). The full per-cell-type evaluation across all 25 cell types, including both the three core properties and the additional peak-level diagnostics are provided in Supplementary Figure S3 (page 9-56).

Collectively, these results demonstrate that simPIC provides the most consistent and robust reconstruction of key scATAC-seq data characteristics across diverse biological settings, than the available tools, with the ln-gamma variant performing comparably to or better than alternatives.

### 2.3. simPIC outperforms competing simulators across quantitative metrics

We quantified simulator performance by computing composite ranks from four distributional similarity measures: Mean Absolute Deviation (MAD), Mean Absolute Error (MAE), Root Mean Square Error (RMSE), and 1 − Pearson Correlation Coefficient (PCC). Rankings were computed per cell type and aggregated to obtain an overall composite rank (lower is better). A Friedman test, treating cell type as a blocking factor confirmed highly significant differences in performance across simulators (Figure 4a). Across 25 cell types, all four simPIC variants consistently achieved lower (better) composite ranks than DiTSim and scMultiSim (Wilcoxon signed-rank tests, all FDR *<* 0.001; Figure 4b), demonstrating a clear and consistent performance advantage. Among simPIC variants, simPIC(ln-gamma) achieved the lowest median overall composite rank, and ranked first across all six datasets (Supplementary Figure S4), supporting its use as default. Compared to sim-CAS, simPIC variants generally achieved lower overall composite ranks across cell types, although statistical significance after FDR correction was observed only for the ln-gamma variant (FDR = 0.0016). This pattern suggests that while simCAS reproduces several individual data characteristics well, simPIC provides more consistently strong performance across benchmark metrics. Comparisons with scDesign3 were significant for simPIC(gamma) (FDR = 0.011) and directionally consistent for the remaining simPIC variants (FDR = 0.058–0.360), supporting that simPIC achieves balanced performance across benchmark metrics (Figure 4b). Collectively, these results show that simPIC consistently ranked among the best-performing simulators across cell types and evaluation metrics, with the ln-gamma parameterisation yielding the strongest overall performance.

**Fig. 4:**
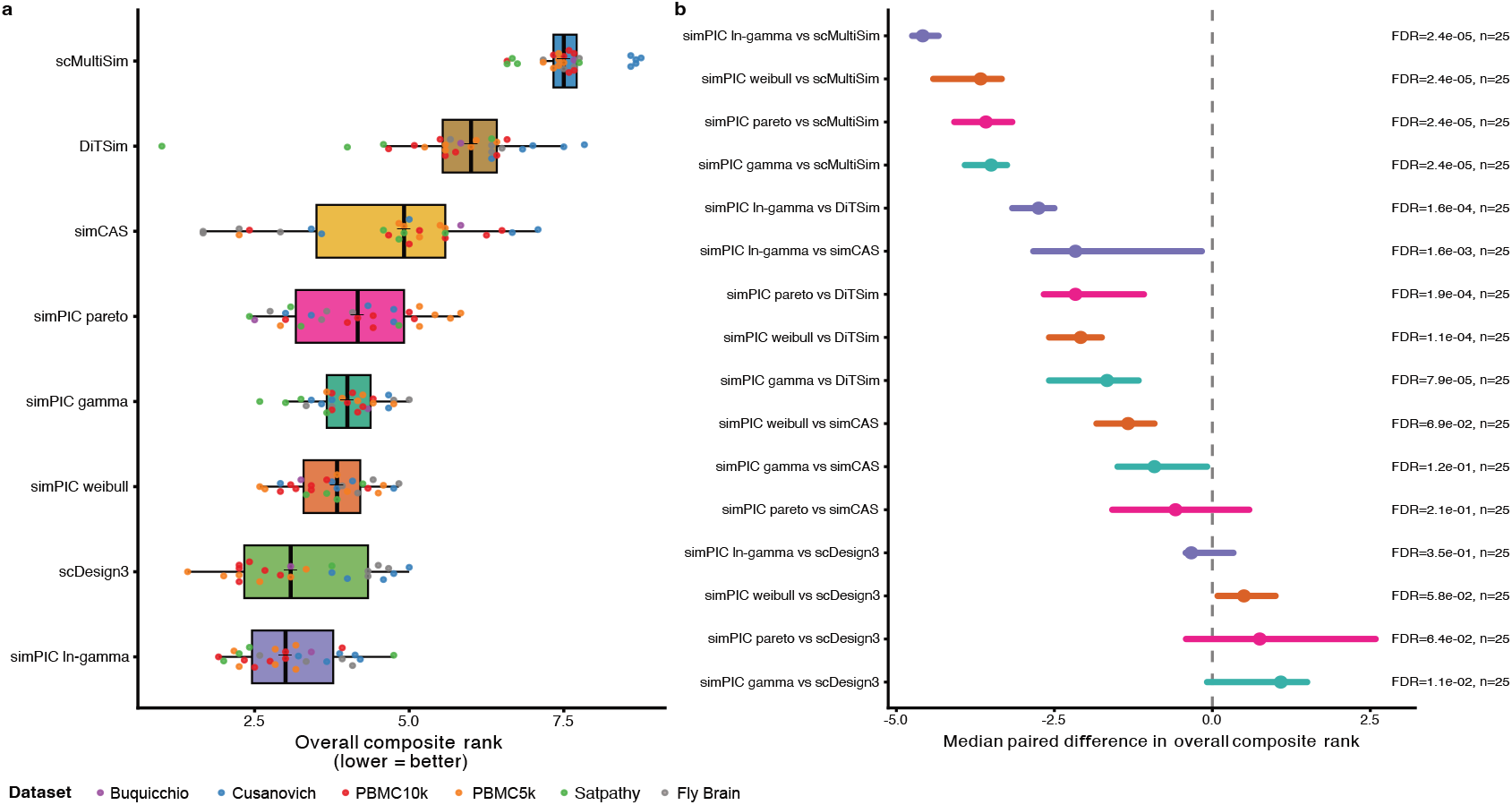
Quantitative benchmarking of simulators across 25 cell types from six datasets. Composite ranks were derived from MAD, MAE, RMSE, and 1 − PCC for library size, peak mean, and cell sparsity; lower rank indicates better performance. (a) Overall composite rank distributions per simulator across all 25 cell types. (b) Median paired difference in overall composite rank between each simPIC variant and each competing method; negative values indicate simPIC outperforms the comparator. BH-corrected FDR values from Wilcoxon signed-rank tests are shown; *n* denotes the number of cell types.

### 2.4. simPIC simulates multiple cell types and batch effects

We next evaluated whether simPIC could model two key sources of heterogeneity in scATAC-seq data: cell-type structure, represented as user-defined groups, and technical batch effects. simPIC enables the user to specify the number of groups, their proportions, and the extent of differential accessibility between them (Supplementary Table S1). Batch effects are modelled as peak-wise multiplicative offsets on peak accessibility, capturing technical variation orthogonal to biological structure.

#### Simulating multiple cell types with differential accessibility

We evaluated group simulation using subsets of PBMC5k, PBMC10k and Fly Brain data [19]. In the two-group setting, we used nGroups = 2 and group.probs = c(0.5, 0.5), the simulated matrix captured distinct group-specific accessibility profiles, with two well-separated clusters in principal component analysis (PCA) space consistent with the specified 1:1 group ratio (Supplementary Figure S5 a-c). With three groups and imbalanced sizes (nGroups = 3, group.probs = c(0.5, 0.25, 0.25)), three distinct clusters were observed with group sizes reflecting the specified proportions (Supplementary Figure S5 a-c), demonstrating that simPIC reproduces both biological heterogeneity and population structure, supporting its use in simulating realistic multi-cell-type scenarios.

#### Simulating batch effects across cell groups

Batch effects were simulated using peak-wise, batch-specific multiplicative offsets to expected peak accessibility (peak means) values. In a two-batch, two-group design (nGroups = 2, group.probs = c(0,5,0.5), nBatches = 2), we generated two batches with equal distribution of cells from each group, specifying batch-level offsets while holding biological group structure constant. The simulated data showed separation in lower-dimensional space by both biological group and batch identity (Supplementary Figure S6a). This batch associated structure was further quantified by K-Nearest neighbours (KNN) batch-mixing within each group (*k* = 50), yielded a median same-batch neighbour fraction of 0.78 (IQR 0.24) in group 1 and 0.76 (IQR 0.20) in group (Supplementary Figure S6b), confirming coherent batch-associated structure. A peak detection rate shift plot (Batch 1 versus Batch 2) showed systematic feature-level deviations from the diagonal within each group, consistent with the peak-wise multiplicative batch model (Supplementary Figure S6c).

### 2.5. simPIC is computationally efficient and scalable

We benchmarked simPIC’s computational performance by measuring runtime and peak RAM for all simulators across 25 cell types from the six datasets, spanning 203–14,633 cells and 36,496–175,151 peaks per cell type (Supplementary Table S4).

simPIC simulated all 25 cell types in under five minutes, with median runtime of 6–20 seconds depending on variant (Figure 5a). simCAS runtime ranged from 14 seconds to 18 minutes. DiTSim required between 12 minutes and 8.9 hours, with the largest cell type (Satpathy CD34 Progenitors: 14,633 cells, 175,151 peaks) requiring approximately 8.9 hours. scMultiSim had a median runtime of 4.7 minutes but required approximately 48.6 hours for Satpathy CD34 Progenitors. scDesign3 was the most computationally demanding, runtime ranging from 1.6 to 103.9 hours across cell types. For the PBMC10k CD14 Monocytes cell type (2,201 cells, 106,967 peaks), scDesign3 required approximately 103.9 hours and 576 GB of RAM, compared with 37-44 seconds and approximately 9.6 GB for simPIC variants.

**Fig. 5:**
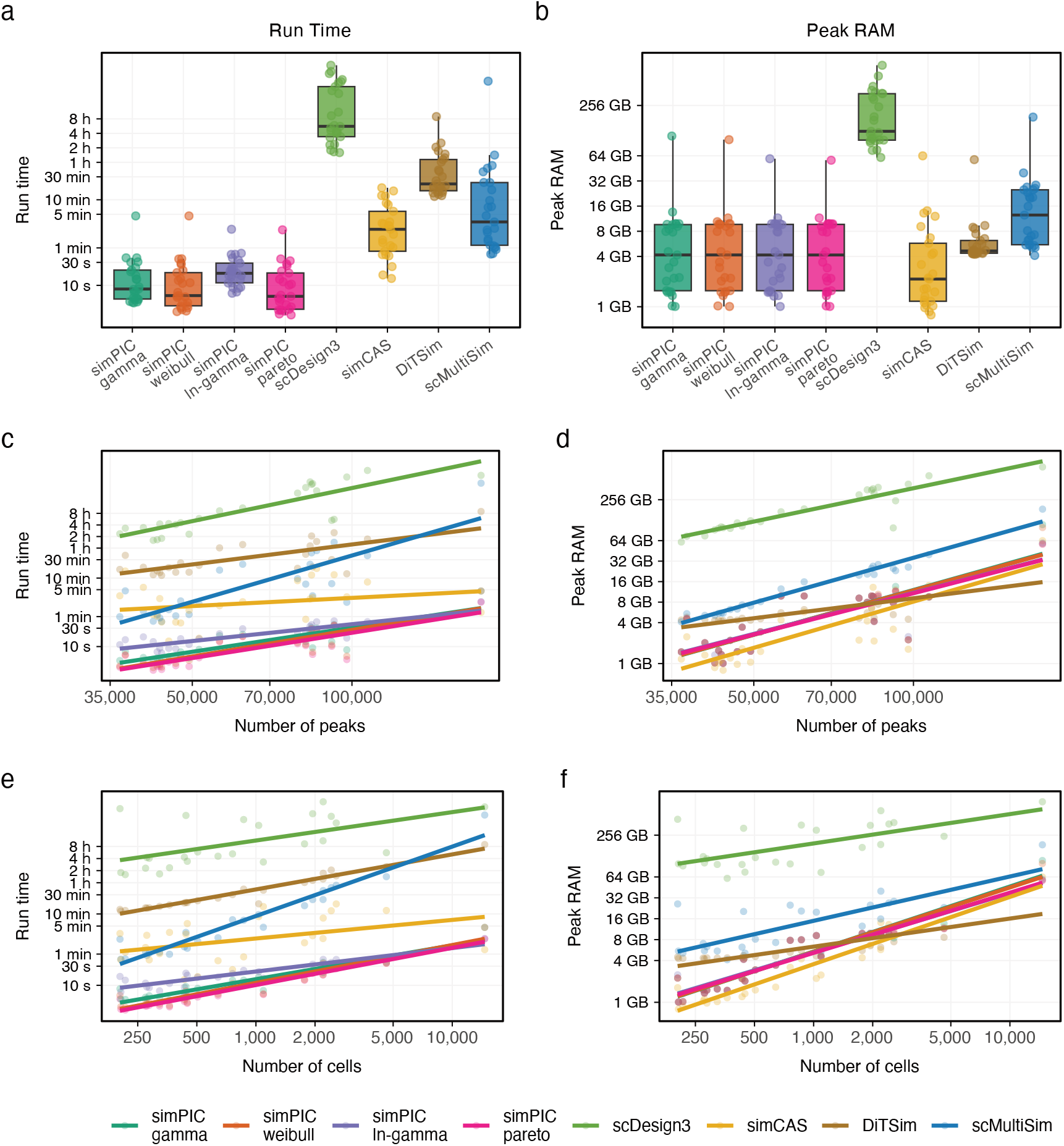
Computational performance across five simulators. (a) Runtime and (b) Peak RAM for all simulators across 25 cell types from six datasets. Each point represents one cell type; boxes summarise the distribution across cell types. (c-d) Runtime and peak RAM as a function of number of peaks. (e-f) Runtime and peak RAM as a function of number of cells.

Peak RAM followed analogous patterns (Figure 5b). simPIC variants used 1.0–110 GB; simCAS used 0.8–63.9 GB; DiTSim used 4.2–57.6 GB; scMultiSim used 4.1–185.9 GB; and scDesign3 used 61.1–774.7 GB. The memory requirements of scDesign3 reflect the scaling behaviour of its copula-based model with increasing feature dimensionality, rendering it infeasible for large cell types on standard computing infrastructure. Among simPIC variants, simPIC(ln-gamma) had a median runtime of approximately 20 seconds— modestly slower than simPIC(Weibull) (7 seconds) and simPIC(Pareto) (6 seconds) but completed all 25 cell types well within practical limits. Given its superior distributional accuracy (Figure 3), the runtime difference between simPIC variants is negligible in practice, and simPIC(ln-gamma) remains the recommended default. Runtime and memory usage scaled smoothly with the number of cells and peaks for all simPIC variants (Figure 5c-f), confirming predictable performance across a range of dataset sizes.

### 2.6. Simulating population scale data with genetic effects

We evaluated simPIC’s ability to simulate biologically realistic inter-individual variation in chromatin accessibility, we generated population-scale scATAC-seq data incorporating genetic effects. We compared these simulations with a publicly available Alzheimer’s disease dataset [22], at the time of writing the only open-access scATAC-seq resource with both genotype and cell-type-resolved accessibility data across individuals. Cell-level parameters were estimated from the individual with the most high-quality cells and population-level parameters were derived from pseudobulk aggregated profiles, following an approach analogous to splatPop [6]. Using these estimates, we simulated accessibility counts for one cell type across multiple individuals and compared the structure of real and simulated data in lower-dimensional PCA space.

In the microglia population (library 5; 344 cells, three individuals, 1,693 peaks), both real and simulated data showed comparable global structure, with approximately 2% of variance explained by the top two principal components (Figure 6a). Mean silhouette widths donor labels were close to zero in both datasets (real: 0.005; simulated: 0.017), consistent with substantial donor overlap (Figure 6b). Mean neighbourhood purity was higher in the real data (0.539) than in the simulation (0.473) (Figure 6c), indicating stronger local donor consistency than simPIC recovers in this library. Comparable patterns were observed across additional earlyAD and nonAD libraries. Across libraries, mean silhouette widths were generally close to zero in both real and simulated data, whereas neighborhood purity was often higher in the real data for nonAD libraries with larger cell counts than the earlyAD libraries (244–604) (e.g., Library7 (ncells = 1012): real 0.284, simulated 0.197; Library10 (ncells = 2057): real 0.257, simulated 0.191), which may partly reflect greater stability of kNN-based estimates at higher cell numbers (Supplementary Figures S7-S13, Supplementary Table S5).

**Fig. 6:**
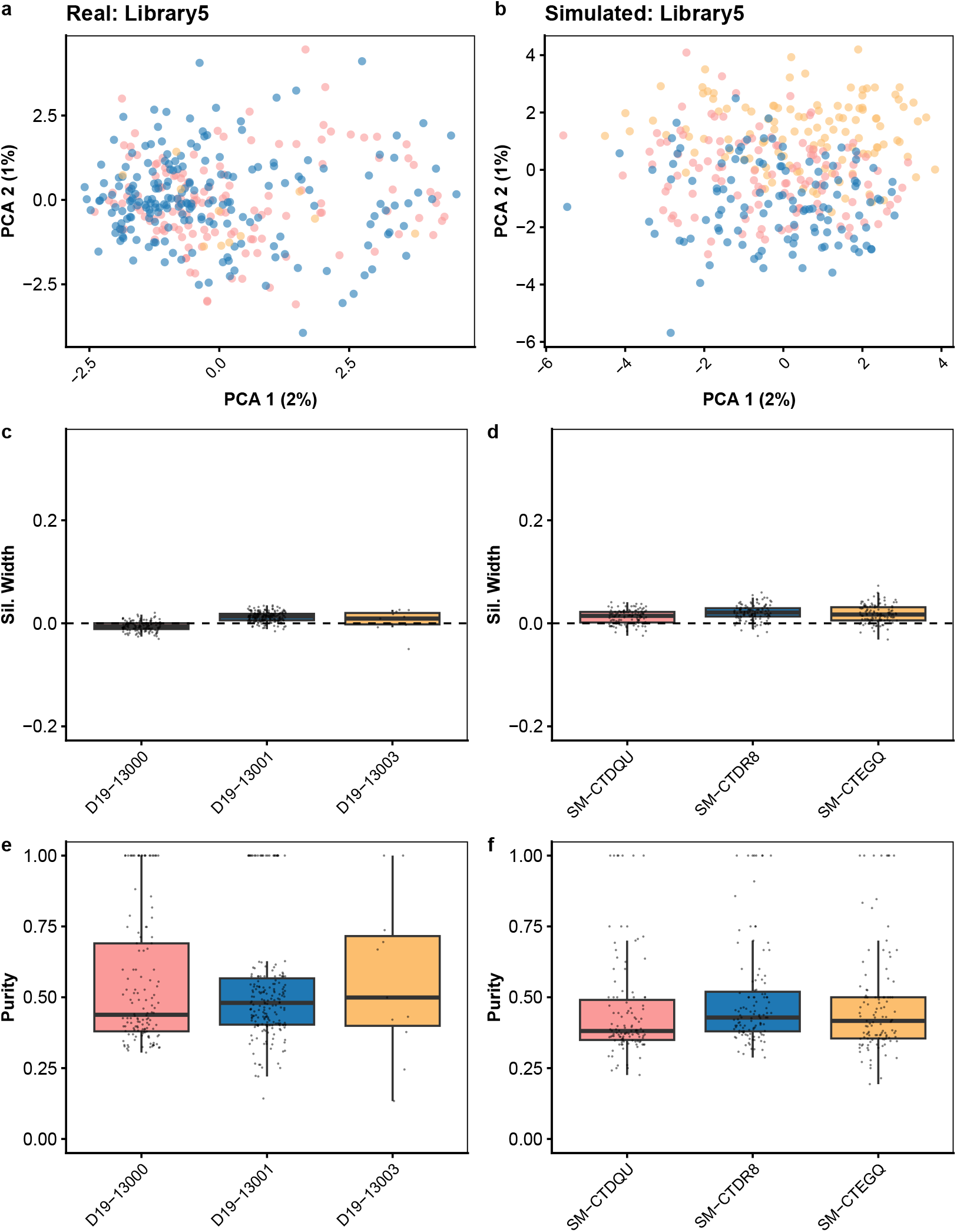
Quantifying clustering in real and simPIC-simulated data with inter-individual genetic variation. (a-b) PCA plot of real and simulated data, coloured by individual, showing the preservation of individual specific structure. (c-d) Silhouette width comparison between real and simulated data, assessing cluster compactness. (e-f) Neighbourhood purity analysis, evaluating local structure consistency across real and simulated datasets.

We assessed simPIC’s modelling of genetic effects by performing cis-caQTL mapping with LIMIX [23] on both real and simulated microglia data for chromosome 22, testing all SNP–peak pairs within a ± 100 kb window using a linear mixed model that accounts for relatedness and technical covariates. We identified 14 significant associations in the real data and 46 in the simulated data at an empirical *p*-value threshold of *<* 0.005 (Figure 7a-b). While no individual SNPs overlapped between datasets, expected given that simPIC samples genotypes probabilistically without using real haplotypes or linkage disequilibrium (LD) structure, the overall number of discoveries was similar, suggesting comparable statistical power. Significant SNPs formed spatially clustered patterns within 400 kb genomic windows in both datasets (Figure 7c-d), consistent with LD-driven regional aggregation; demonstrating that simPIC preserves local enrichment patterns driven by linkage disequilibrium typical of regulatory hotspots, even without reproducing exact loci. Effect-size distributions across significant SNP-peak pairs were nearly identical in real and simulated data (Figure 7e), suggesting that simPIC accurately reproduces the genome-wide spectrum of genetic effect magnitudes.

**Fig. 7:**
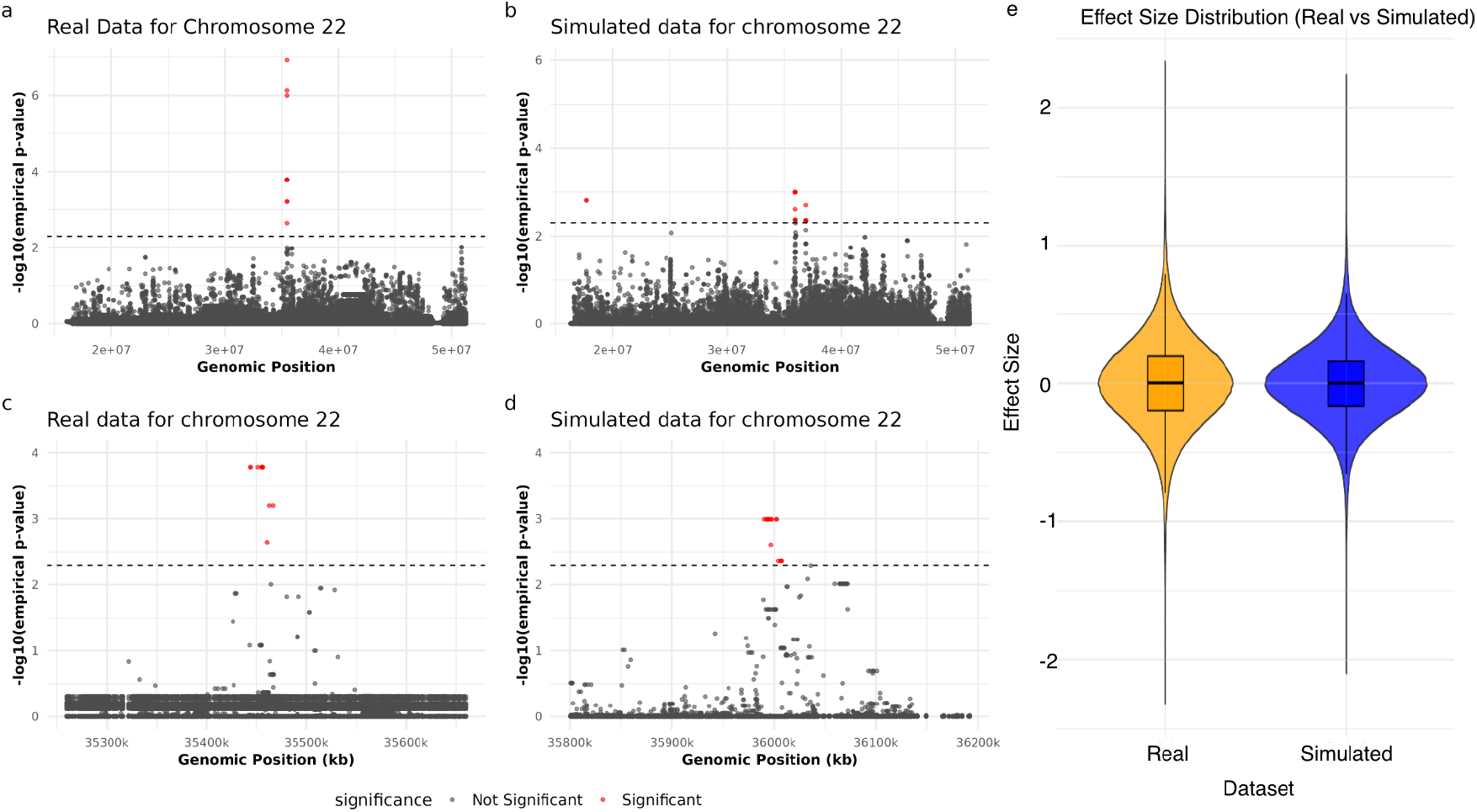
cis-caQTL mapping results for chromosome 22 in real and simPIC-simulated microglia data. (a-b) Manhattan plots of − log_10_(empirical *p*-value) for SNP-peak associations in (a) real and (b) simulated data; horizontal dashed lines indicate the significance threshold (*p <* 0.005). (c-d) Zoomed-in views of 400kb regions showing spatial clustering of significant associations in each dataset. (e) Effect-size distributions across significant SNP-peak pairs in real and simulated data.

Together, these results demonstrate that simPIC can simulate population-scale scATAC-seq data with genetic effects that closely mirrors real data in structure, variability, and association statistics, making it a valuable tool for benchmarking caQTL discovery pipelines and modelling regulatory variation across individuals.

### 2.7. simPIC-simulated data preserves differential accessibility method rankings

A key use case for simulation tools is benchmarking computational methods under controlled ground truth. To assess whether simPIC-simulated data produces DA method rankings consistent with those reported using real data, we benchmarked ten DA detection methods via the Libra framework [24] across 25 cell types: six pseudobulk approaches (edgeR log-likelihood ratio test (LRT), edgeR quasi-likelihood F-test (QLF), DESeq2 Wald, DESeq2 LRT, limma-trend, limma-voom) and four single-cell approaches (Wilcoxon, *t*-test, negative binomial, logistic regression). Simulations were run at two effect sizesda.facLoc = 0.1 and 0.5, with ground-truth differential peaks defined by the simulation parameters (Supplementary Figure S14a).

At moderate effect size (da.facLoc = 0.5), pseudobulk methods collectively outperformed single-cell methods, with logistic regression and negative binomial achieving the highest AUC-ROC among all ten methods (0.707; Supplementary Figure S14a). This ordering is consistent with rankings reported by Teo et al. [24] using real scATAC-seq data with orthogonal validation, supporting the utility of simPIC for DA method benchmarking. AUC-PR values, which account for class imbalance, ranged from 0.305 to 0.337 against a random baseline of 0.131 (Supplementary Figure S14a-b). At low effect size da.facLoc = 0.1), sensitivity was reduced across all methods, with the relative ordering of pseudobulk versus single-cell approaches maintained (Supplementary Figure S14c-d). We note that this validation is specific to the DA task and the generative assumptions of simPIC; the degree to which simulation-based rankings reflect real-data rankings will depend on how closely the simulator’s generative model matches the true data-generating process for the task of interest.

## 3. Discussion

The shift of single-cell technologies such as scATAC-seq to population-scale studies has created unprecedented opportunities for dissecting the genetic regulation of chromatin accessibility, but has also introduced new analytical challenges. Simulation tools are essential for method development because they enable controlled data generation under defined assumptions and study designs. The R/Bioconductor package simPIC provides a flexible and interpretable framework that captures key distributional properties of scATAC-seq data. Across 25 cell types from six datasets, simPIC with ln-gamma peak means achieved the highest overall fidelity to empirical distributions while remaining computationally tractable, whereas alternative methods required hours to days or became infeasible at scale. By estimating parameters directly from empirical data and allowing users to control group structure, batch, and genetic effects, simPIC balances realism with usability. Through integration with splatPop [6], simPIC further supports population-scale simulations with explicit genetic effects, a capability not supported by existing scATAC-seq simulators.

Empirical benchmarking strategies based on orthogonal information, such as motif enrichment, transcription factor binding from ChIP-seq, or concordance with paired scRNA-seq, are valuable for validating specific aspects of scATAC-seq analyses but typically provide indirect proxies rather than explicit locus-by-cell ground truth. Simulation addresses this limitation by generating datasets with known structure. simPIC generates peak-by-cell count matrices with explicitly defined cell-group labels, effect sizes, batch structure, and genetic effects, enabling controlled evaluation of analytical methods. These capabilities are particularly important for population-scale analyses where ground truth is not directly observable in empirical datasets and experimental designs and confounding factors are difficult to disentangle. Consistent with this rationale, differential accessibility method rankings derived from simPIC-simulated data across 25 cell types were concordant with those reported in real-data benchmarks [24] using real scATAC-seq data with orthogonal validation. These results provide supporting evidence that simPIC can serve as a useful tool for method benchmarking beyond the simulation setting. We note that this concordance is task-specific and depends on the fidelity of the generative model; simulation-based rankings should not be expected to transfer universally to all real datasets.

A defining strength of simPIC is its ability to model genetic effects at population scale, making it uniquely suited for benchmarking caQTL mapping approaches. By simulating ground-truth causal variants, effect-size distributions, and realistic association patterns, simPIC enables systematic evaluation of association methods under controlled conditions. Researchers can use simPIC to assess statistical power across cohort sizes, test caQTL mapping accuracy in high-LD regions, and evaluate the reproducibility of QTL detection in the presence of batch effects and cell-typespecific heterogeneity. This functionality addresses a major limitation for caQTL mapping method development, namely the limited availability of real population-scale scATAC-seq datasets, which remain costly and scarce.

Beyond genetic analyses, simPIC enables controlled assessment of pre-processing and downstream workflows. Users can vary sparsity, sequencing depth, and batch effects to evaluate normalisation and batch-correction methods under known conditions. Similarly, user-defined group structure provides ground truth for clustering, annotation, and feature selection, ensuring that observed performance reflects recovery of biological signal rather than artefactual structure. These properties also support emerging applications, including benchmarking of variant-effect prediction and integrative multi-omic analyses, where empirical ground truth remains limited. simPIC therefore complements, but does not replace, empirical validation.

Like all simulators, simPIC has limitations. simPIC models peaks as conditionally independent given peak- and cell-level parameters, and therefore does not explicitly model peak–peak covariance (co-accessibility). Although simPIC recovers the central mass of the correlation distribution, it captures tail behaviour inconsistently across cell types, reflecting this modelling assumption. We therefore report peak-peak correlation diagnostics to quantify the extent to which this modelling choice impacts co-accessibility patterns (Supplementary Figure S2, Supplementary Table S5).

More broadly, because ground truth is generally unavailable in empirical scATAC-seq datasets, there is no definitive external reference for establishing a “true” method ranking across real applications. Accordingly, we evaluate simPIC by: (i) assessing fidelity to empirically observable properties (e.g., library size, peak means, and sparsity), and (ii) enabling controlled benchmarking under explicit generative assumptions where ground truth is known. A large-scale comparison of method rankings across multiple empirical datasets and analysis tasks is beyond the scope of this manuscript; our goal here is to introduce simPIC and demonstrate that it reproduces key measurable properties while enabling ground-truth benchmarking under controllable settings. Exact positional concordance of significant SNPs with those observed in empirical datasets is unlikely, reflecting simplified assumptions about recombination and effectsize distributions. Model complexity also imposes practical constraints on alternative approaches. For example, scDesign3 achieved competitive performance for some peaklevel metrics but less effectively captured cell-level metrics such as library size and cell sparsity, while also requiring substantial memory and extended runtimes, limiting its applicability at scale. In contrast, simPIC maintained consistent performance across datasets with modest computational requirements. Nonetheless, simPIC successfully captures higher-order structural properties that represent core features of complex trait architecture. Current gaps include the lack of conditional simulations across disease states, time points, and the absence of explicit modelling of epigenetic modifications or chromatin interactions. Future extensions that incorporate these aspects, as well as multi-omic layers linking accessibility to gene expression or proteomics will further expand its scope and impact.

In summary, simPIC provides an interpretable, flexible, computationally efficient and reproducible framework for simulating scATAC-seq data with explicit control over experimental design and genetic effects. By enabling benchmarking under known ground truth while maintaining fidelity to empirical data, simPIC provides a valuable foundation for advancing reproducible and scalable single-cell epigenomics research.

## 4 Conclusion

simPIC provides and interpretable, computationally efficient and extensible R/Bioconductor framework for simulating single-cell chromatin accessibility data across diverse experimental settings. Released under a GPL-3 license, simPIC supports single-sample and population-scale simulations, including configurable cell groups, batch effects, and genotype-dependent effects on chromatin accessibility. Across 25 cell types from six datasets, simPIC with ln-gamma peak means achieved the highest overall distributional fidelity while requiring substantially less computation than deep-learning and copula-based alternatives. To the best of our knowledge, only simPIC explicitly supports population-scale simulation with genetic effects, enabling controlled benchmarking of caQTL mapping when empirical ground truth is unavailable. Together, its modular design, interpretable parameterisation, and diagnostic framework make simPIC a scalable platform for benchmarking, method development, and experimental design in single-cell epigenomics.

## 5 Methods

### 5.1 Datasets and pre-processing

We processed all scATAC-seq datasets under a uniform pipeline to minimise variation arising from different pre-processing approaches and sequencing technologies (Table 1). Raw or processed data were downloaded from the Gene Expression Omnibus (GEO) [25] or the source repositories described below. Peak-by-cell count matrices were generated using the PIC counting() function from the PICsnATAC R package (v0.2.3) [13], which applies a paired-insertion counting model to fragment files. The resulting matrices were then imported into ChromatinAssays using the Signac package (v1.11.9) [26] and Seurat (v4.9.9) [27] with no initial cell or feature filtering (min.cells=min.features= 0).

Cell type labels were assigned using published metadata when available. For the two 10x Genomics PBMC datasets, labels were assigned by transferring annotations from paired single-cell RNA-seq data. Peaks were retained if detected in at least 1% of cells within a given cell type and dataset. The number of cells and peaks per cell type are provided in Supplementary Table S6. Cell types with fewer than 200 cells were excluded to ensure sufficient data for analysis. Finally, the filtered matrices were converted into SingleCellExperiment (SCE) objects for all downstream analyses.

The following six datasets were used:

- **10x Genomics PBMC datasets**: Two peripheral blood mononuclear cell (PBMC) datasets were downloaded directly from 10x Genomcis website: PBMC10k (ID:10k_pbmc_ATACv2_nextgem_Chromium_Controller) and PBMC5k (ID:atac_v1_pbmc5k).
- **Satpathy et al**. [18]: Processed count matrix, fragment files and cell metadata were downloaded from GEO (accession GSE129785).
- **Janssens et al**. (Fly brain) [21]: Peak regions, cell barcodes, and cell metadata were extracted from the cisTopic object L3P12 cisTopic.Rds, downloaded from flybrain. Cells with CellType lvl1 labels ‘unk’ or - were excluded as unannotated.
- **Cusanovich et al**. [20]: BAM files and cell metadata were downloaded from mouse-atac. Fragment files required as input for PIC counting were generated using Python package sinto (v0.10) with default parameters. This atlas dataset consists of chromatin accessibility profiles across 13 different mouse tissues; from which we randomly selected three tissues: cerebellum, kidney, and spleen to represent distinct tissue contexts while keeping computational demands tractable.
- **Buquicchio et al**. (Liver dataset): Fragment files were downloaded from GEO (accession GSE199799) and processed using ArchR as described in the original manuscript [19].
- **Alzheimer’s disease dataset (population-scale analysis)**. The dataset from Xiong et al. [22] was downloaded from the Synapse portal as described in the original publication. Sample names were harmonised to match individual identifiers, yielding a final cohort of 83 individuals.

### 5.2. simPIC simulation framework

The simPIC simulation framework has four key components: (i) estimating parameters from empirical data at both single sample and population levels, (ii) simulating cell groups and batches for a single sample, (iii) modelling peak means with genetic effects across a population, and (iv) simulating single-cell counts for individual and population-scale data.

#### 5.2.1. Parameter estimation

##### a) Single-sample parameter estimation

Single-sample simulation requires three key parameters estimated from a user-supplied peak-by-cell count matrix or SCE object using the simPICestimate() function: library size, peak-mean, and cell sparsity. Library size is modelled using a log-normal distribution, with parameters mean(*µ*) and standard deviation (*σ*). Peak-mean is computed after normalizing each cell to the dataset’s median total count. Peaks with zero counts across all cells are removed (i.e., peaks for which every single entry is zero) before parameter estimation. A lognormal-gamma mixture distribution is then fitted to the non-zero peak means using the fitdistrplus package [28] (Supplementary Note 1). This estimates the gamma shape (*α*) and rate (*β*), the lognormal location (*µ*) and scale (*σ*), and the mixture weight *w*. Alternative distributions from the generalized gamma family (e.g., gamma,Weibull, Pareto) are also available and may be selected when they better fit the data. Cell-level sparsity is modeled by estimating the proportion of zero entries per cell and fitting a Bernoulli distribution with parameter *π*.

When simulating multiple cell groups or subpopulations (nGroups *>* 1) within a single sample, parameter estimation is performed separately within each group, capturing group-specific variation in accessibility and sparsity profiles. When batch effects are included (nBatches *>* 1), they are incorporated by applying peak-wise multiplicative factors across all cells within a batch, using either internally assigned or user-provided batch labels. When both multiple groups and batches are specified, simPIC models biological variability using the biological coefficient of variation (BCV), which captures the mean-variance relationship typical of scATAC-seq data [29, 30]. As shown in simPIC model framework (Figure 2), we first calculate a base cell mean 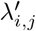 derived from normalized peak means and library size. To incorporate biological noise, we estimate a peak-specific BCV (*B*_*i,j*_) using a scaled inverse chi-square distribution: 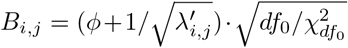, where *ϕ* is the common dispersion and *df*_0_ represents the prior degrees of freedom (see Supplementary Note 2 for detailed BCV calculation). The final trended cell mean *λ*_*i,j*_ is then sampled from a gamma distribution, 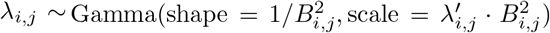. This stochastic transformation ensures that *λ*_*i,j*_ maintains an expected value equal to the base accessibility 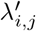 while reflecting realistic biological dispersion.

Since edgeR has been observed to overestimate the common dispersion in single-cell synthetic data [5], we empirically calibrated a linear correction by comparing edgeR common dispersion estimates to the known true dispersions in simulations parameterised from six heterogenous real scATAC-seq datasets spanning diverse organisms, platforms and sequencing technologies (Table 1). All six datasets exhibited a similar inflation pattern, and fitting a single linear model across them yielded the fixed correction 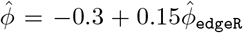 which is applied by default to better match observed mean–variance relationships (Supplementary Note 3, Supplementary Figure S1). This correction is an empirically calibrated default rather than a dataset-specific adjustment; users may disable it or provide alternative scaling by directly specifying BCV parameters (e.g. bcv.common and bcv.df) via setsimPICparameters(). All estimated parameters are stored in a simPICcount object for downstream simulation and default settings are available for exploratory runs.

##### b) Population-scale parameter estimation

Population-scale simulation incorporates inter-individual variability using an approach adapted from splatPop [6]. Accessibility counts from a peak-by-cell matrix annotated with individual identities are first aggregated across cells to generate a peak-by-individual matrix for parameter estimation (Figure 2, Supplementary Table S1). Parameter estimation then follows a two-step procedure. First, population-wide peak-mean parameters were estimated by fitting a ln-gamma mixture distribution to the empirical distribution of mean accessibility values aggregated across individuals. The model estimates the gamma shape (*α*_*p*_) and rate (*β*_*p*_), the lognormal location (*µ*_*p*_), and scale (*σ*_*p*_) and the mixture weight *w*_*p*_. Second, to account for the mean-variance relationship, peaks are grouped into bins of 50 based on mean accessibility (as in splat-Pop). The coefficient of variation (CV) is computed within each bin, and a gamma distribution is fit to these values to estimate bin-specific shape (*α*_*v*_) and rate (*β*_*v*_) parameters. Parameters controlling for caQTL effect sizes (*α*_*c*_) and (*β*_*c*_) are estimated by fitting a gamma distribution to effect sizes from an empirical caQTL mapping study [22]

#### 5.2.2. Simulating cell-groups and batches for single sample

simPIC simulates samples with multiple cell groups by defining distinct cell groups where peaks exhibit differential accessibility. Consistent with previous reports that approximately 13% of peaks are differentially accessible in single-cell ATAC-seq data [31], a multiplicative differential accessibility (DA) factor is assigned to each peak *i* and applied to its baseline mean (*λ*_*i*_). DA factors for differentially accessible peaks are sampled from a log-normal distribution; non-differentialy accessible peaks are assigned a factor of one. The number of groups and cell membership probabilities are user-defined, as are parameters governing the probability of DA and the magnitude and direction of DA factors per group. The resulting SCE object contains group assignments and corresponding DA factors for each cell.

Batch effects represent technical variation introduced during sample handling, preparation, or processing that can obscure true biological signals. simPIC models these effects by applying multiplicative scaling factors to peak accessibility values across cells within each batch. Specifically, for each peak *i* and batch *b*, a peak-wise scaling factor 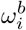 is drawn from a log-normal distribution, 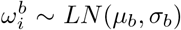, and applied to all cells assigned to batch *b*. These factors scale the expected peak means (i.e., the base mean 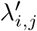 for cells in batch *b* is multiplied by 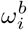), thereby inducing a systematic batch-specific shift in peak accessibility rather than unstructured per-cell noise (Figure 2, Supplementary Table S1). Users specify the number of batches *nBatches* and the number of cells per batch *batchCells*, the total number of cells equal to the sum of *batchCells*. Batch effects are simulated independently of any batch annotations or batch structure in the reference dataset, allowing users to generate arbitrary user-defined batch configurations for benchmarking purposes. The magnitude of batch effects is governed by the log-normal parameters (default *batch*.*facLoc* and *batch*.*facScale* in the simPICcount object), which can be modified via setsimPICparameters(). Adjusting these parameters allows control over the degree of batch variation, with larger or more variable scaling factors producing stronger batch effects.

#### 5.2.3. Simulating peak means with genetic effects across a population

To simulate population-scale single-cell ATAC-seq data, we created a wrapper around splatPop extending the simPIC framework [6] by adapting splatPop’s gene-based model to chromatin accessibility peaks. For each peak *i*, the population-wide accessibility mean (*λ*_*i*_) was sampled from a ln-gamma mixture distribution. This distribution is parameterised by the gamma shape (*α*) and rate (*β*), the lognormal location (*µ*), and scale (*σ*) and the mixture weight *w*. Peak-specific variability (*σ*_*i*_) was sampled from a gamma distribution, with an optional scaling factor (*s*) to adjust overall variability. Baseline accessibility for each peak in each individual (*λ*_*i,j*_) was drawn from a normal distribution centered on (*λ*_*i*_) and scaled by (*σ*_*i*_). When empirical data were provided to the splatPopSimulate() function, these sampling steps were bypassed and empirical means and variances were used directly.

caQTL effects are simulated using real genotype and peak annotation data from Xiong et. al [22], including VCF and GFF files. Parameters including the proportion of accessible peaks associated with caQTLs (caPeaks), the minor allele frequency of the associated SNPs (caSNPs), and the distance between each caPeak and its linked SNP were defined to reflect the population structure of the reference dataset. caQTL effect sizes (*ω*_*i*_) are sampled from a gamma distribution with parameters *α*_*c*_ and *β*_*c*_ and applied to peak accessibility based on genotype (*G*_*i,j*_, encoded as 0, 1, or 2). The framework also supports group-specific caQTLs affecting defined cell groups, and condition-specific effects tailored to specific cohorts.

Differential accessibility (DA) between groups or conditions is simulated by scaling peak accessibility using factors drawn from a log-normal distribution with parameters (*µ*_da_, *σ*_da_). Separate parameters for cell groups and cohorts allow independent control of these effects and the proportion of peaks assigned negative DA factors (reduced accessibility) can be specified to control directional heterogeneity across the simulated dataset.

#### 5.2.4. Simulating single-cell counts for individual and population scale data

Single-cell scATAC-seq peak-by-cell count matrices are generated using two core functions: simPICsimulate() for one cell group simulations and simPICsimulateGroup() for multi-group or batch simulations. In both cases, peaks are simulated conditionally independently given the peak-wise and cell-wise parameters (e.g., peak means/dispersion, library size, and sparsity) and any specified group, batch, or population effects. simPIC therefore targets marginal and mean-variance properties and does not explicitly model peak-peak covariance (co-accessibility).

For single-group simulation peak means (*λ*_*i*_) are drawn from a ln-gamma distribution and cell-specific library sizes (*L*_*j*_) are sampled from a log-normal distribution. Counts *Y*_*ij*_ are sampled from a Poisson distribution with mean parameter 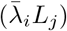. Sparsity is introduced by a sparsity indicator *Z*_*ij*_ controlled by parameter *π*_*j*_ which randomly zeros out entries using a Bernoulli-distribution to reflect highly sparse nature of scATAC-seq data.

For multi-group and batch simulations, peak means are normalized and scaled by library size to compute cell-adjusted values (*λ*_*i,j*_) which are then sampled from a gamma distribution. Biological variability is further captured using a biological coefficient of variation (BCV), computed from user-defined parameters (*ϕ, df*_0_) and used to scale dispersion across peaks. Larger mean values result in lower BCV, consistent with the inverse mean-variance relationship in scATAC-seq data. Counts are then sampled and sparsity introduced as in single-group simulation. Population scale counts are simulated as described in splatPop [6].

### 5.3. Comparative analysis of simulation frameworks

Tools were included in the benchmark if they satisfy three criteria: (i) the simulation output is a peak-by-cell count matrix, enabling direct comparison with real scATAC-seq data; (ii) a publicly available implementation exists; and (iii) the tool executes successfully on at least one benchmark dataset. Under these criteria, we compared four simPIC distribution variants (Weibull, gamma, lognormal-gamma, and Pareto) against four competing simulators: simCAS [10], scDesign3 [12], DiTSim [15], and scMultiSim [14].

Each comparator was run using its recommended interface and default parameters unless otherwise noted. simCAS was executed via its command-line interface in pseudo-cell-type mode; input SCE objects were converted to AnnData format (as required by simCAS) using the SCE2AnnData() function from the zellkonverter R package (v3.18) [32] and converted back to SCE format after simulation using AnnData2SCE. scDesign3 was run using the scdesign3 function from the scDesign3 R package (v1.0.0) with the family used set to negative binomial and copula set to gaussian. DiTSim was executed using its default diffusion transformer pipeline; count matrix output was used directly. scMultiSim was run using the sim true counts function with *tree* = *Phyla*1() and no GRN specification. All tools were applied to simulate the same number of cells and peaks as in the reference real dataset for each cell type. Comparison was performed across 25 celltypes from six datasets (Table 1). Input objects were read as peak-by-cell count matrices; supported object types were Single-CellExperiment, dgCMatrix, and Matrix. For SCE objects, the counts assay was used when present and X otherwise (e.g. DiTSim). Furthermore, since no existing tool simulates population-scale scATAC-seq data with inter-individual genetic variation, we did not perform comparisons in that setting. Simulated data were evaluated against real data using the simPICcompare() function, which generates diagnostic plots to assess similarity across key features.

### 5.4. Evaluation metrics

Simulation performance was evaluated across three key characteristics: library size, peak mean, and cell sparsity. For each matched real–simulated pair, real and simulated value vectors were first aligned by quantile on a common grid of *n* = 1024 points, four error metrics were then computed from the aligned vectors: Median Absolute Deviation (MAD), Mean Absolute Error (MAE), Root Mean Square Error (RMSE), and 1 minus Pearson Correlation Coefficient (1 − PCC). Together these metrics capture both central tendency and tail behaviour of distributional discrepancy.

To complement the quantitative metrics, we computed two-sample Kolmogorov–Smirnov (KS) statistics for each property. The KS statistic is defined as *D* = sup_*x*_ |*F*_real_(*x*) − *F*_sim_(*x*)|, where *F* denotes the empirical cumulative distribution function. KS values were computed per cell type for each tool and property, and summarised across tools and properties in a summary heatmap (Figure 3j).

Distributional agreement was further assessed visually using kernel density estimates for each property computed separately for real and simulated data. Density difference plots were constructed by subtracting the real kernel density estimate from the simulated estimate on a common evaluation grid, with positive values indicating regions where the simulator overestimates density and negative values indicating underestimation. Library size and peak mean distributions were plotted on a log_1p_ scale; cell sparsity was plotted on a proportion scale. These plots are shown in Figure 3.

To aggregate performance across properties and metrics, a composite rank score was computed for each method within each matched cell type. Within each dataset × cell type × property × metric combination, methods were ranked by ascending error value (lower error = rank 1). The composite rank for each method within each cell type was then defined as the mean rank across all three properties and all four metrics, yielding 12 rank values per cell type. The distribution of composite rank across cell types per method is shown in Figure 4a.

To assess whether differences in composite rank between simPIC variants and competing methods were statistically supported, each simPIC variant was compared against each competing tool using matched cell types only. For each pairwise comparison, paired differences in composite rank were computed per cell type. The median paired difference, bootstrap 95% confidence interval for the median, Wilcoxon signedrank test *p*-value, and Benjamini–Hochberg (BH) adjusted FDR were computed for each comparison. A negative median paired difference indicates better performance of the simPIC variant relative to the comparator. Results are shown in Figure 4b.

To assess whether simulated data preserves global peak–peak dependence, we additionally computed pairwise peak–peak correlations on TF–IDF transformed accessibility for the top 2,000 variable peaks (after filtering peaks detected in ≥ 10 cells). Distributions of signed and absolute correlations |*r*| between real and simulated data were compared, with tail behaviour summarised using the 95th/99th percentiles of |*r*| and the number of peak pairs with |*r*| *>* 0.3 and |*r*| *>* 0.5 (Supplementary Fig. S2; Supplementary Table S5).

### 5.5. Computational efficiency

Computational efficiency was benchmarked for all eight methods compared in this study: simPIC (gamma, Weibull, lognormal-gamma, and Pareto variants), simCAS, scDesign3, DiTSim, and scMultiSim. Runtime (seconds) and peak memory usage (MB) were recorded for each method across all matched dataset–cell type combinations (Figure 5).

To assess overall differences across methods, Friedman tests were applied separately for runtime and peak memory, blocking by dataset × cell type. Pairwise differences were evaluated using paired Wilcoxon signed-rank tests applied separately for runtime and peak memory, with *p*-values adjusted using BH FDR correction. Runtime and peak memory distributions are shown as boxplots with jittered points.

To characterise how runtime and memory scale with dataset size, we additionally fitted separate log–log linear regression models for each method regressing runtime and peak memory against number of cells and number of peaks. Four scaling plots are shown in Figure 5 c-f.

For simPIC, scDesign3 and scMultiSim, elapsed time was recorded using R’s system.time and peak memory was measured using the mem used() function from the lobstr package (v1.2.1). For simCAS and DiTSim runtime and memory were recorded using Python’s time and tracemalloc modules, respectively.

### 5.6. caQTL mapping analyses

We simulated a larger dataset based on parameter estimates from library 5 of the microglia cell type from the Xiong et.al, dataset [22]. The simulation included 100 cells per individual for 83 samples, matching the real dataset, within a single batch. caQTL effects were assigned to 70% of the simulated peaks. Single-cell counts were normalized using scran (v1.28.2), then aggregated by individual through mean aggregation and quantile normalized to a standard normal distribution [33]. caQTL mapping was conducted on all SNPs within 100 kb upstream and downstream of each peak on chromosome 22 using a linear mixed model implemented in LIMIX [23]. The model included SNP genotype as a fixed effect, the top 15 principal components from chromatin accessibility PCA as fixed effects to control unwanted variation, and a kinship matrix calculated with PLINK (v1.90) [34] as a random effect to adjust for population structure. Associations with empirical feature p-values below 0.005 were considered significant. Manhattan plots showing the full chromosome 22 and regional 400 kb windows were generated for both real and simulated data.

### 5.7. Differential accessibility benchmarking

To evaluate the ability of DA methods to recover known signal under controlled conditions, we generated synthetic scATAC-seq datasets using simPIC’s multi-group simulation mode (nGroups = 2), in which DA factor assignments constitute a known ground truth. Simulations were parameterised separately for six representative datasets spanning diverse biological contexts and cell counts (Table 1 and Supplementary Table S6). To reflect realistic cell numbers, the number of cells per replicate was scaled to each reference cell type as cells_per_rep = ⌊*N*_cells_*/*(*n*_rep_ × 2)⌋, with *n*_rep_ = 4 replicates per group (8 pseudo-replicates per cell type in total). The resulting cells per replicate were: 275 (CD14 Monocytes, PBMC10k), 1800 (CD34 Progenitors, Satpathy), 575 (Liver TRM, Buquicchio), 275 (B cells, Cusanovich), 40 (OL Type I NB, Fly Brain), and 100 (CD8 Effector T cells, PBMC10k). DA factors for differentially accessible peaks were drawn from a log-normal distribution; non-differentially accessible peaks were assigned a factor of one. Ground truth DA status and factor magnitude are known by construction for each simulation.

Ten DA methods implemented in the Libra R package were evaluated using this design, following the comparative framework of Teo et al. (2024) [24]: edgeR quasilikelihood F-test (edgeR-QLF), edgeR likelihood ratio test (edger-LRT), DESeq2 Wald test, DESeq2 likelihood ratio test (DESeq2-LRT), limma voom, limma trend, logistic regression, Wilcoxon signed-rank test, negative binomial regression, and *t*-test. All methods were applied under their default recommended settings as described in Teo et al.[24]

Performance was assessed against the known simulation ground truth using five metrics: true positive rate (TPR), area under the receiver operating characteristic curve (AUC-ROC), and area under the precision–recall curve (AUC-PR). TPR was computed at a significance threshold of FDR *<* 0.05. AUC-ROC and AUC-PR provide threshold-independent summaries of discriminative performance. This analysis constitutes a parallel design applied to simPIC output and is intended to characterise the relative sensitivity and specificity of DA methods under the simPIC generative framework.

The ground truth in this benchmarking is defined by the simulation model’s assumptions regarding the distribution of DA factors and the proportion of accessible peaks subject to differential accessibility. Results therefore characterise method behaviour under the simPIC generative model and should be interpreted within this scope. Method rankings may not generalise to all real biological contexts, particularly where the true DA effect size distribution or peak-accessibility structure differs substantially from the simulated setting.

## 6 Data and code availability

simPIC is available on Bioconductor https://bioconductor.org/packages/release/bioc/html/simPIC.html and on GitHub https://github.com/sagrikachugh/simPIC. Data and code to reproduce the figures in this manuscript are available under CC BY license on GitHub repository https://github.com/sagrikachugh/simPIC-paper, a website accompanying the analysis is available here https://sagrikachugh.github.io/simPIC-paper/

## 7 Acknowledgements

The authors would like to thank Chun Fung Kwok for discussions on implementation of mixture model distributions, and Aaron Wing Cheung Kwok and Jeffrey Pullin for helpful discussions on scATAC-seq data.

## 8 Funding

S.C. was supported by Melbourne Research Scholarship, Graeme Clark Institute of Biomedical Research Top-Up Scholarship and St. Vincent’s Institute of Medical Research Top-Up Scholarship. This work was further supported by funding from the National Health and Medical Research Council of Australia (GNT1195595) and the National Institutes of Health of the United States of America (R01HG011886) awarded to D.J.M.

## 9 Author contribution

D.J.M, H.S. and S.C. conceived the study. S.C developed the simPIC software, conducted the study and wrote the manuscript. D.J.M. and H.S. supervised the study. All authors contributed to, read, and approved the final manuscript.

## Notes

### Competing Interest Statement

The authors have declared no competing interest.

### Summary of Updates

Extensively revised all sections of the manuscript including supplementary data.

