## Supplementary Figures for "simPIC: flexible simulation of single-cell ATAC-seq paired-insertion counts from individuals to populations"

***Supplementary Note 1: Distribution for modelling peak means***

We provide four distributions for simulating peak means ($\lambda$), namely Weibull, gamma and lognormal-gamma mixture(default), and Pareto.

The probability density function (PDF) of the lognormal-gamma mixture model is given by:

$$f_{\text{lngamma}}(x|\alpha,\beta,\mu,\sigma^2) = \pi \cdot f_{\text{Gamma}}(x|\alpha,\beta) + (1-\pi) \cdot f_{\text{Lognormal}}(x|\mu,\sigma^2) \quad (1)$$

where $\pi$ is the proportion of non-zero elements/counts ($0 \leq \pi \leq 1$), $\alpha$ and $\beta$ are the shape and rate parameter of the gamma distribution respectively,$\mu$ and $\sigma^2$ are the mean and variance of the lognormal distribution,

The lognormal-gamma mixture model is particularly useful when the data exhibit characteristics that could be modeled by both a gamma distribution (for positive skewness and flexibility in shape) and a lognormal distribution (for right-skewed data where values are strictly positive and logarithmic transformations are appropriate).

***Supplementary Note 2: Mathematical Derivation of BCV and Trended Means***

The biological coefficient of variation (BCV) for each peak $i$ in cell $j$ ($B_{i,j}$) is simulated to reflect the mean–variance trend observed in scATAC-seq data. Using the common dispersion $\phi$ and prior degrees of freedom $df_0$ estimated via *edgeR*, $B_{i,j}$ is derived as follows:

**1. BCV Calculation**

If $df_0$ is finite, $B_{i,j}$ is sampled using a scaled inverse chi-square distribution:

$$B_{i,j} = \left( \phi + \frac{1}{\sqrt{\lambda'_{i,j}}} \right) \cdot \sqrt{\frac{df_0}{\chi^2_{df_0}}} \tag{2}$$

where $\chi^2_{df_o}$ is a chi-square-distributed variable with $df_o$ degrees of freedom. In the limiting case where $df_0$ is infinite, the BCV simplifies to the global trend:

$$B_{i,j} = \phi + \frac{1}{\sqrt{\lambda'_{i,j}}} \tag{3}$$

This formulation ensures that peaks with lower base accessibility ($\lambda'_{i,j}$) exhibit higher relative variability, accurately reflecting the sparsity-driven noise in scATAC-seq data.

**2. Gamma Sampling for Trended Means**

To incorporate this variability, the final trended cell mean ($\lambda_{i,j}$) is sampled from a Gamma distribution. The distribution is re-parameterized using the base mean and the simulated BCV to ensure the expectation $E[\lambda_{i,j}] = \lambda'_{i,j}$:

$$\lambda_{i,j} \sim \text{Gamma} \left( \text{shape} = \frac{1}{B^2_{i,j}}, \text{scale} = \lambda'_{i,j} \cdot B^2_{i,j} \right) \tag{4}$$

By defining the shape as $1/B^2_{i,j}$ and the scale as $\lambda'_{i,j} \cdot B^2_{i,j}$, the model captures the biological dispersion of the reference dataset while maintaining the discrete, stochastic nature of single-cell chromatin accessibility.

##### ***Supplementary Note 3: Dispersion estimate***

It is well established that scATAC-seq data exhibits a strong mean-variance trend, where regions with higher mean accessibility tend to show greater variability [1, 2]. This pattern arises due to inherent biological variability, technical noise in single-cell measurements and fundamental statistical property of count data. To accurately replicate this trend in simulated data, I enforce it in extended simPIC framework (Main Figure 1) by estimating biological coefficient of variation (BCV) from real data, with parameters $\phi$ and $df_0$ from a scaled inverse chi-squared distribution using the `estimateDisp` function from the `edgeR` package [3], where the scaling factor is a function of the peak mean. This ensures that simulated datasets capture realistic variability patterns observed in real single-cell ATAC-seq experiments. However, upon testing this estimation procedure on simulated datasets, I observed that the `edgeR` estimate of common dispersion was inflated, as shown in Figure S1. This inflation has also been reported in the simulation of single-cell RNA-seq data using the `splatter` package [4]. To address this issue, I applied a linear correction to the edgeR-estimated dispersions using the formula $\hat{\phi} = -0.3 + 0.15\hat{\phi}_{\texttt{edgeR}}$.The coefficients of the linear dispersion correction were obtained from a joint regression of true dispersion values against edgeR's common dispersion estimates using simulations parameterised from six distinct real scATAC-seq datasets (Table X). Despite differences in organism, platform, and sequencing technology, all datasets showed a highly similar inflation of edgeR-estimated dispersions. Applying a single set of coefficients consistently reduced this inflation and improved agreement between simulated and empirical mean–variance relationships. Owing to this consistency, we summarise the behaviour using a single representative example in Supplementary Figure S1 and adopt the resulting correction as an empirically calibrated default in simPIC. This correction helps to adjust for the overestimation of dispersion, leading to more accurate simulated data that better reflects the true variability observed in real datasets.

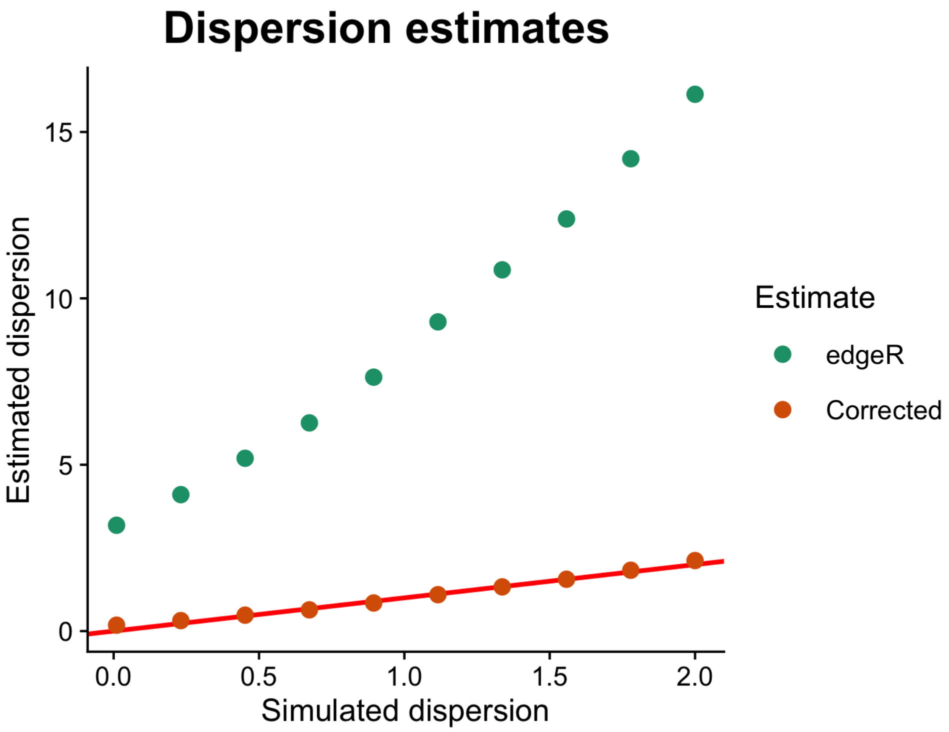


**Fig. S1** Scatter plot of estimated dispersions against the true simulated values. Estimates of common dispersion obtained from edgeR (green) can be inflated for single-cell data. The SIMPIC simulation uses a linearly corrected value (orange) in its estimation procedure. The red line shows the true values estimates.

### Supplementary Figures

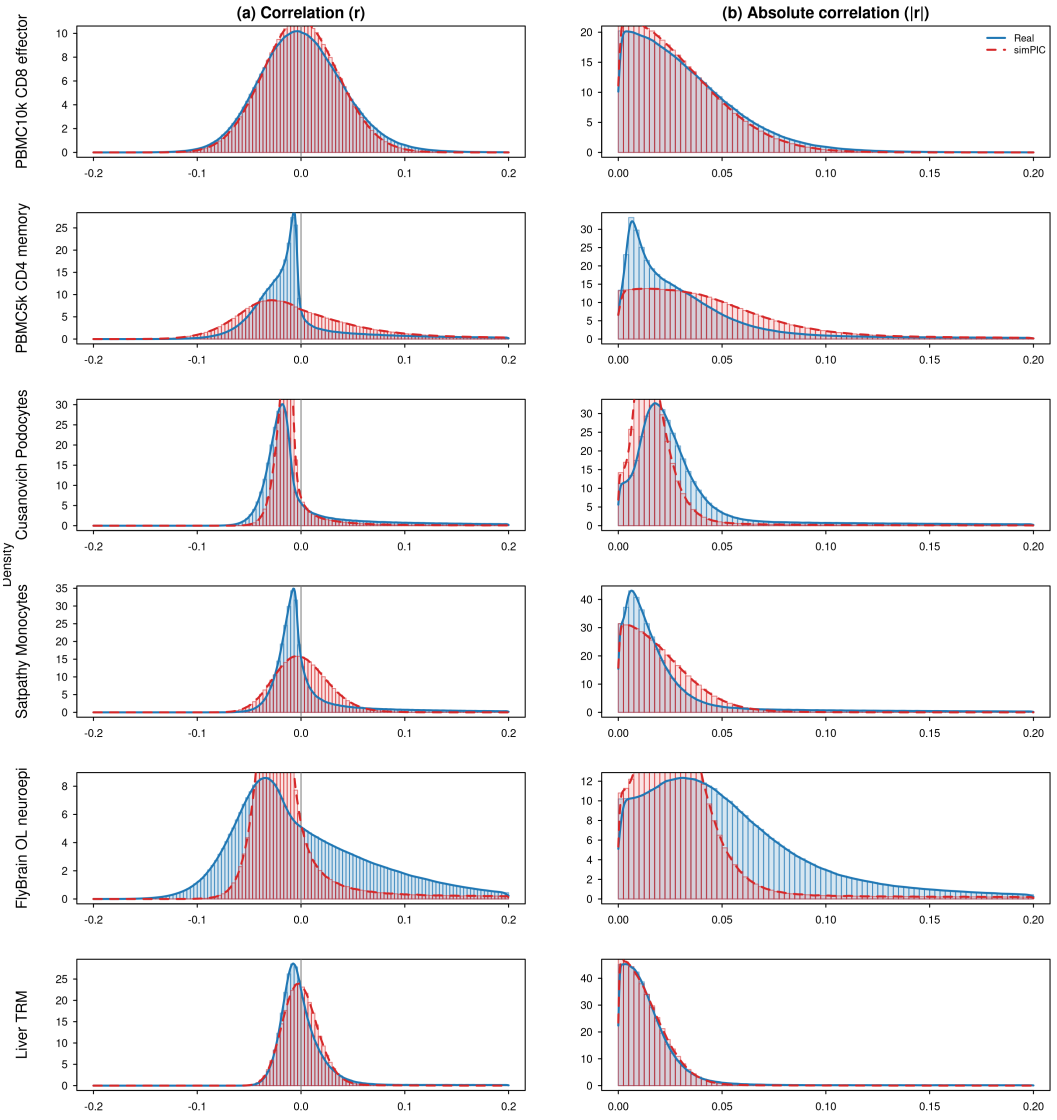


**Fig. S2** Peak–peak correlation structure in real versus simPIC data. Pairwise peak–peak correlations across cells were computed on TF–IDF transformed accessibility using the top 2,000 variable peaks (filtered to peaks detected in $\geq 10$ cells). (a) Signed correlations ($r$). (b) Absolute correlations ($|r|$). Blue solid: real; red dashed: simPIC.

#### Supplementary Figure S3

The below figures show all the 25 cell types evaluated in this study. The figures are divided into two pages for each cell type with cell type name indicated at top.

All figures present the same set of diagnostics in different cell types across the six datasets.

**Top panels (a–i)** show cell-level and peak-level summary distributions: (a) log library size, (b) log peak mean, (c) cell sparsity; (d–f) corresponding density distributions of log library size, log peak mean and cell sparsity; (g–i) density differences relative to real data.

**The next page is of the same cell type and shows additional peak-level characteristics and mean-variance relationships:** (a) peak sparsity, (b) log peak variance, (c) peak mean–variance relationship, (d) peak mean vs non-zero proportion, (e) peak mean vs peak sparsity; (f–g) density overlaps for peak sparsity and log peak variance; (h–i) corresponding density differences.

The numbers on top of violins indicate KS statistic.

Cusanovich_Kidney_Podocytes

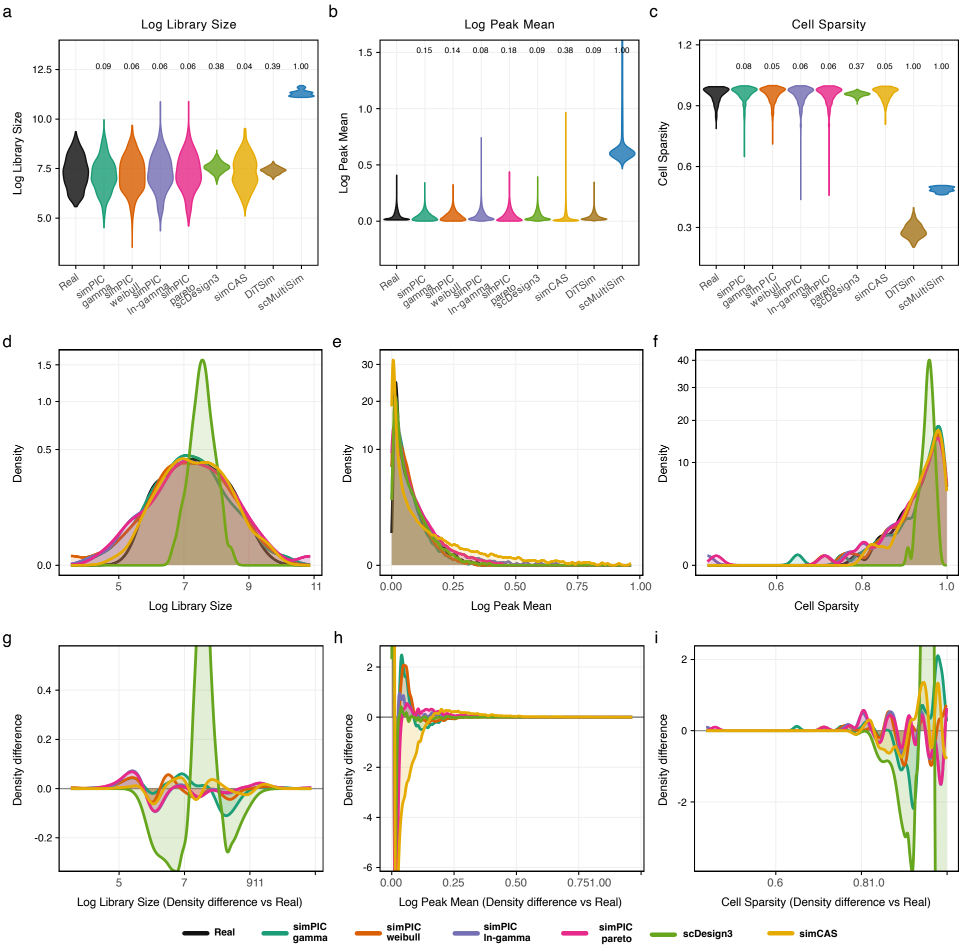

Cusanovich_Kidney_Podocytes

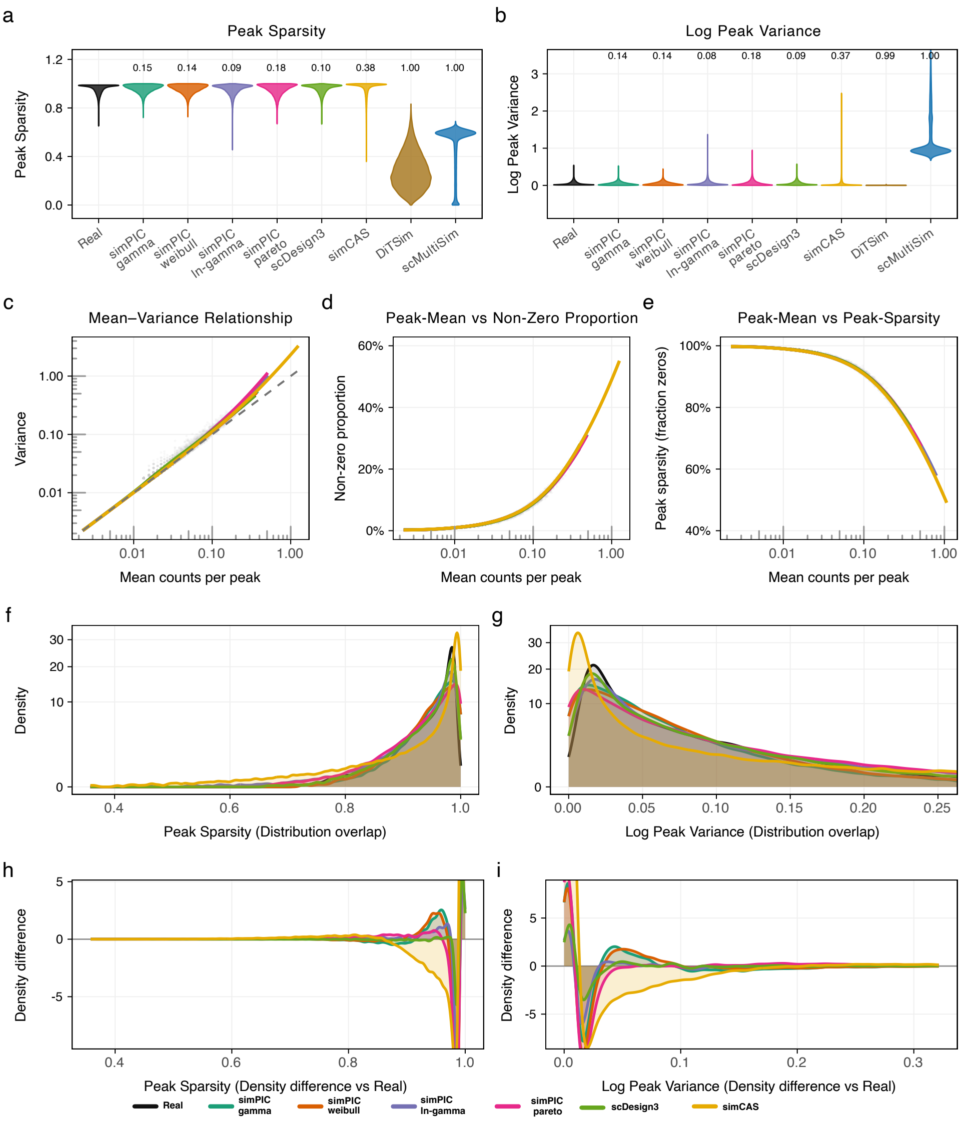

Cusanovich_Cerebellum_Astrocytes

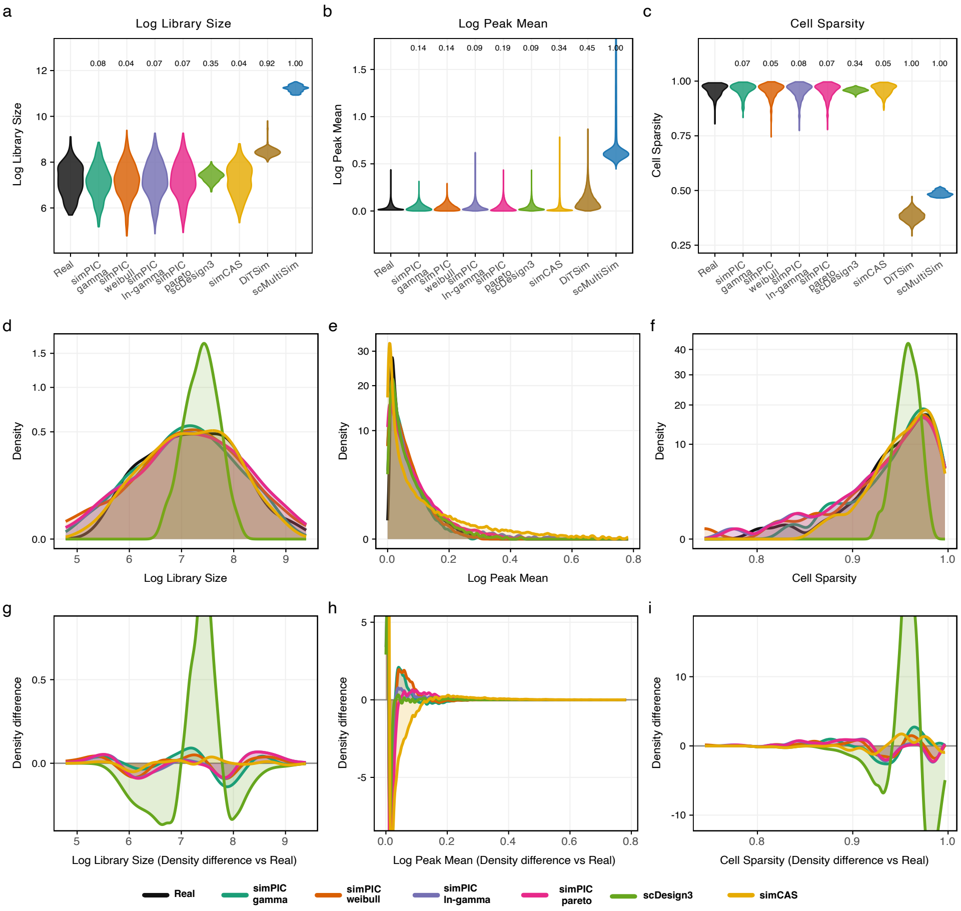

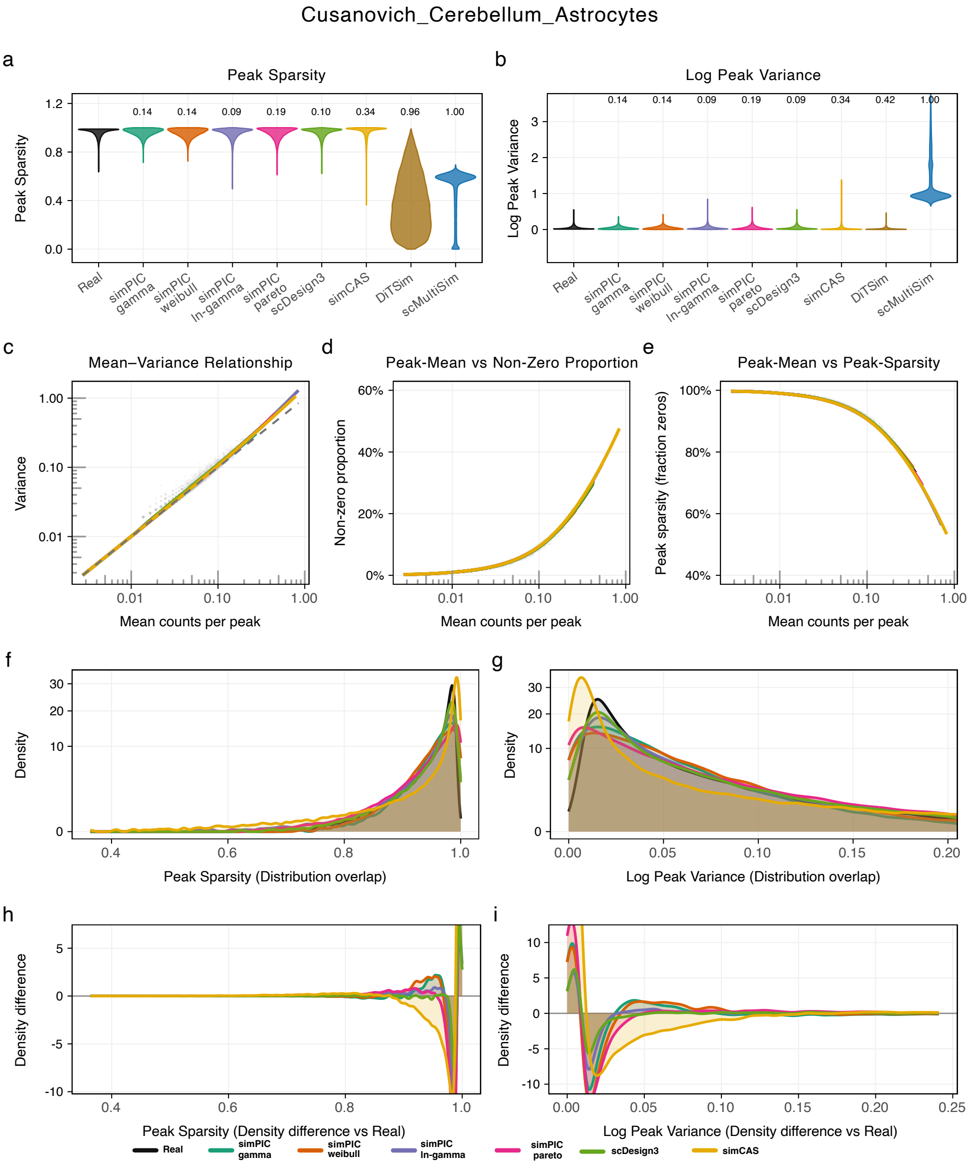
Cusanovich_Cerebellum_Astrocytes
a
Peak Sparsity
b
Log Peak Variance
c
Mean–Variance Relationship
d
Peak-Mean vs Non-Zero Proportion
e
Peak-Mean vs Peak-Sparsity
f
g
h
i
Peak Sparsity
Log Peak Variance
Variance
Non-zero proportion
Peak sparsity (fraction zeros)
Mean counts per peak
Density
Density difference
Peak Sparsity (Distribution overlap)
Log Peak Variance (Distribution overlap)
Peak Sparsity (Density difference vs Real)
Log Peak Variance (Density difference vs Real)
Real
simPIC gamma
simPIC weibull
simPIC ln-gamma
simPIC pareto
scDesign3
simCAS
DiTSim
scMultiSim

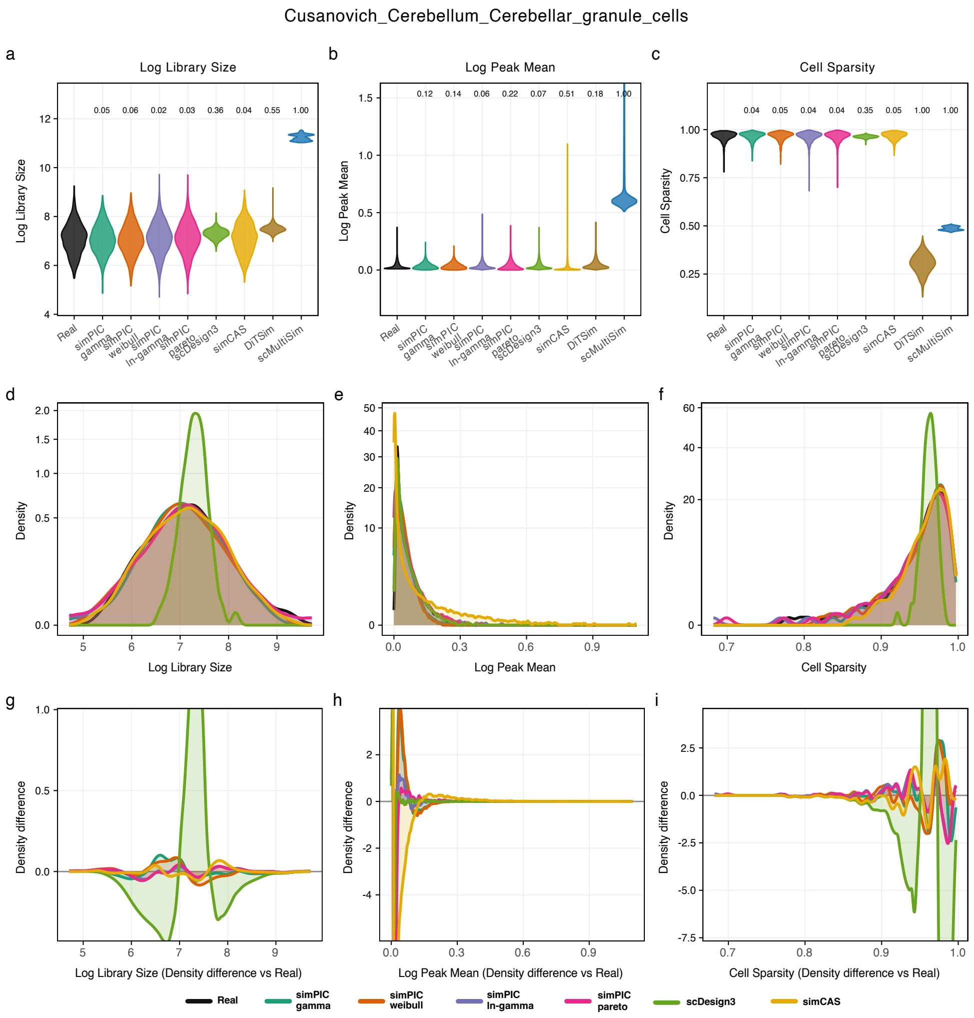

Cusanovich_Cerebellum_Cerebellar_granule_cells
Log Library Size
Log Peak Mean
Cell Sparsity
Density
Density difference
Log Library Size (Density difference vs Real)
Log Peak Mean (Density difference vs Real)
Cell Sparsity (Density difference vs Real)
Real
simPIC gamma
simPIC weibull
simPIC ln-gamma
simPIC pareto
scDesign3
simCAS
DiTSim
scMultiSim

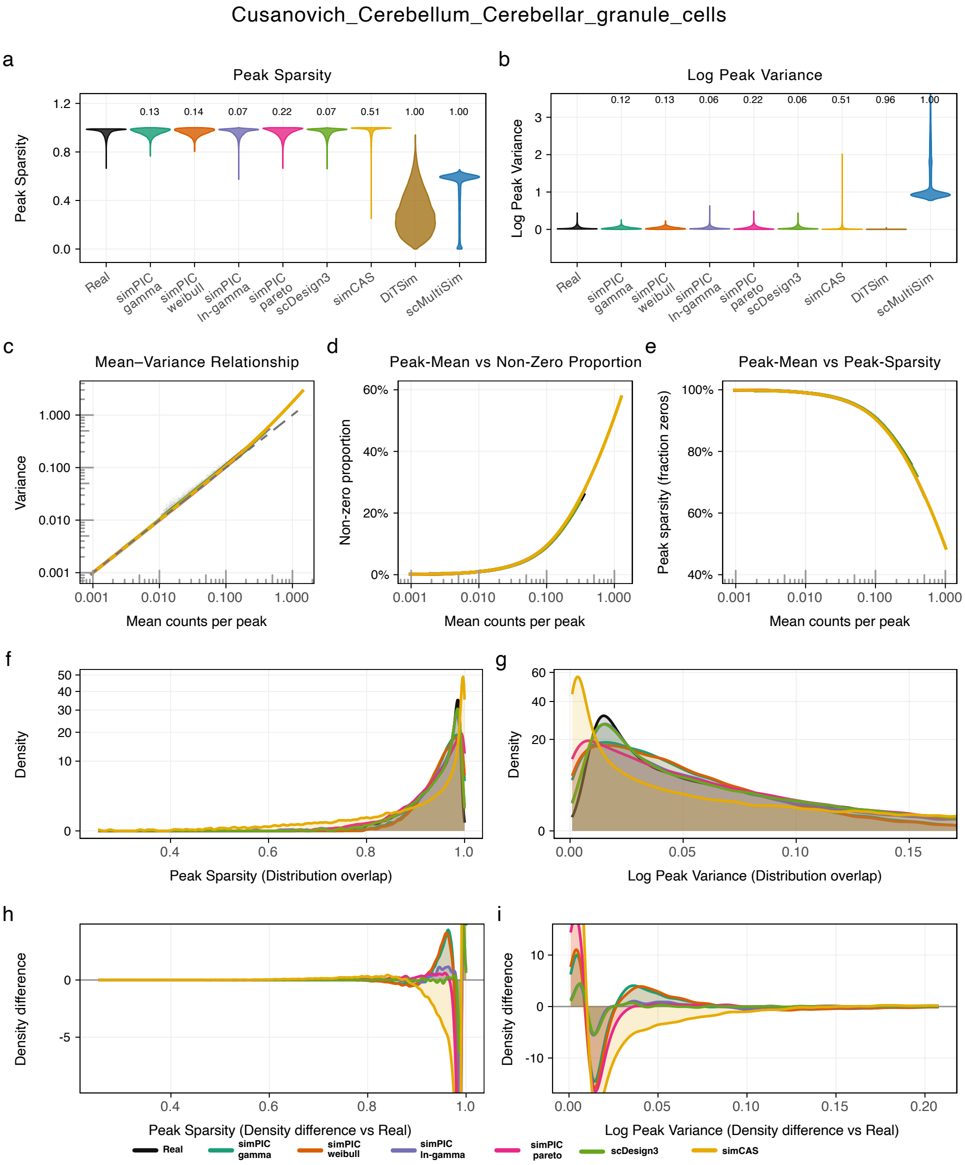
Cusanovich_Cerebellum_Cerebellar_granule_cells
a
Peak Sparsity
Peak Sparsity
0.13 0.14 0.07 0.22 0.07 0.51 1.00 1.00
Real simPIC gamma simPIC weibull simPIC ln-gamma simPIC pareto scDesign3 simCAS DiTSim scMultiSim
b
Log Peak Variance
Log Peak Variance
0.12 0.13 0.06 0.22 0.06 0.51 0.96 1.00
c
Mean–Variance Relationship
Variance
Mean counts per peak
d
Peak-Mean vs Non-Zero Proportion
Non-zero proportion
Mean counts per peak
e
Peak-Mean vs Peak-Sparsity
Peak sparsity (fraction zeros)
Mean counts per peak
f
Density
Peak Sparsity (Distribution overlap)
g
Density
Log Peak Variance (Distribution overlap)
h
Density difference
Peak Sparsity (Density difference vs Real)
i
Density difference
Log Peak Variance (Density difference vs Real)
Real
simPIC gamma
simPIC weibull
simPIC ln-gamma
simPIC pareto
scDesign3
simCAS

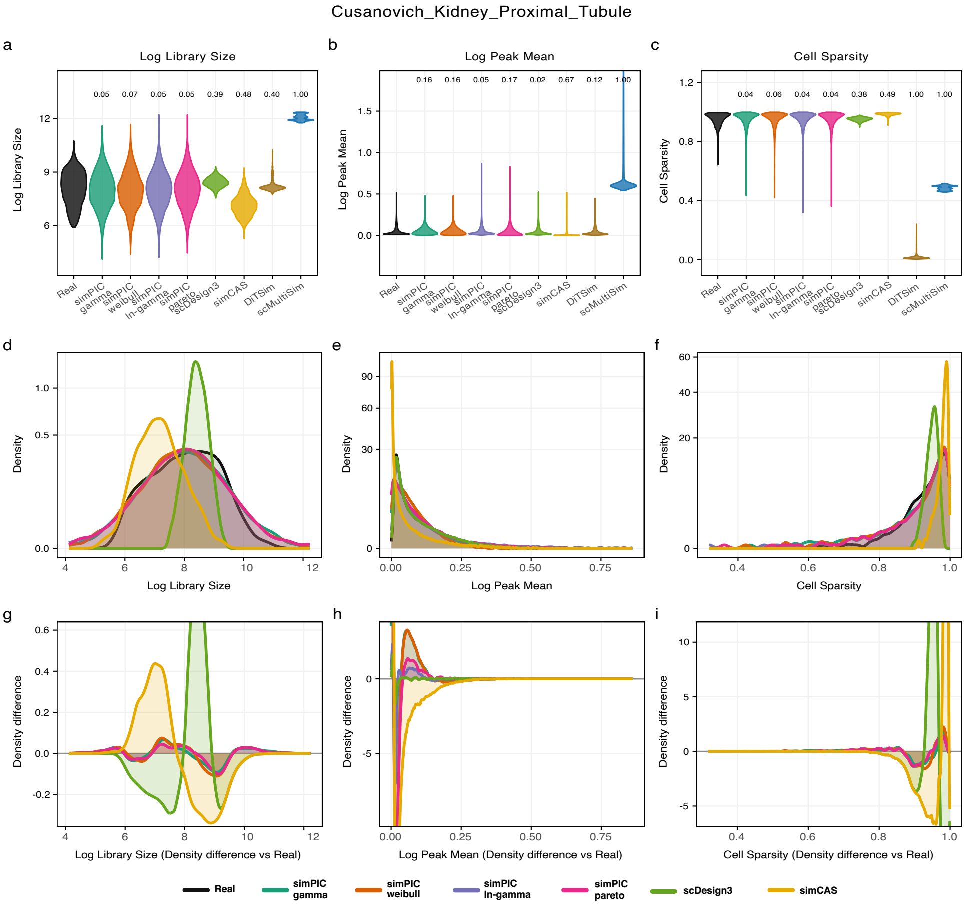
Cusanovich_Kidney_Proximal_Tubule
a
Log Library Size
b
Log Peak Mean
c
Cell Sparsity
d
e
f
g
h
i
Log Library Size
Log Peak Mean
Cell Sparsity
Density
Density difference
Log Library Size (Density difference vs Real)
Log Peak Mean (Density difference vs Real)
Cell Sparsity (Density difference vs Real)
Real
simPIC gamma
simPIC weibull
simPIC ln-gamma
simPIC pareto
scDesign3
simCAS
DiTSim
scMultiSim

Cusanovich_Kidney_Proximal_Tubule

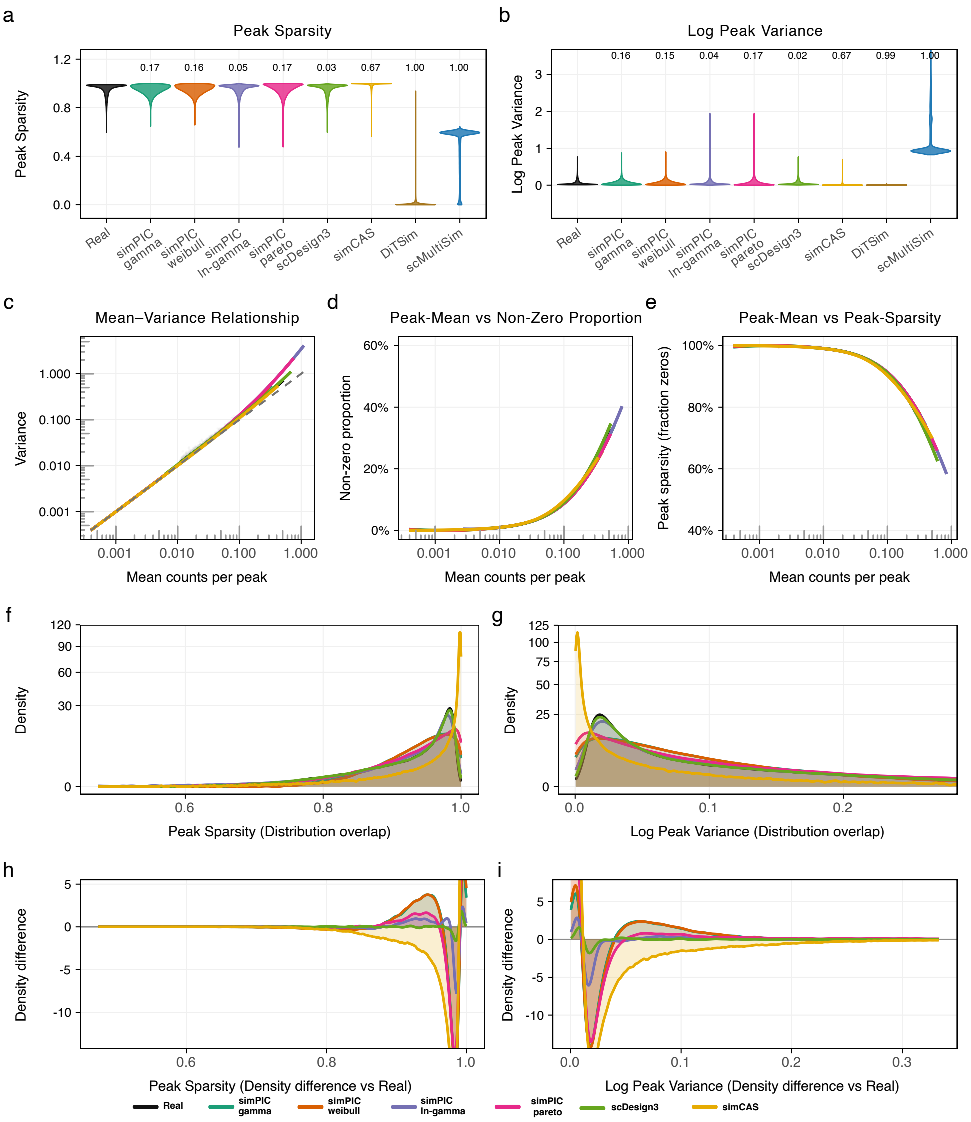

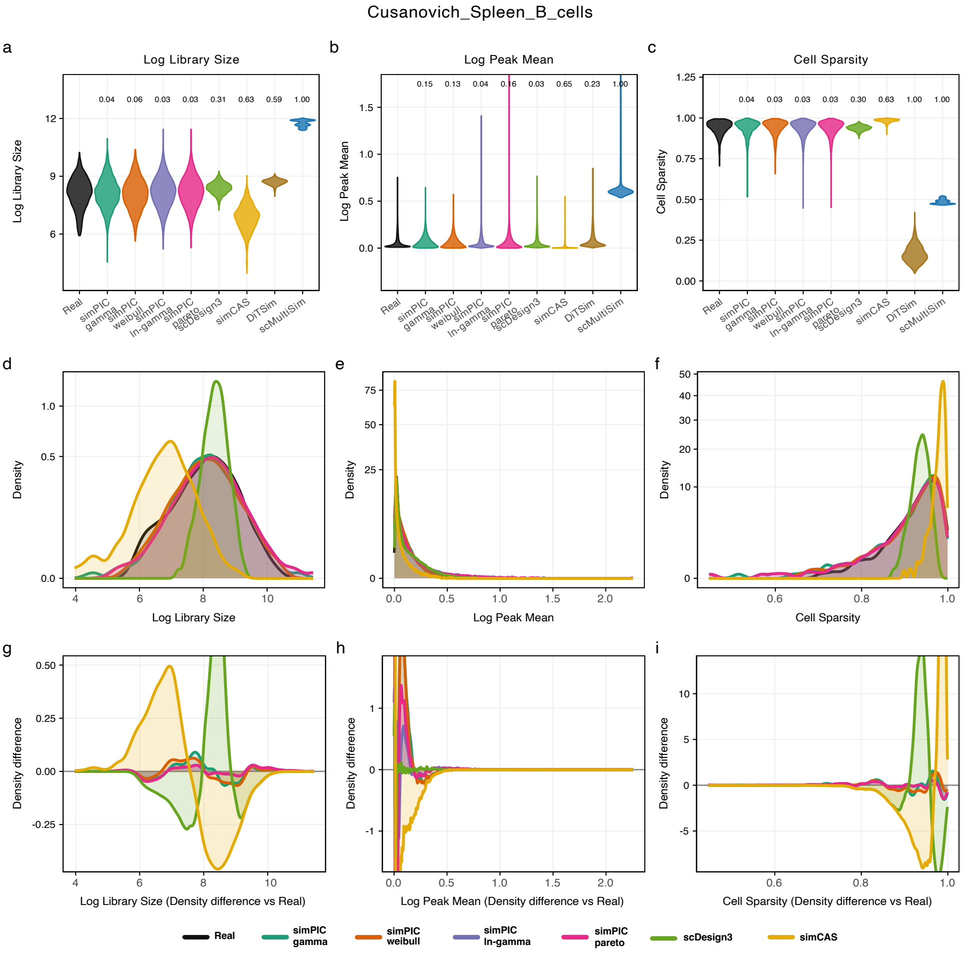

Cusanovich_Spleen_B_cells
a
Log Library Size
b
Log Peak Mean
c
Cell Sparsity
d
e
f
g
h
i
Log Library Size (Density difference vs Real)
Log Peak Mean (Density difference vs Real)
Cell Sparsity (Density difference vs Real)
Real
simPIC gamma
simPIC weibull
simPIC ln-gamma
simPIC pareto
scDesign3
simCAS
DiTSim
scMultiSim

Cusanovich_Spleen_B_cells

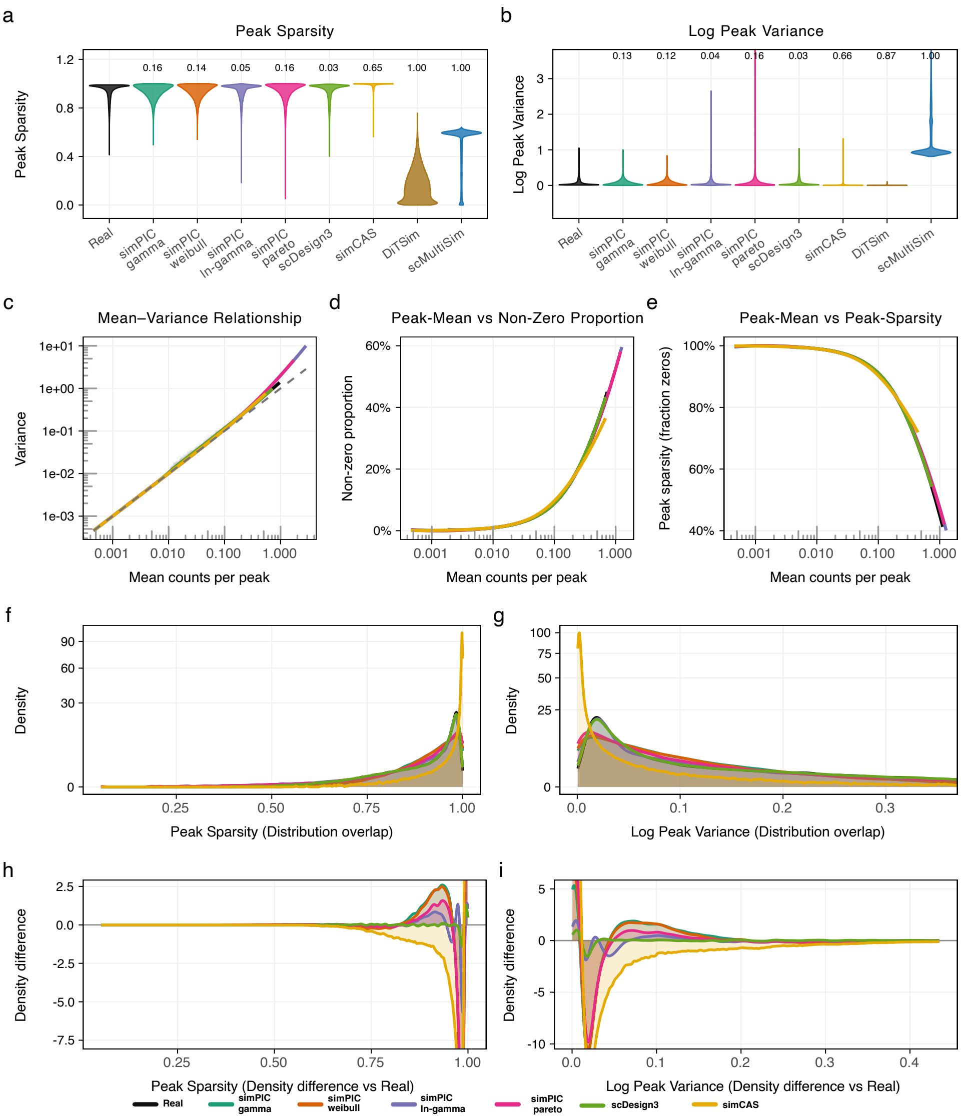

Fly_Brain_Capa

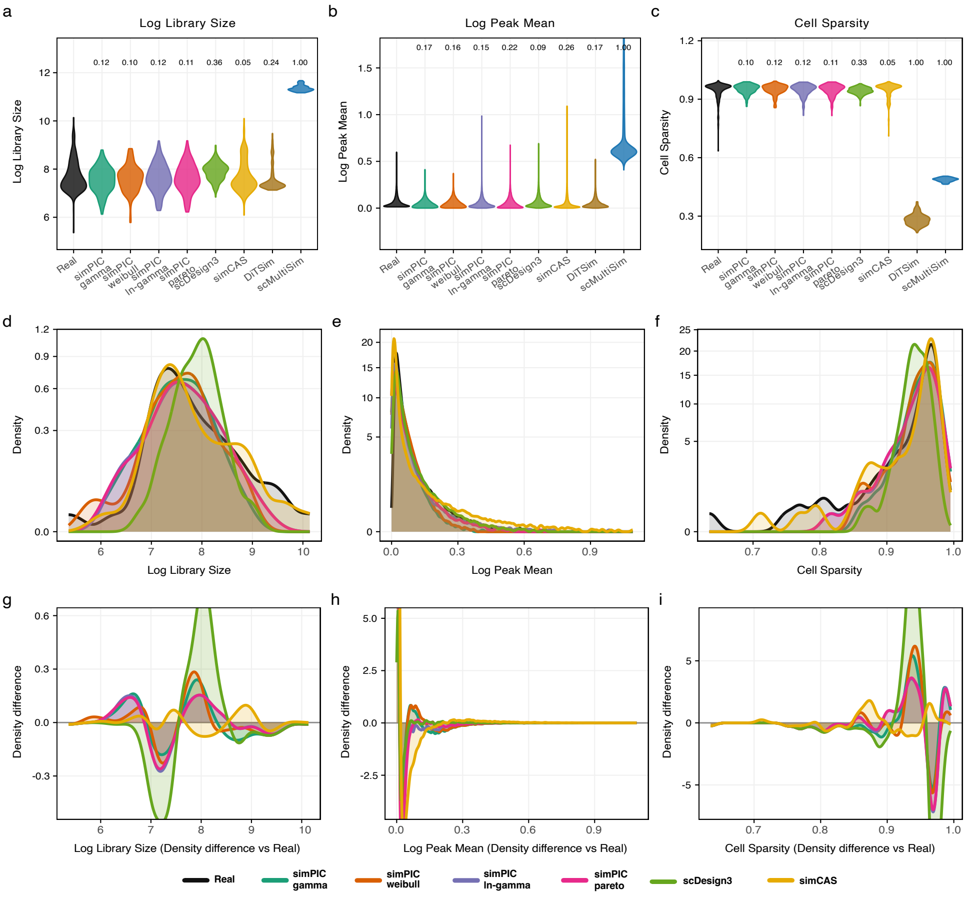

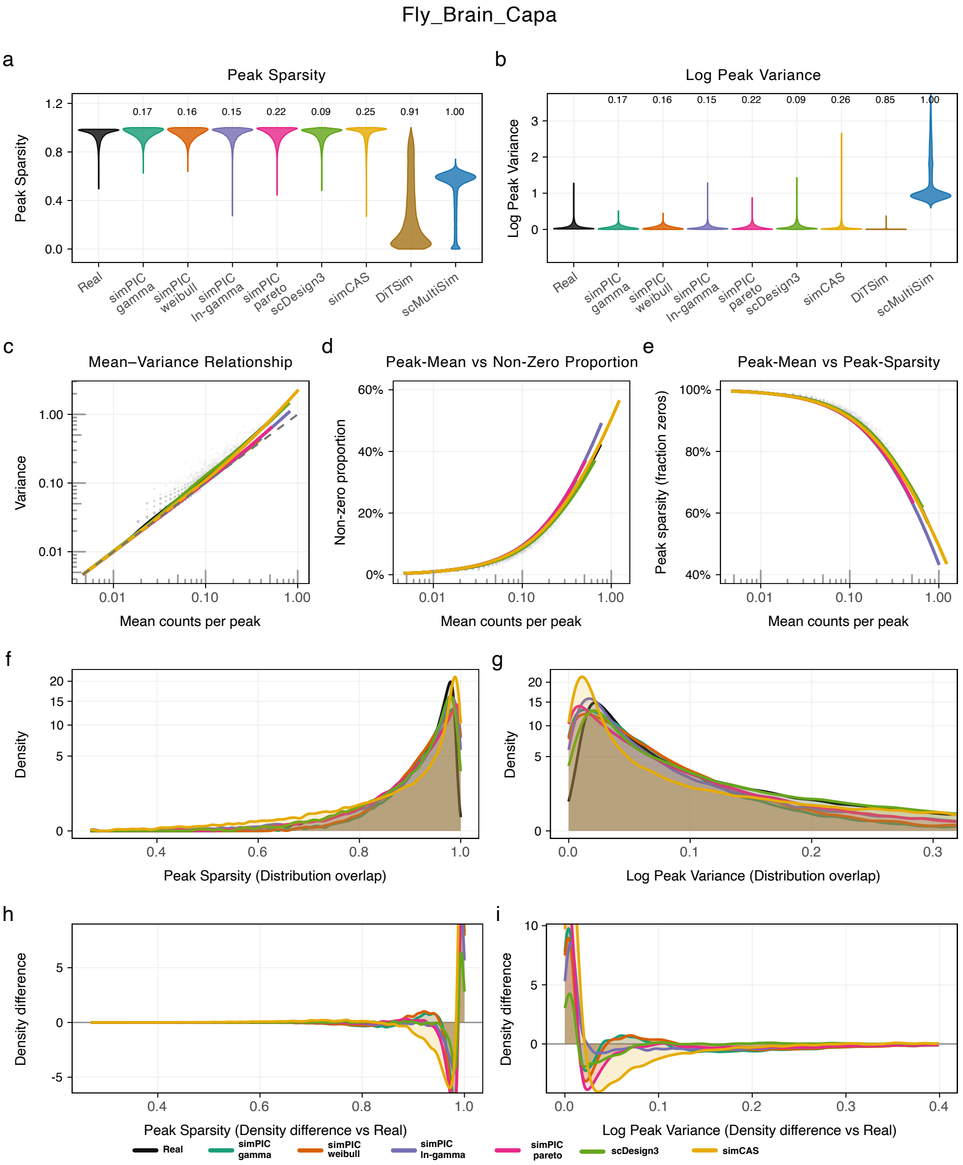
Fly_Brain_Capa
a
Peak Sparsity
Peak Sparsity
0.17
0.16
0.15
0.22
0.09
0.25
0.91
1.00
Real
simPIC gamma
simPIC weibull
simPIC ln-gamma
simPIC pareto
scDesign3
simCAS
DiTSim
scMultiSim
b
Log Peak Variance
Log Peak Variance
0.17
0.16
0.15
0.22
0.09
0.26
0.85
1.00
c
Mean–Variance Relationship
Variance
Mean counts per peak
d
Peak-Mean vs Non-Zero Proportion
Non-zero proportion
Mean counts per peak
e
Peak-Mean vs Peak-Sparsity
Peak sparsity (fraction zeros)
Mean counts per peak
f
Density
Peak Sparsity (Distribution overlap)
g
Density
Log Peak Variance (Distribution overlap)
h
Density difference
Peak Sparsity (Density difference vs Real)
i
Density difference
Log Peak Variance (Density difference vs Real)
Real
simPIC gamma
simPIC weibull
simPIC ln-gamma
simPIC pareto
scDesign3
simCAS

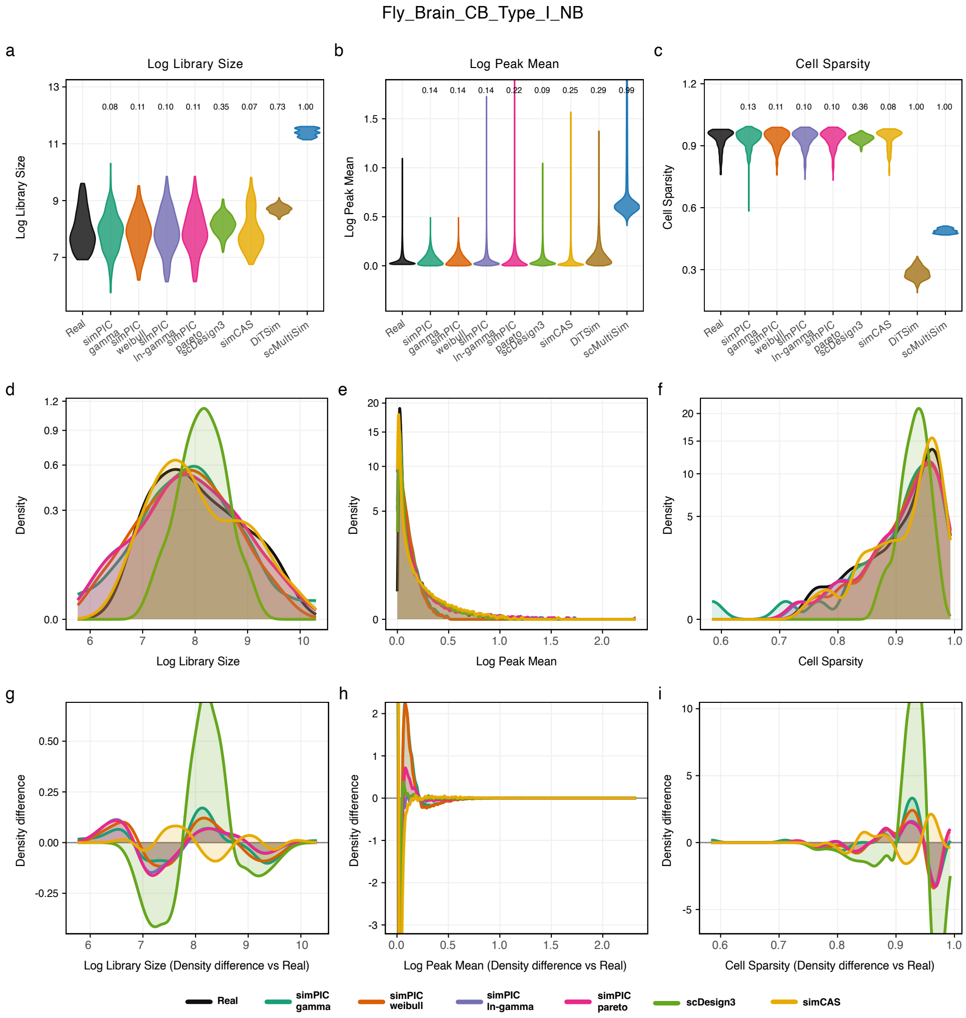

Fly_Brain_CB_Type_I_NB
a
Log Library Size
b
Log Peak Mean
c
Cell Sparsity
Real
simPIC gamma
simPIC weibull
simPIC ln-gamma
simPIC pareto
scDesign3
simCAS
DiTSim
scMultiSim
d
e
f
Density
Log Library Size
Log Peak Mean
Cell Sparsity
g
h
i
Density difference
Log Library Size (Density difference vs Real)
Log Peak Mean (Density difference vs Real)
Cell Sparsity (Density difference vs Real)

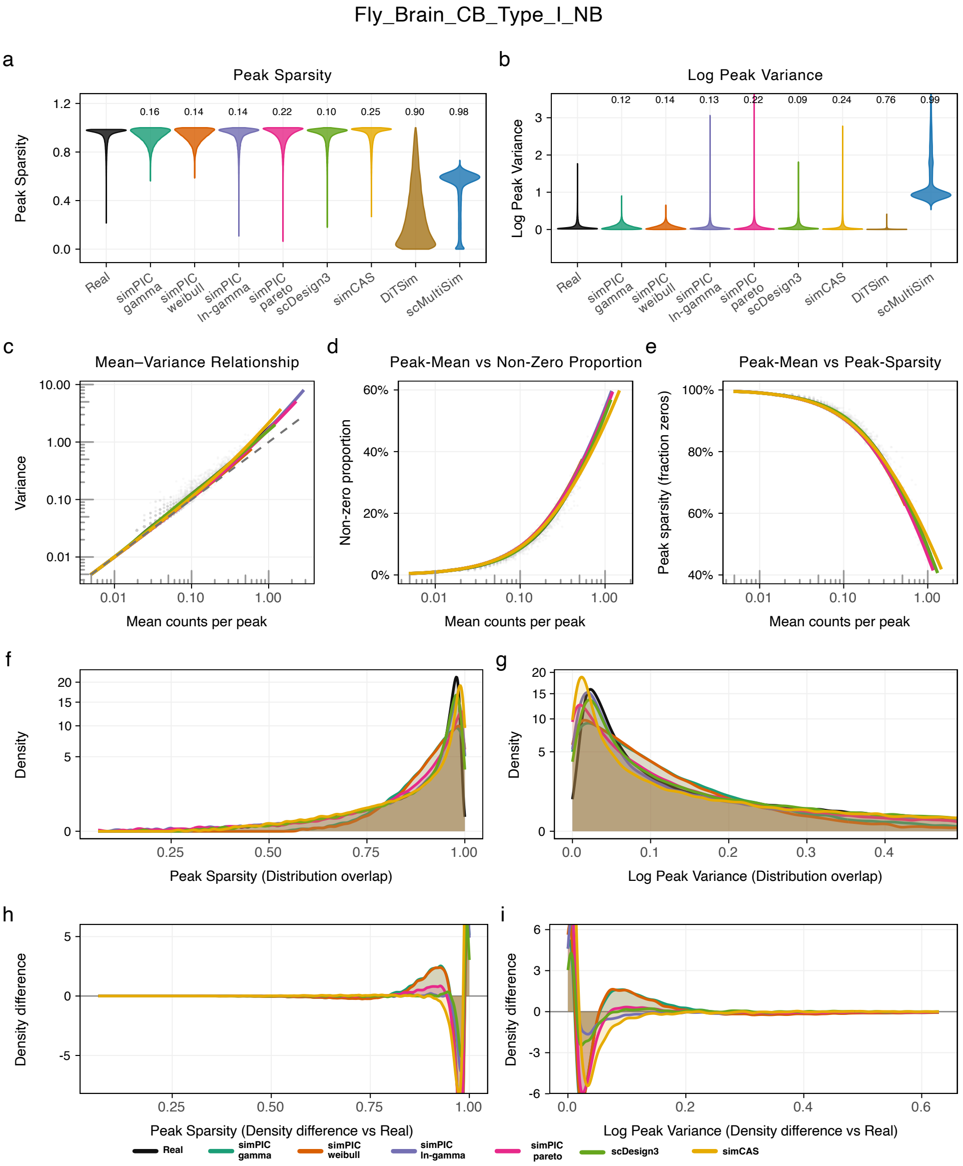
Fly_Brain_CB_Type_I_NB
a
Peak Sparsity
Peak Sparsity
0.16 0.14 0.14 0.22 0.10 0.25 0.90 0.98
Real
simPIC gamma
simPIC weibull
simPIC ln-gamma
simPIC pareto
scDesign3
simCAS
DiTSim
scMultiSim
b
Log Peak Variance
Log Peak Variance
0.12 0.14 0.13 0.22 0.09 0.24 0.76 0.99
c
Mean–Variance Relationship
Variance
Mean counts per peak
d
Peak-Mean vs Non-Zero Proportion
Non-zero proportion
Mean counts per peak
e
Peak-Mean vs Peak-Sparsity
Peak sparsity (fraction zeros)
Mean counts per peak
f
Density
Peak Sparsity (Distribution overlap)
g
Density
Log Peak Variance (Distribution overlap)
h
Density difference
Peak Sparsity (Density difference vs Real)
i
Density difference
Log Peak Variance (Density difference vs Real)
Real
simPIC gamma
simPIC weibull
simPIC ln-gamma
simPIC pareto
scDesign3
simCAS

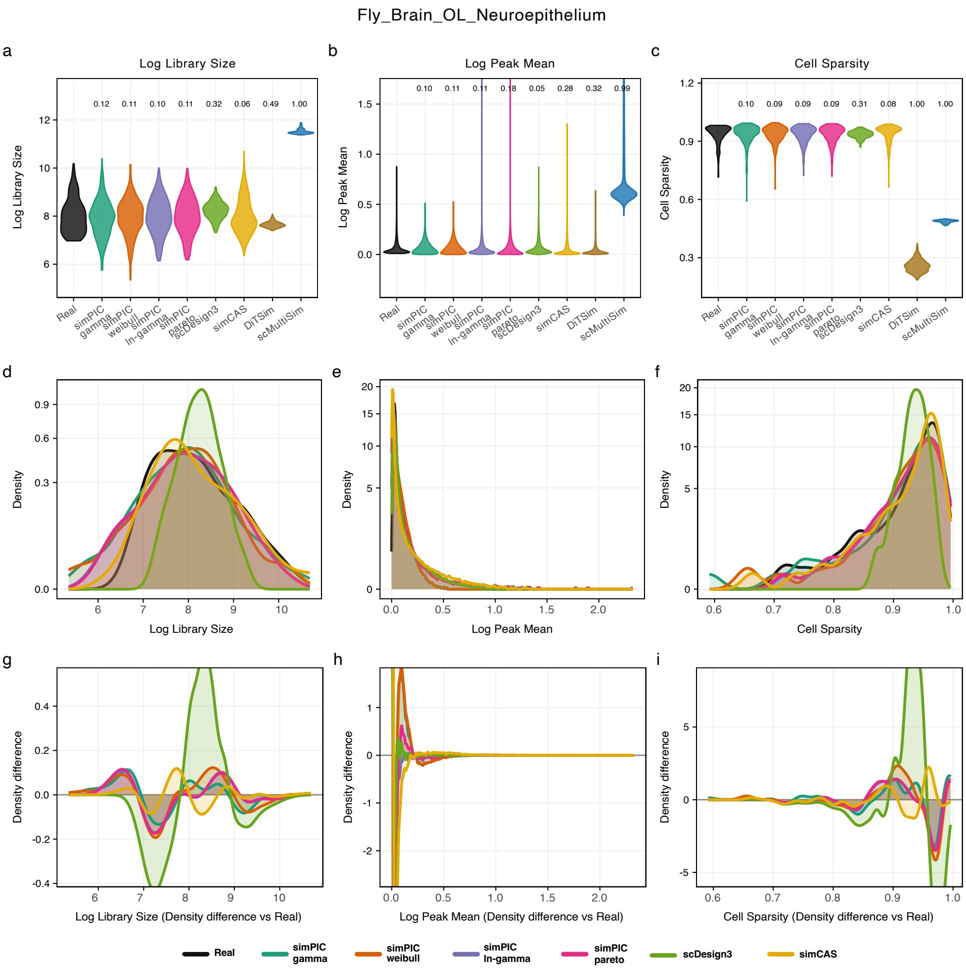

Fly_Brain_OL_Neuroepithelium
a
Log Library Size
b
Log Peak Mean
c
Cell Sparsity
d
e
f
g
h
i
Log Library Size
Log Peak Mean
Cell Sparsity
Density
Log Library Size (Density difference vs Real)
Log Peak Mean (Density difference vs Real)
Cell Sparsity (Density difference vs Real)
Density difference
Real
simPIC gamma
simPIC weibull
simPIC ln-gamma
simPIC pareto
scDesign3
simCAS
DiTSim
scMultiSim

Fly_Brain_OL_Neuroepithelium

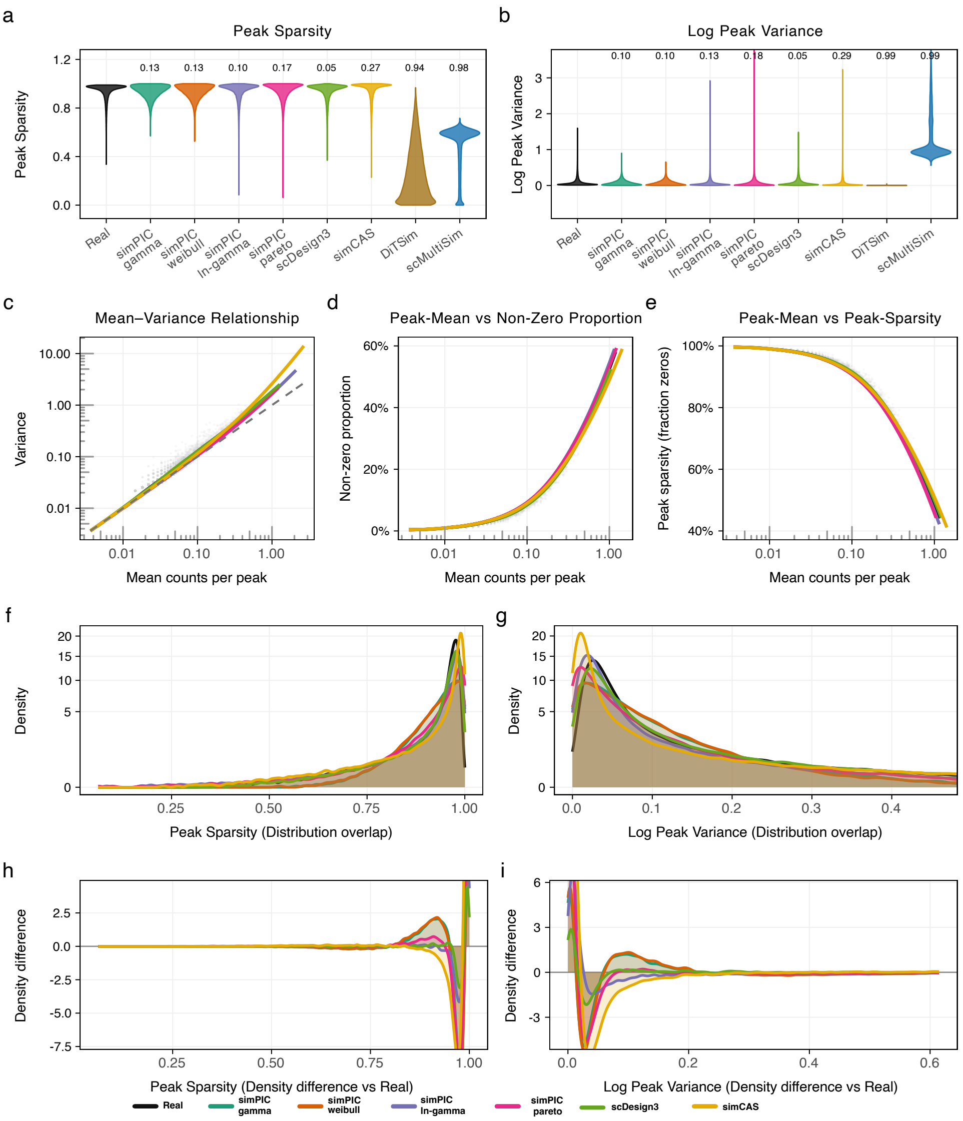

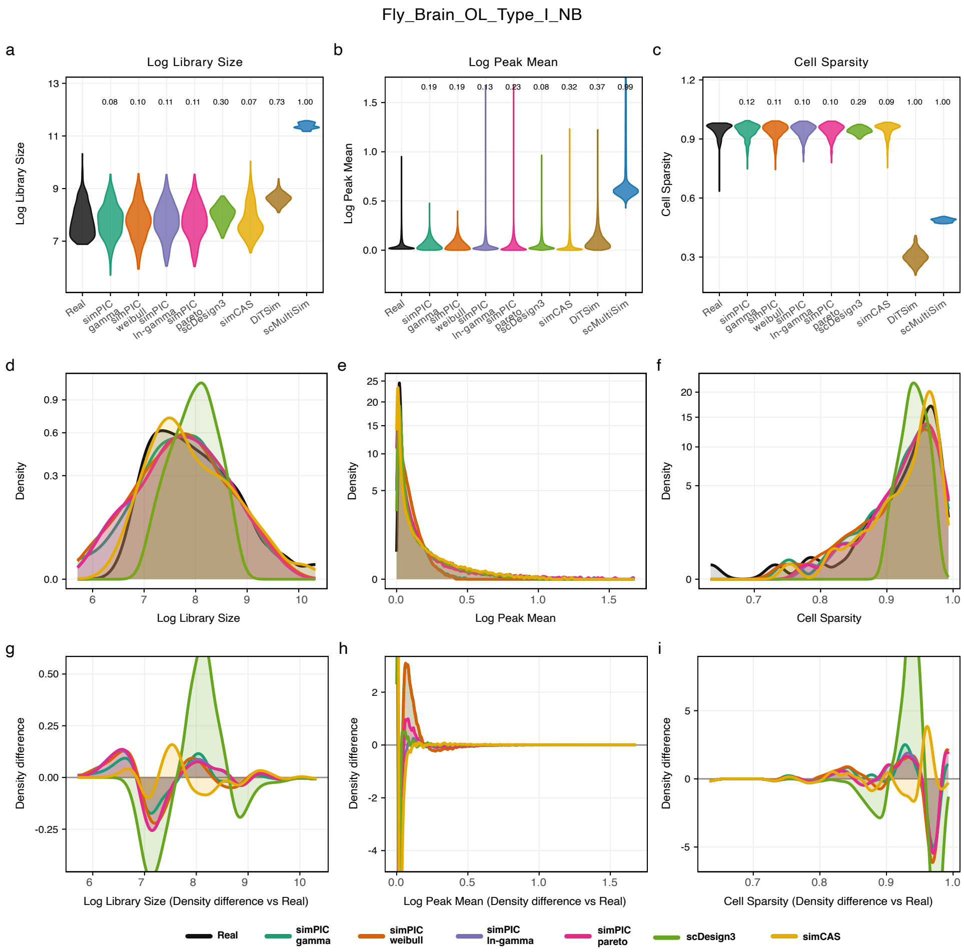
Fly_Brain_OL_Type_I_NB
a
Log Library Size
Log Library Size
0.08 0.10 0.11 0.11 0.30 0.07 0.73 1.00
b
Log Peak Mean
Log Peak Mean
0.19 0.19 0.13 0.23 0.08 0.32 0.37 0.99
c
Cell Sparsity
Cell Sparsity
0.12 0.11 0.10 0.10 0.29 0.09 1.00 1.00
Real
simPIC gamma
simPIC weibull
simPIC ln-gamma
simPIC pareto
scDesign3
simCAS
DiTSim
scMultiSim
d
e
f
Density
Log Library Size
Log Peak Mean
Cell Sparsity
g
h
i
Density difference
Log Library Size (Density difference vs Real)
Log Peak Mean (Density difference vs Real)
Cell Sparsity (Density difference vs Real)
Real
simPIC gamma
simPIC weibull
simPIC ln-gamma
simPIC pareto
scDesign3
simCAS

Fly_Brain_OL_Type_I_NB

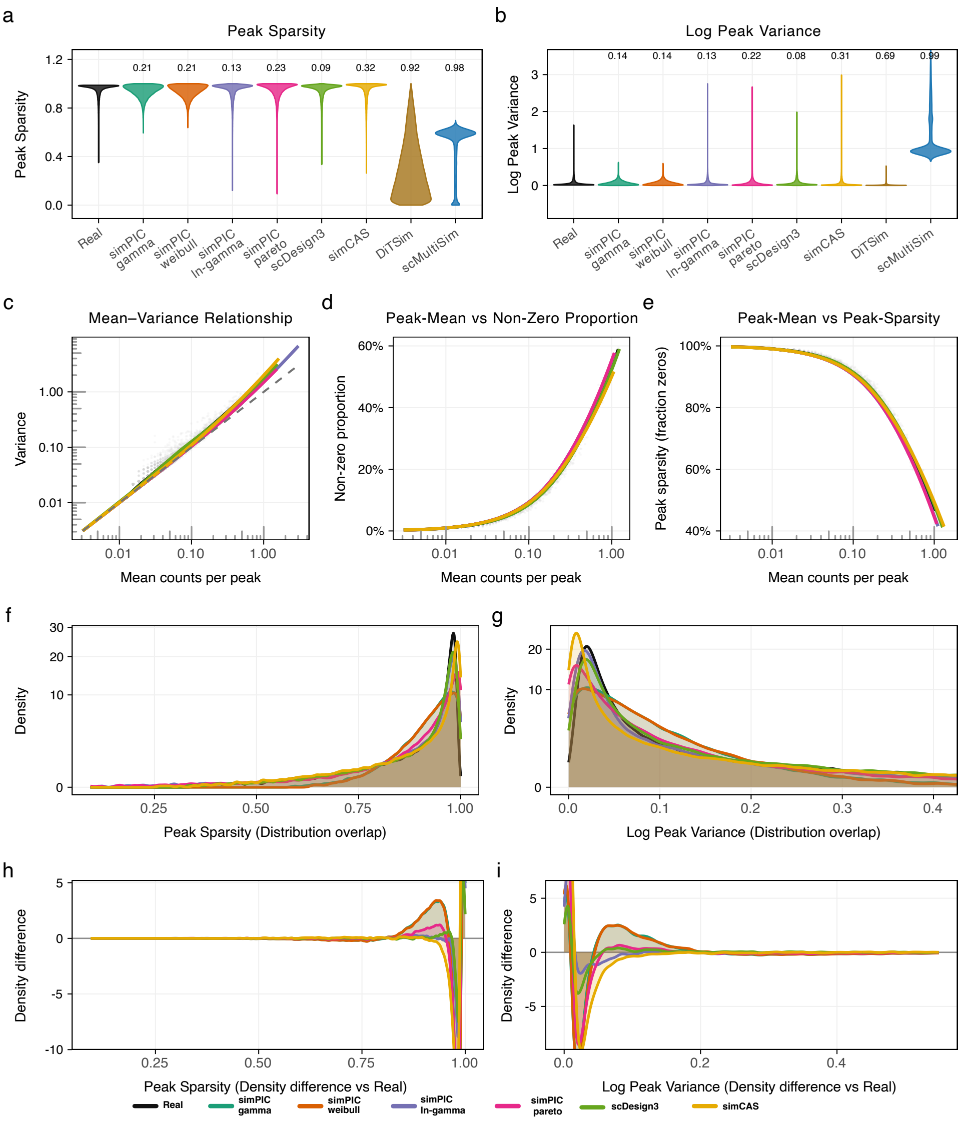

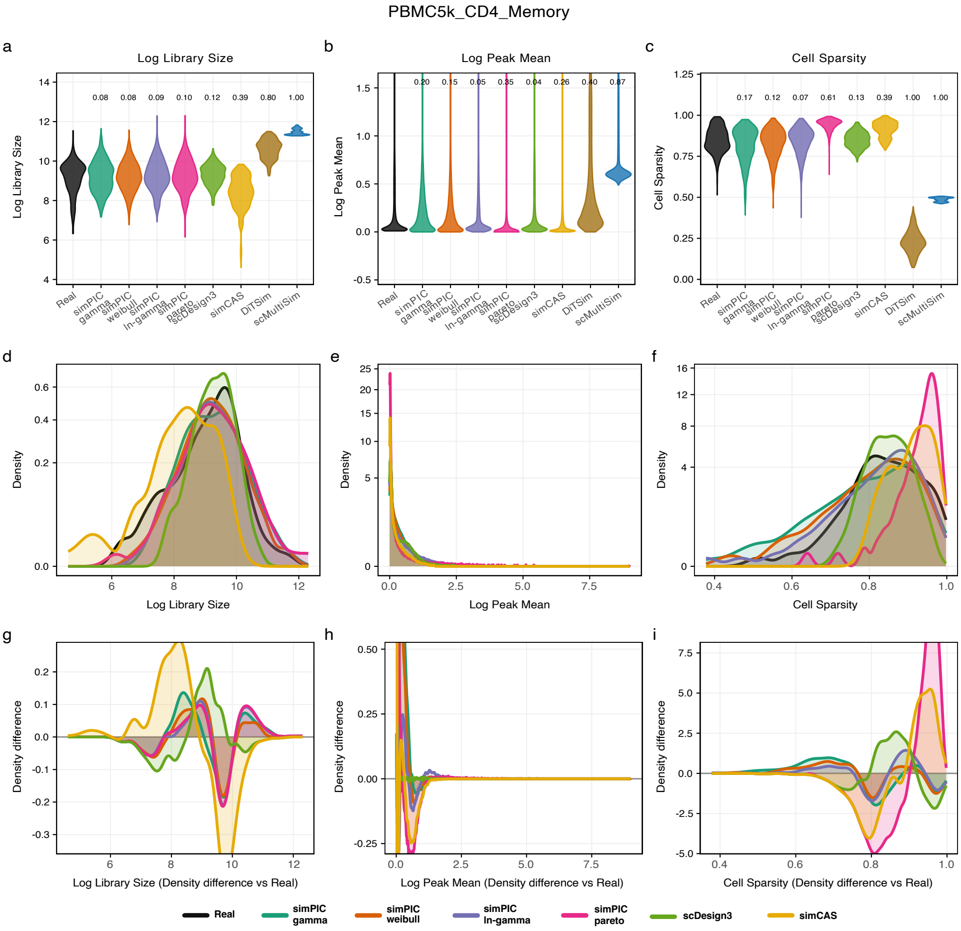
PBMC5k_CD4_Memory
a
Log Library Size
b
Log Peak Mean
c
Cell Sparsity
d
e
f
g
h
i
Log Library Size
Log Peak Mean
Cell Sparsity
Density
Log Library Size (Density difference vs Real)
Log Peak Mean (Density difference vs Real)
Cell Sparsity (Density difference vs Real)
Density difference
Real
simPIC gamma
simPIC weibull
simPIC ln-gamma
simPIC pareto
scDesign3
simCAS
DiTSim
scMultiSim

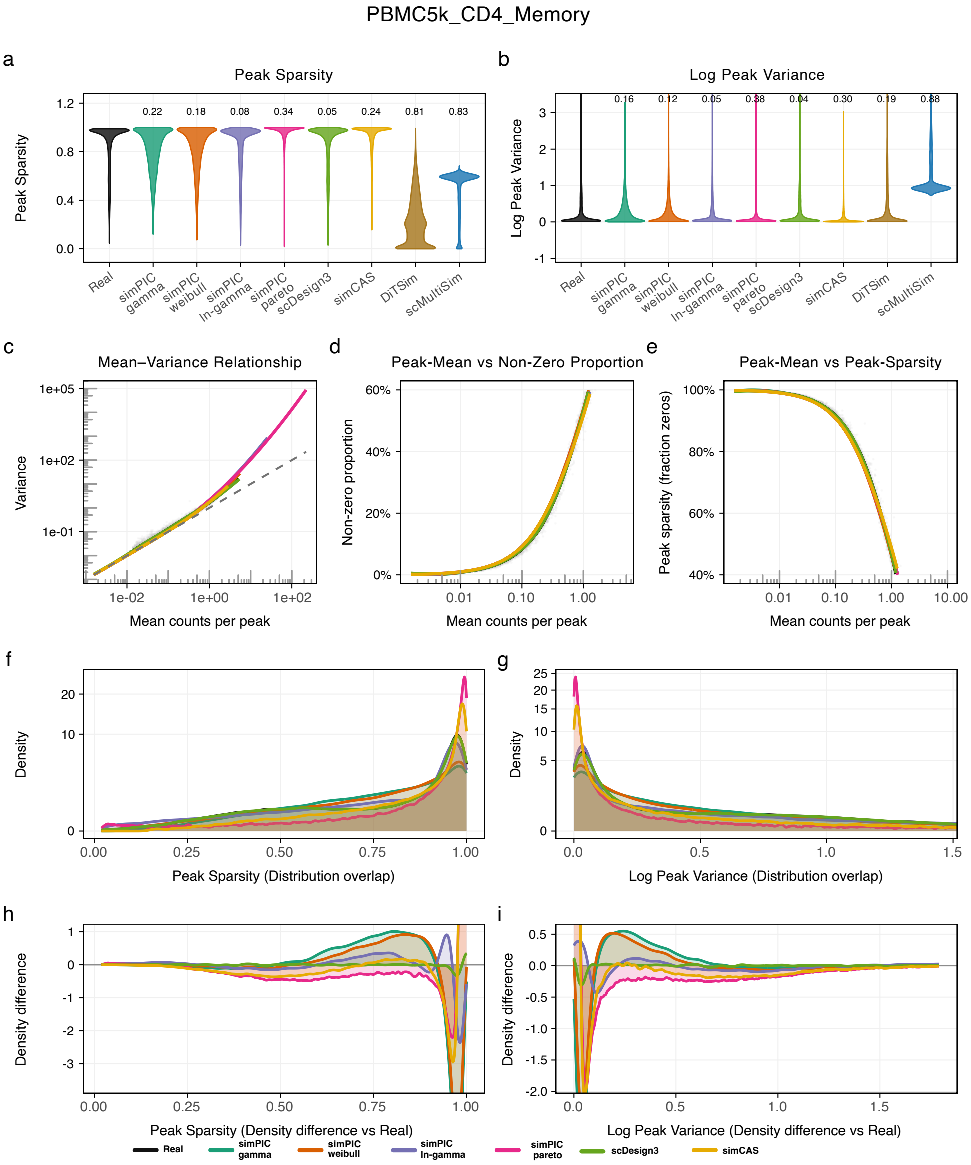
PBMC5k_CD4_Memory
a
Peak Sparsity
Peak Sparsity
0.22
0.18
0.08
0.34
0.05
0.24
0.81
0.83
Real
simPIC gamma
simPIC weibull
simPIC ln-gamma
simPIC pareto
scDesign3
simCAS
DiTSim
scMultiSim
b
Log Peak Variance
Log Peak Variance
0.16
0.12
0.05
0.38
0.04
0.30
0.19
0.88
c
Mean–Variance Relationship
Variance
Mean counts per peak
d
Peak-Mean vs Non-Zero Proportion
Non-zero proportion
Mean counts per peak
e
Peak-Mean vs Peak-Sparsity
Peak sparsity (fraction zeros)
Mean counts per peak
f
Density
Peak Sparsity (Distribution overlap)
g
Density
Log Peak Variance (Distribution overlap)
h
Density difference
Peak Sparsity (Density difference vs Real)
i
Density difference
Log Peak Variance (Density difference vs Real)
Real
simPIC gamma
simPIC weibull
simPIC ln-gamma
simPIC pareto
scDesign3
simCAS

PBMC5k_CD4_Naive

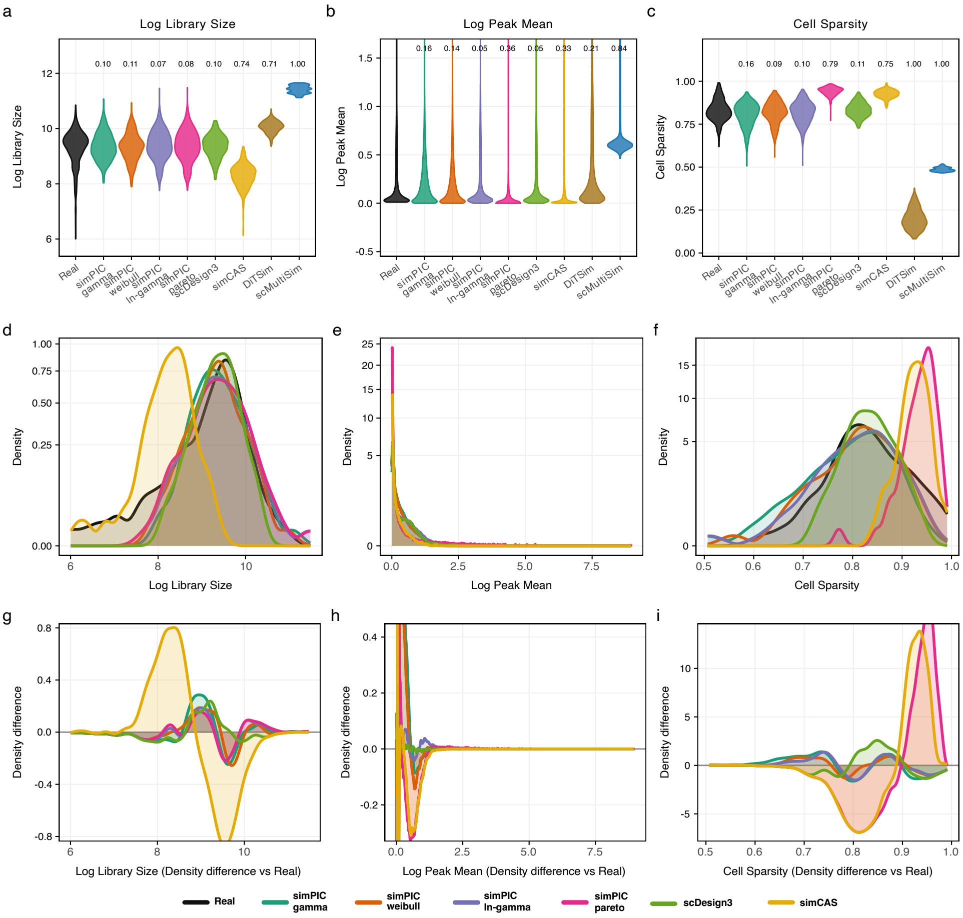

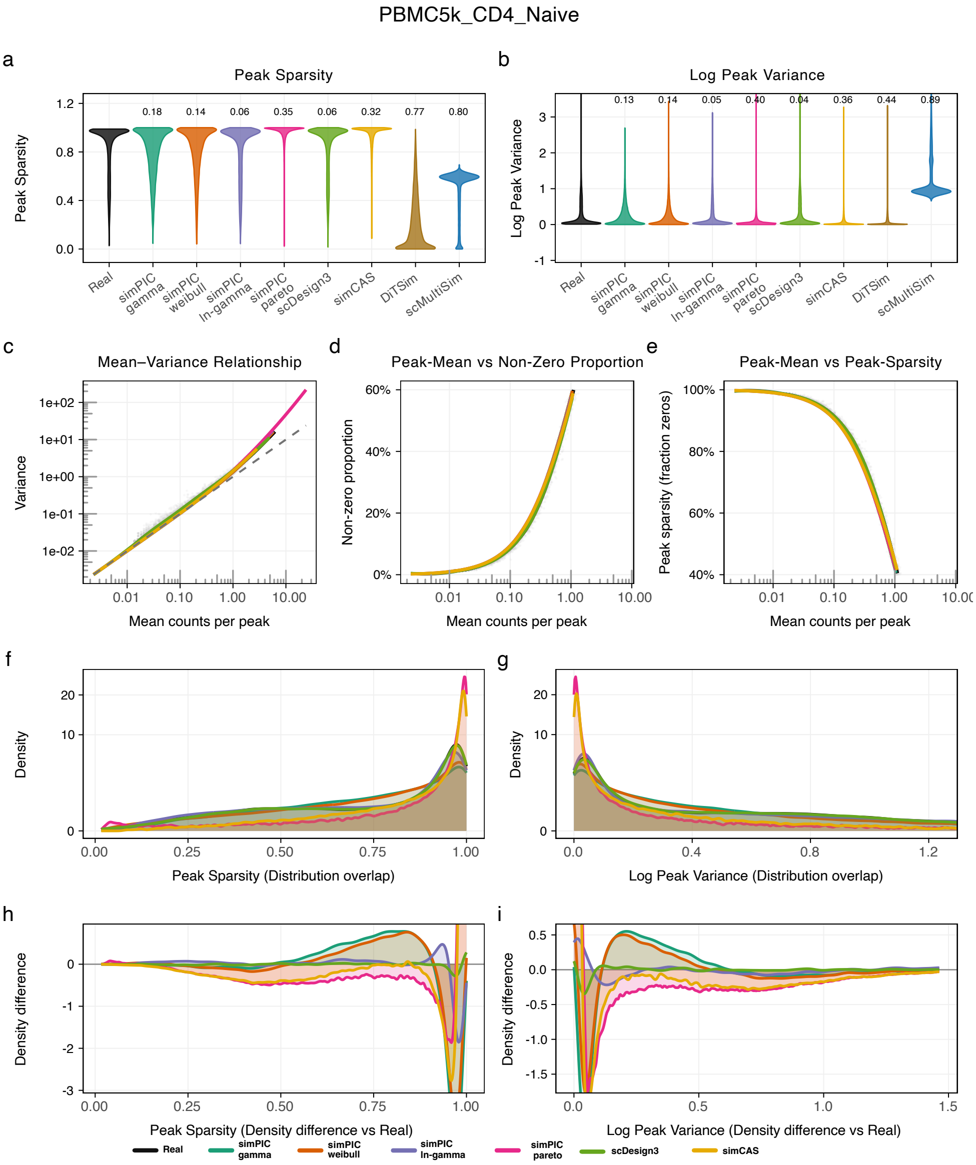
PBMC5k_CD4_Naive
a
Peak Sparsity
Peak Sparsity
0.18 0.14 0.06 0.35 0.06 0.32 0.77 0.80
Real
simPIC gamma
simPIC weibull
simPIC ln-gamma
simPIC pareto
scDesign3
simCAS
DiTSim
scMultiSim
b
Log Peak Variance
Log Peak Variance
0.13 0.14 0.05 0.40 0.04 0.36 0.44 0.89
c
Mean–Variance Relationship
Variance
Mean counts per peak
d
Peak-Mean vs Non-Zero Proportion
Non-zero proportion
Mean counts per peak
e
Peak-Mean vs Peak-Sparsity
Peak sparsity (fraction zeros)
Mean counts per peak
f
Density
Peak Sparsity (Distribution overlap)
g
Density
Log Peak Variance (Distribution overlap)
h
Density difference
Peak Sparsity (Density difference vs Real)
i
Density difference
Log Peak Variance (Density difference vs Real)
Real
simPIC gamma
simPIC weibull
simPIC ln-gamma
simPIC pareto
scDesign3
simCAS

PBMC5k_CD8_Effector

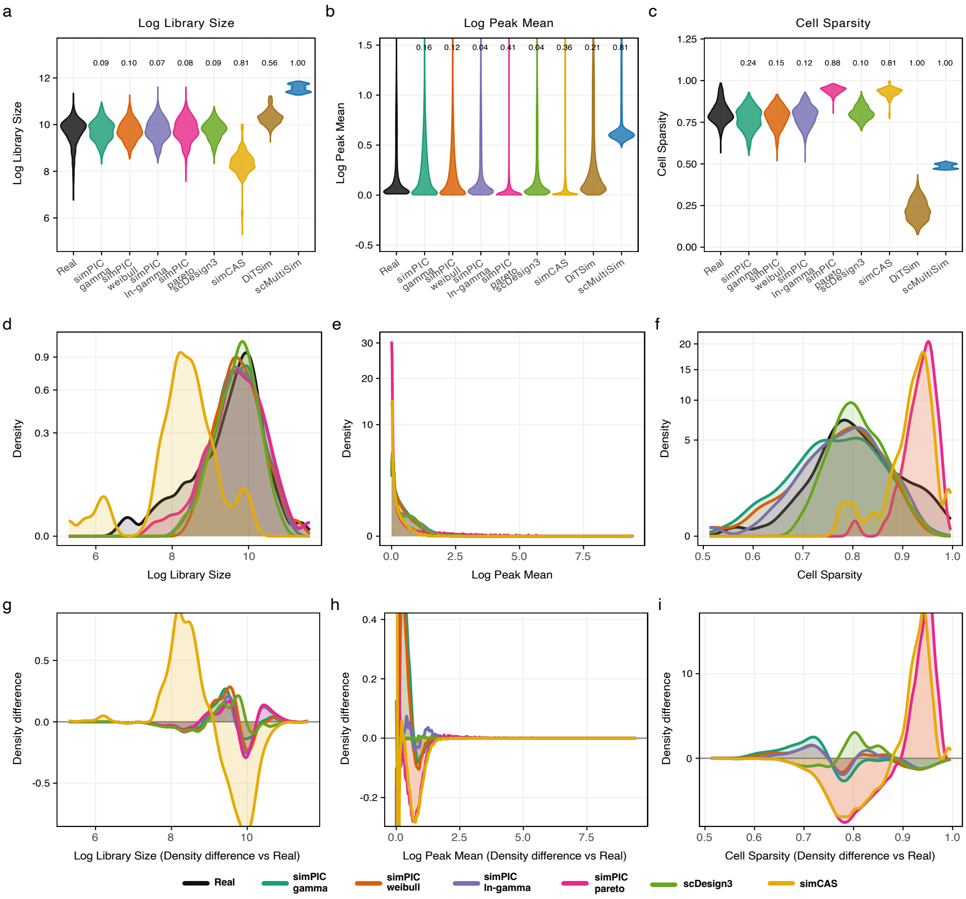

PBMC5k_CD8_Effector

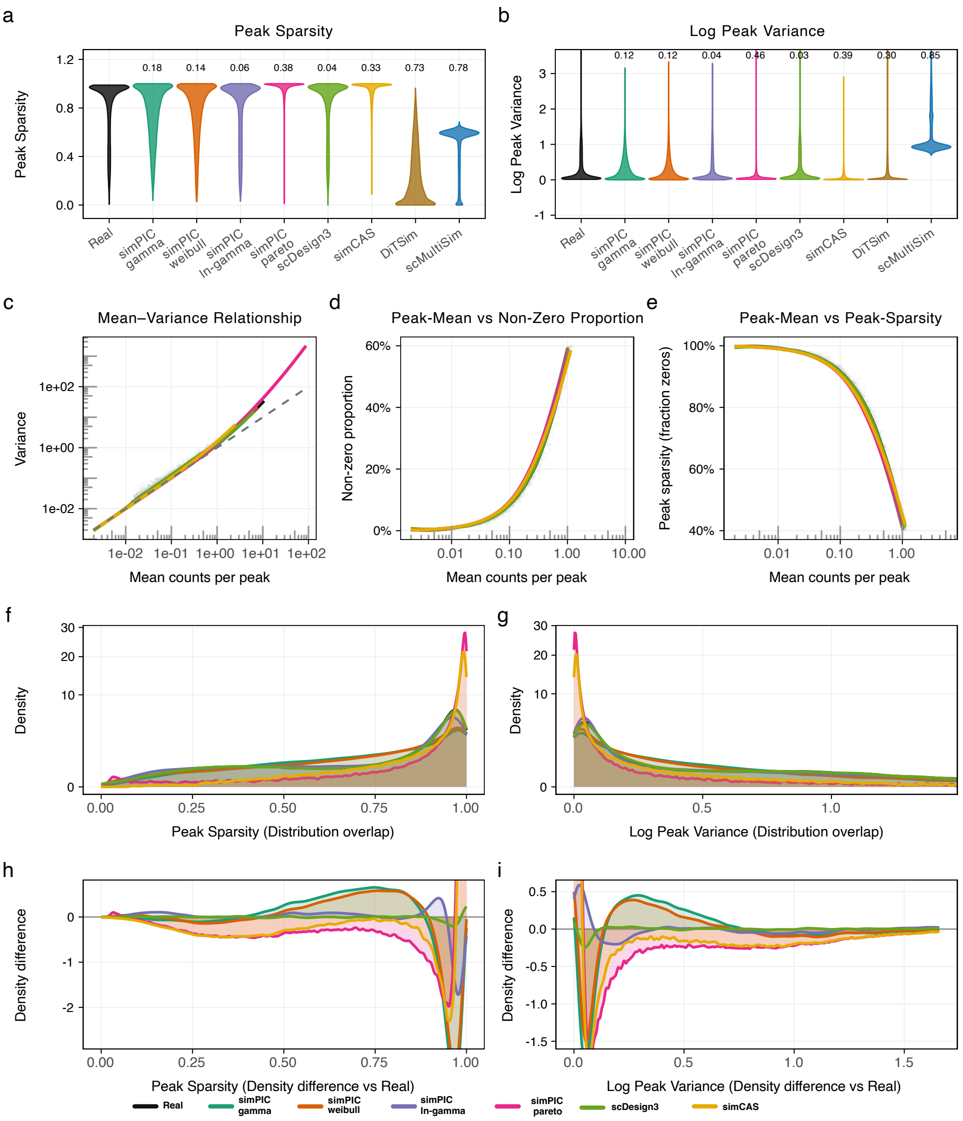

PBMC5k_CD8_Naive

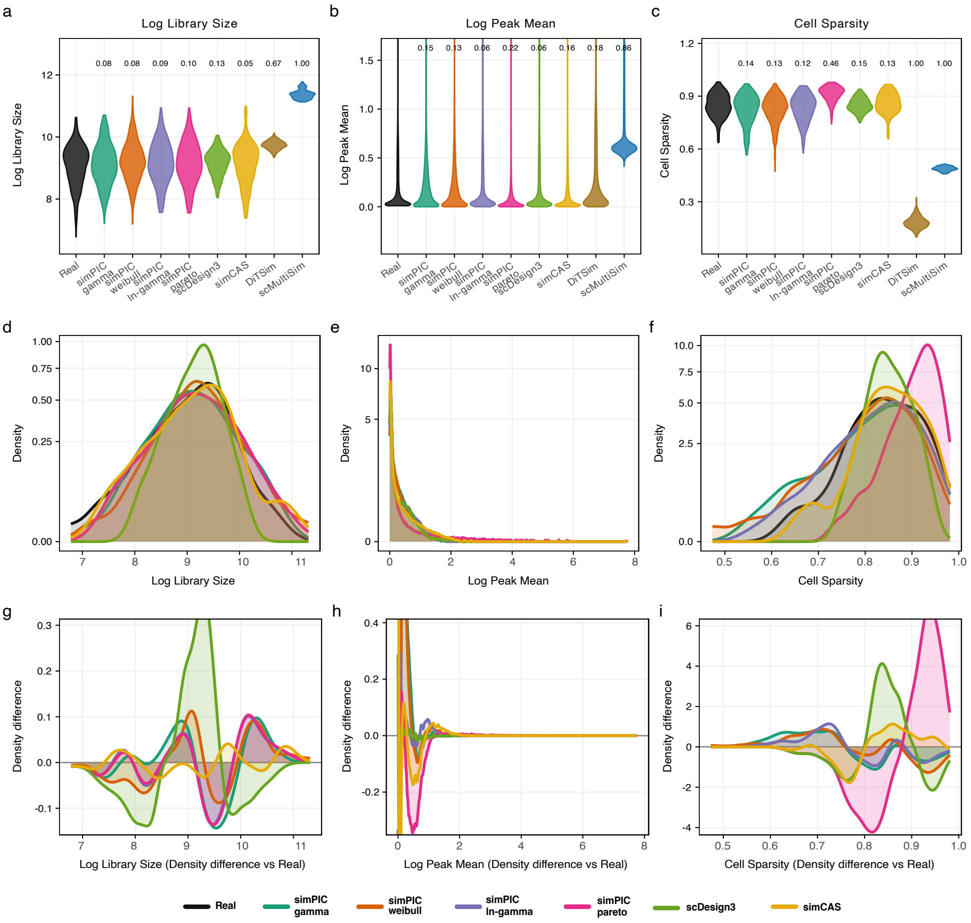

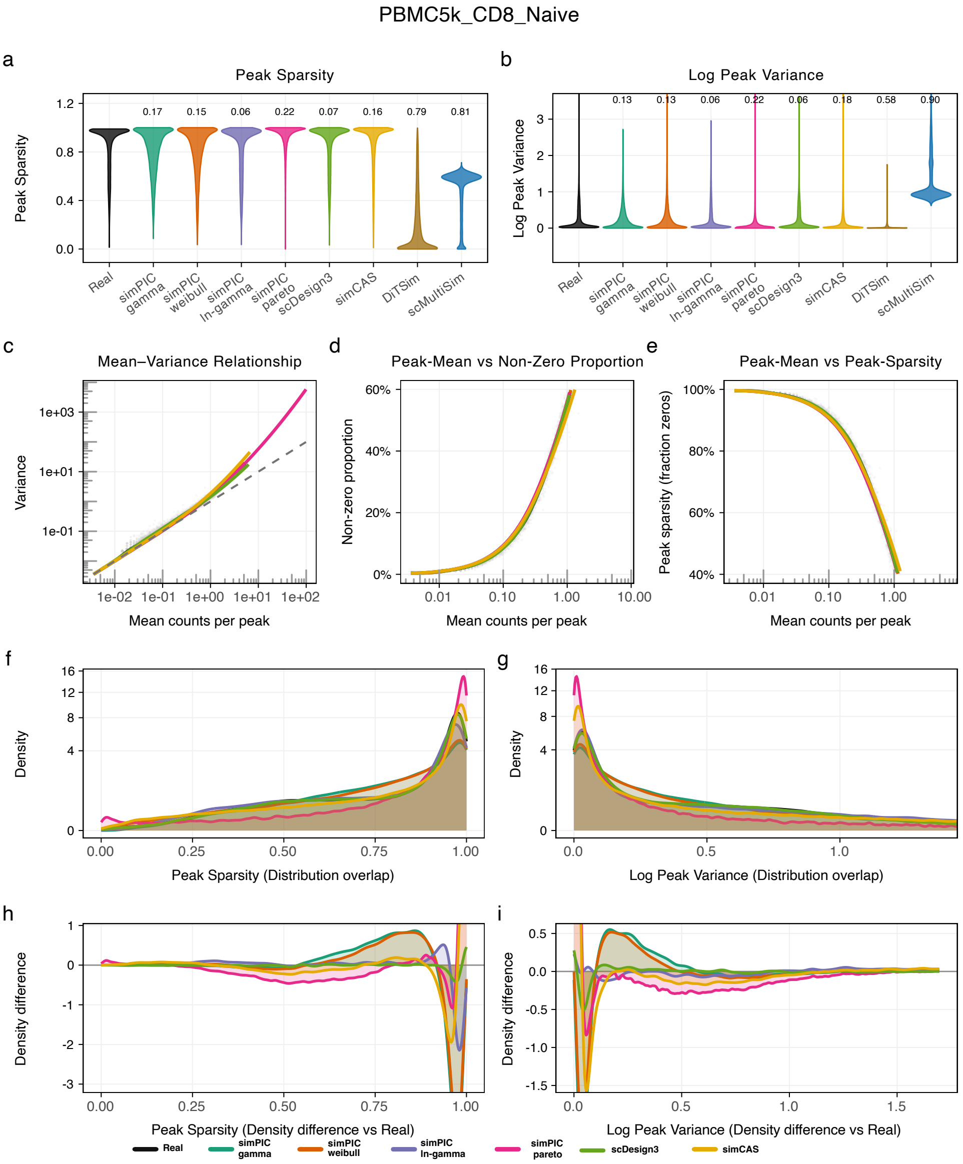
PBMC5k_CD8_Naive
a
Peak Sparsity
Peak Sparsity
0.17
0.15
0.06
0.22
0.07
0.16
0.79
0.81
Real
simPIC gamma
simPIC weibull
simPIC ln-gamma
simPIC pareto
scDesign3
simCAS
DiTSim
scMultiSim
b
Log Peak Variance
Log Peak Variance
0.13
0.13
0.06
0.22
0.06
0.18
0.58
0.90
c
Mean–Variance Relationship
Variance
Mean counts per peak
d
Peak-Mean vs Non-Zero Proportion
Non-zero proportion
Mean counts per peak
e
Peak-Mean vs Peak-Sparsity
Peak sparsity (fraction zeros)
Mean counts per peak
f
Density
Peak Sparsity (Distribution overlap)
g
Density
Log Peak Variance (Distribution overlap)
h
Density difference
Peak Sparsity (Density difference vs Real)
i
Density difference
Log Peak Variance (Density difference vs Real)
Real
simPIC gamma
simPIC weibull
simPIC ln-gamma
simPIC pareto
scDesign3
simCAS

PBMC5k_CD14_Monocytes

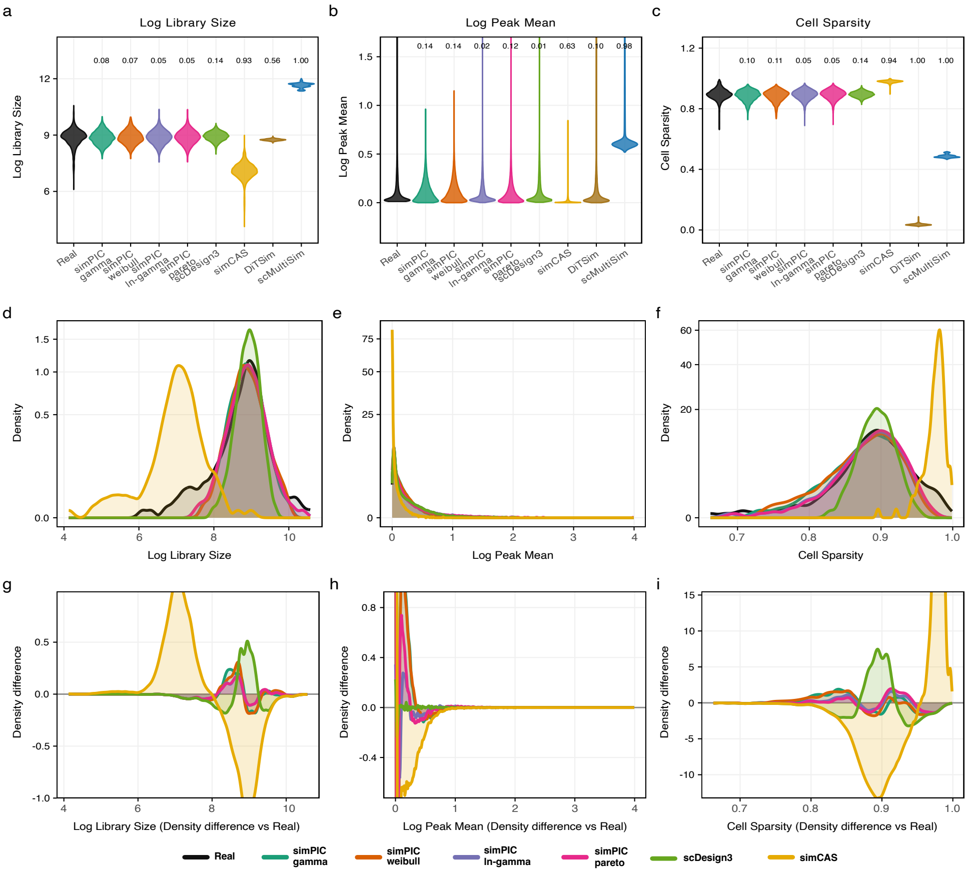

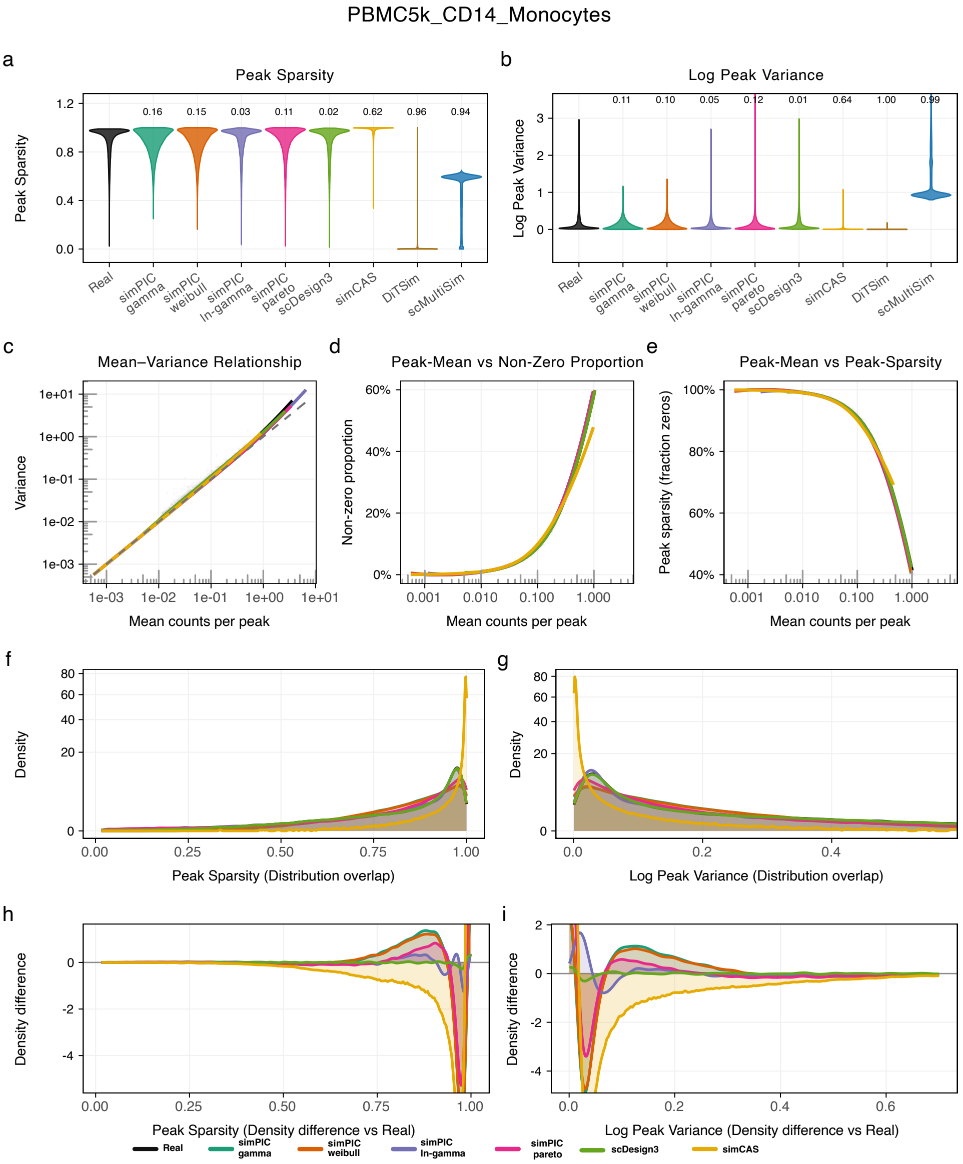
PBMC5k_CD14_Monocytes
a
Peak Sparsity
Peak Sparsity
0.16
0.15
0.03
0.11
0.02
0.62
0.96
0.94
Real
simPIC gamma
simPIC weibull
simPIC ln-gamma
simPIC pareto
scDesign3
simCAS
DiTSim
scMultiSim
b
Log Peak Variance
Log Peak Variance
0.11
0.10
0.05
0.12
0.01
0.64
1.00
0.99
c
Mean–Variance Relationship
Variance
Mean counts per peak
d
Peak-Mean vs Non-Zero Proportion
Non-zero proportion
Mean counts per peak
e
Peak-Mean vs Peak-Sparsity
Peak sparsity (fraction zeros)
Mean counts per peak
f
Density
Peak Sparsity (Distribution overlap)
g
Density
Log Peak Variance (Distribution overlap)
h
Density difference
Peak Sparsity (Density difference vs Real)
i
Density difference
Log Peak Variance (Density difference vs Real)
Real
simPIC gamma
simPIC weibull
simPIC ln-gamma
simPIC pareto
scDesign3
simCAS

PBMC5k_Double_Negative_T_cell

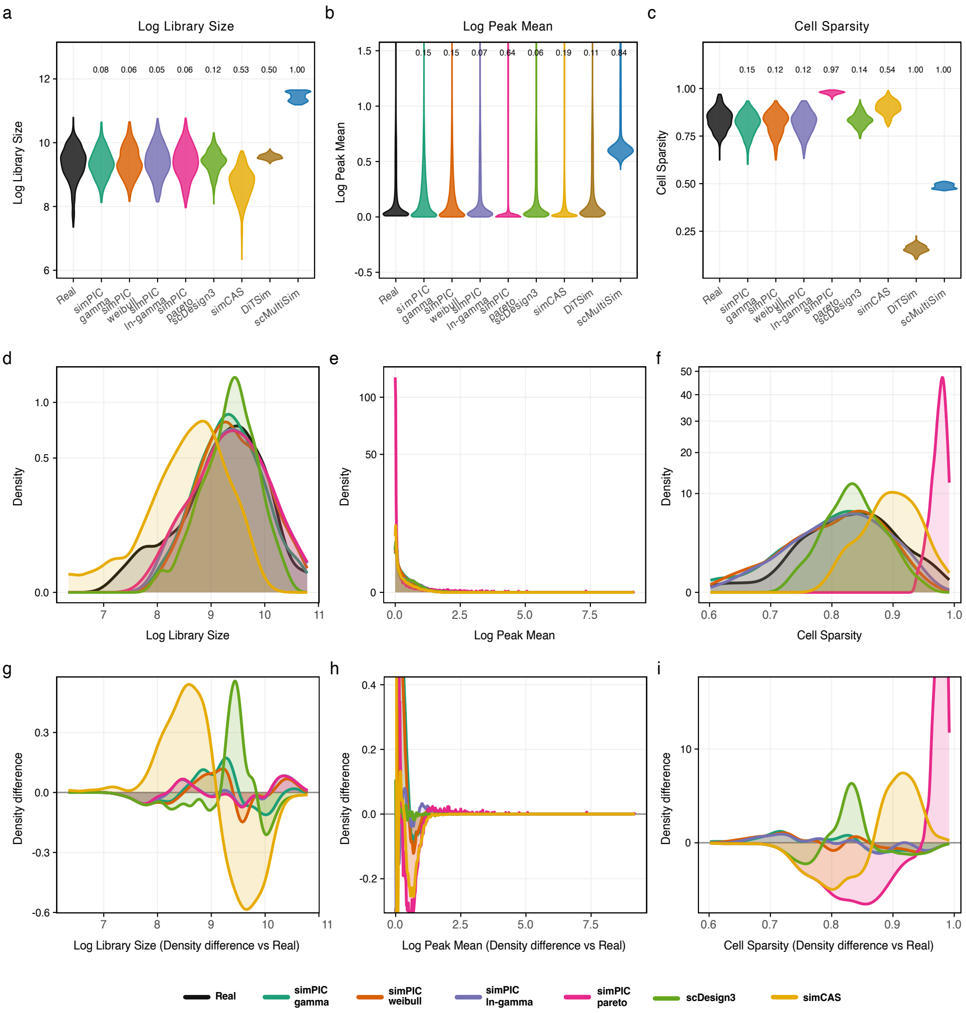

PBMC5k_Double_Negative_T_cell

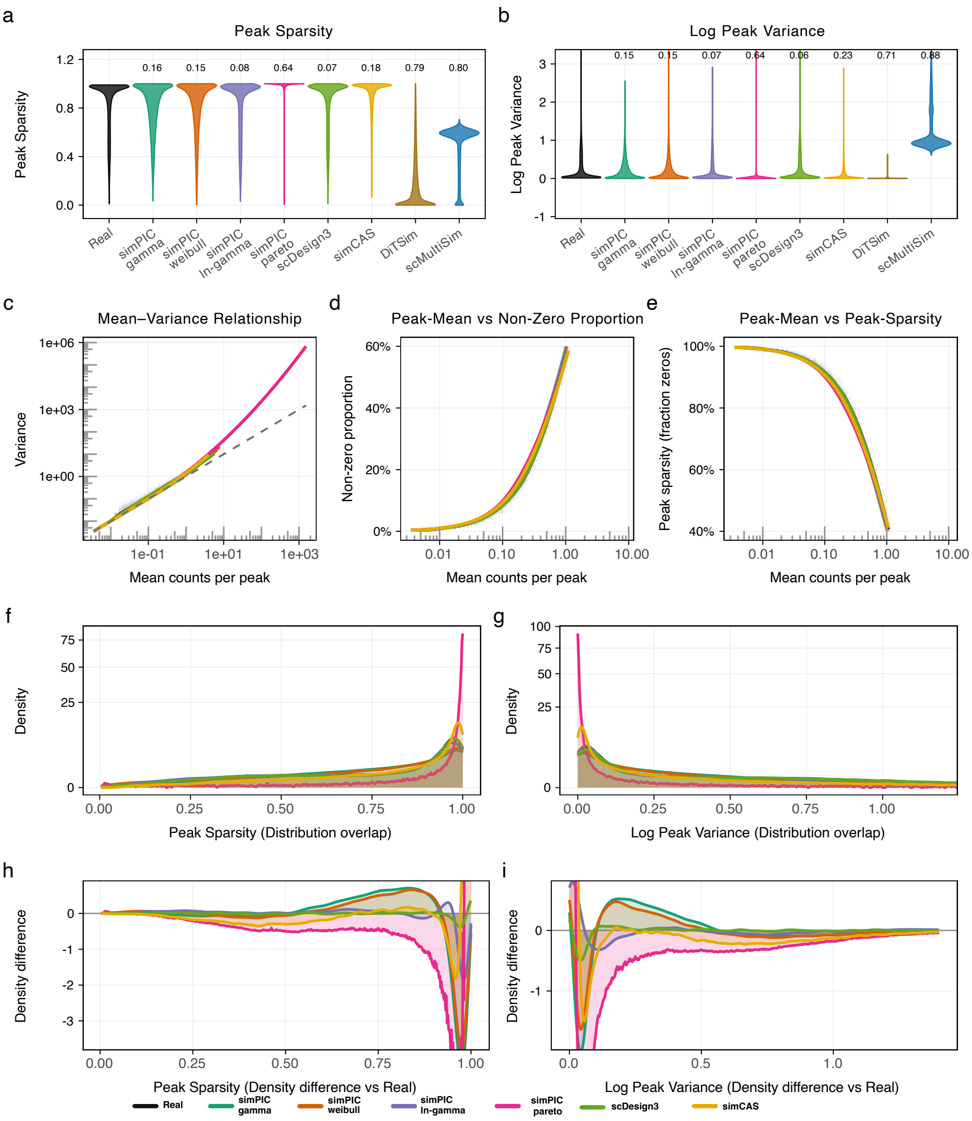

PBMC10k_CD14_Monocytes

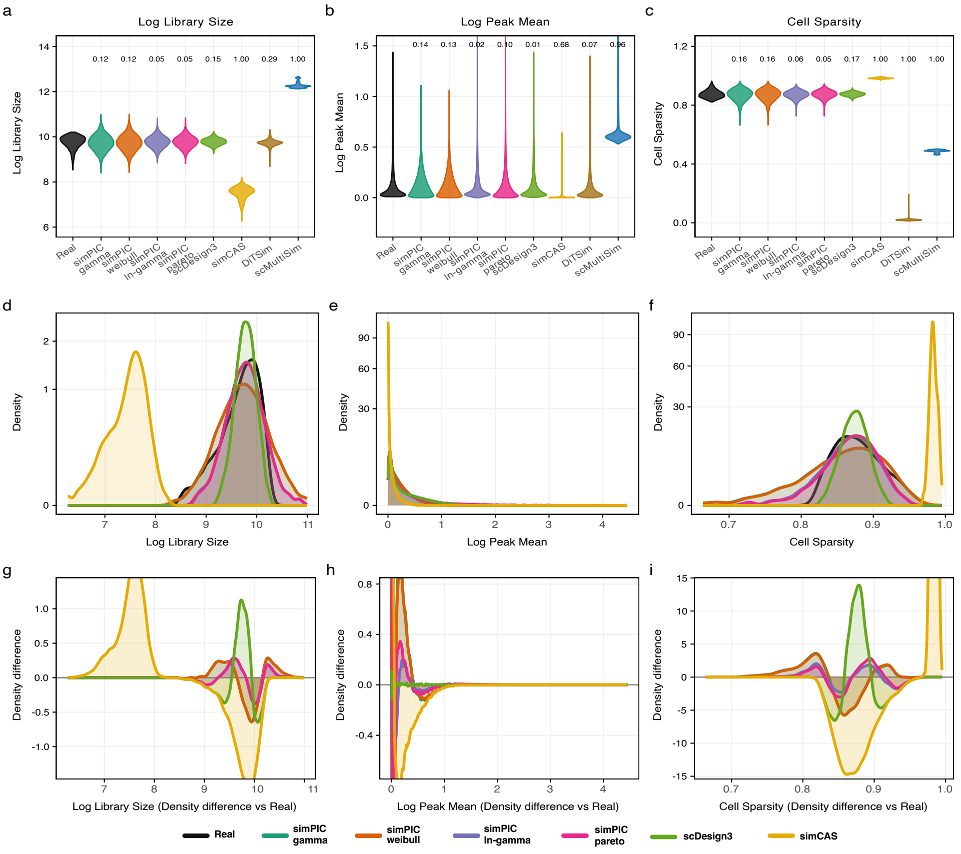

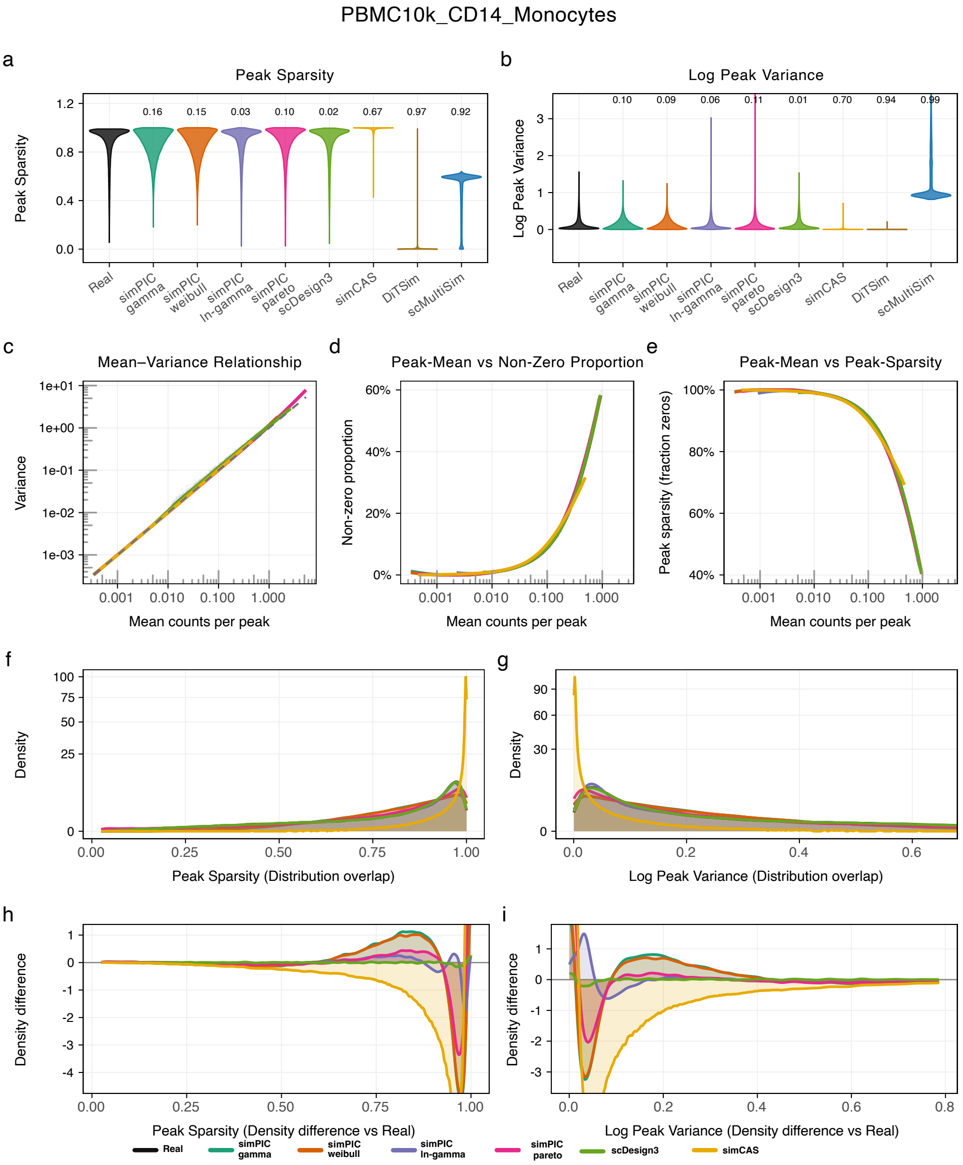

PBMC10k_CD14_Monocytes
a
Peak Sparsity
Peak Sparsity
0.16
0.15
0.03
0.10
0.02
0.67
0.97
0.92
Real
simPIC gamma
simPIC weibull
simPIC ln-gamma
simPIC pareto
scDesign3
simCAS
DiTSim
scMultiSim
b
Log Peak Variance
Log Peak Variance
0.10
0.09
0.06
0.11
0.01
0.70
0.94
0.99
c
Mean−Variance Relationship
Variance
Mean counts per peak
d
Peak-Mean vs Non-Zero Proportion
Non-zero proportion
Mean counts per peak
e
Peak-Mean vs Peak-Sparsity
Peak sparsity (fraction zeros)
Mean counts per peak
f
Density
Peak Sparsity (Distribution overlap)
g
Density
Log Peak Variance (Distribution overlap)
h
Density difference
Peak Sparsity (Density difference vs Real)
i
Density difference
Log Peak Variance (Density difference vs Real)
Real
simPIC gamma
simPIC weibull
simPIC ln-gamma
simPIC pareto
scDesign3
simCAS

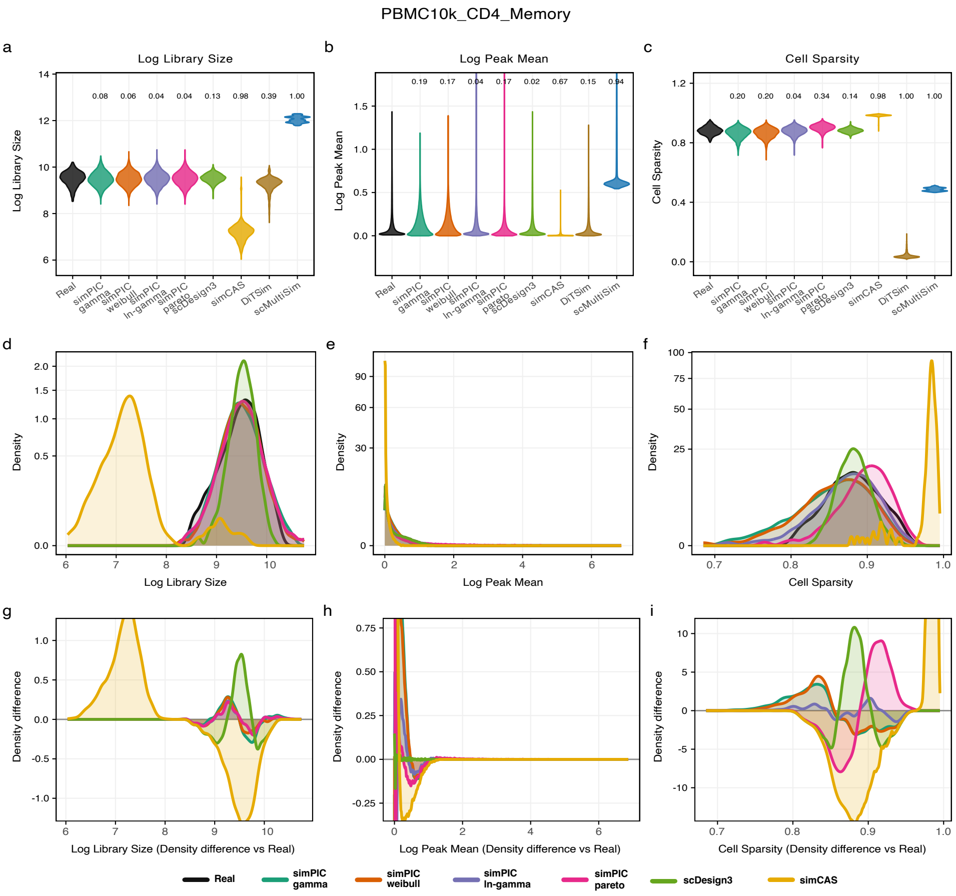

PBMC10k_CD4_Memory
a
Log Library Size
b
Log Peak Mean
c
Cell Sparsity
0.08 0.06 0.04 0.04 0.13 0.98 0.39 1.00
0.19 0.17 0.04 0.17 0.02 0.67 0.15 0.94
0.20 0.20 0.04 0.34 0.14 0.98 1.00 1.00
Real
simPIC gamma
simPIC weibull
simPIC ln-gamma
simPIC pareto
scDesign3
simCAS
DiTSim
scMultiSim
d
e
f
Density
Log Library Size
Log Peak Mean
Cell Sparsity
g
h
i
Density difference
Log Library Size (Density difference vs Real)
Log Peak Mean (Density difference vs Real)
Cell Sparsity (Density difference vs Real)

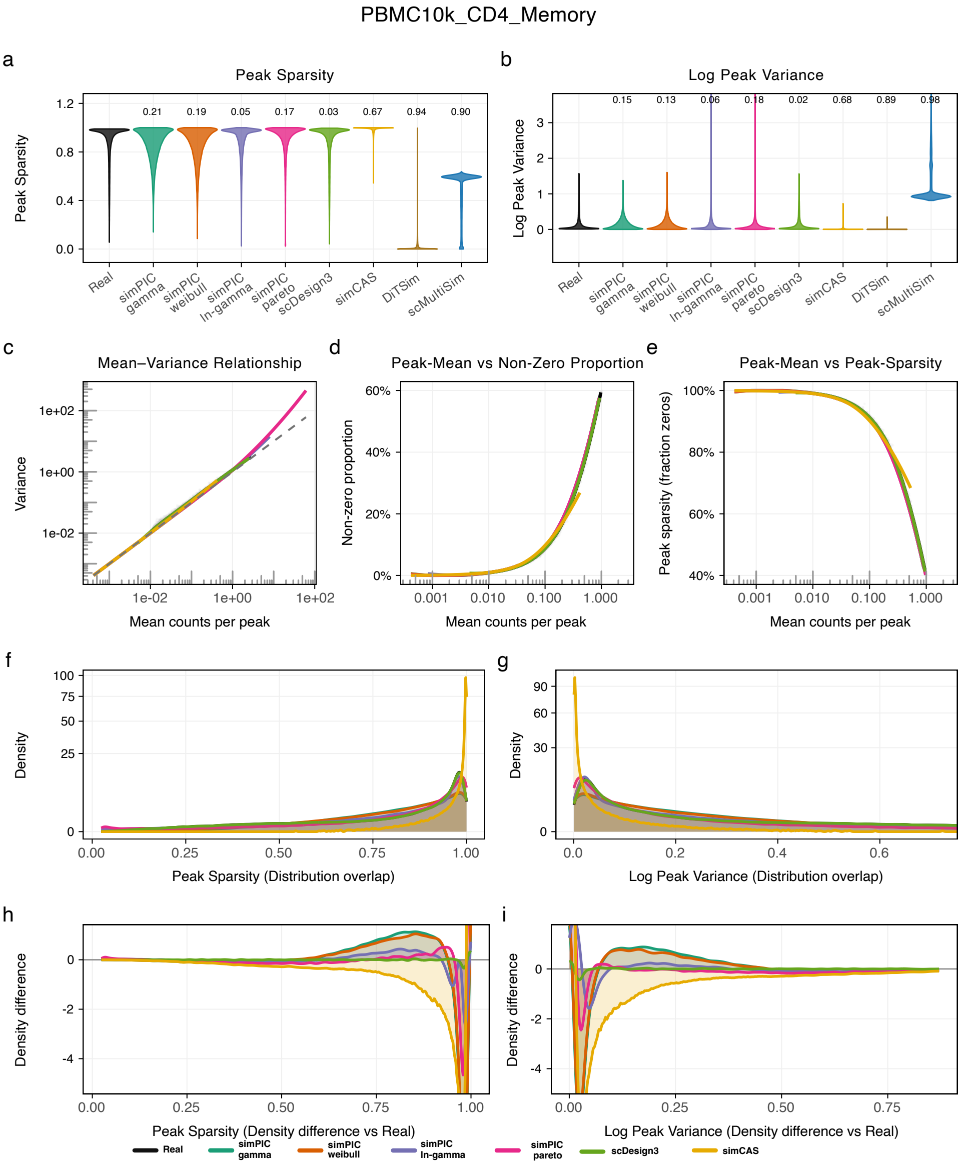
PBMC10k_CD4_Memory
a
Peak Sparsity
Peak Sparsity
0.21
0.19
0.05
0.17
0.03
0.67
0.94
0.90
Real
simPIC gamma
simPIC weibull
simPIC ln-gamma
simPIC pareto
scDesign3
simCAS
DiTSim
scMultiSim
b
Log Peak Variance
Log Peak Variance
0.15
0.13
0.06
0.18
0.02
0.68
0.89
0.98
c
Mean–Variance Relationship
Variance
Mean counts per peak
d
Peak-Mean vs Non-Zero Proportion
Non-zero proportion
Mean counts per peak
e
Peak-Mean vs Peak-Sparsity
Peak sparsity (fraction zeros)
Mean counts per peak
f
Density
Peak Sparsity (Distribution overlap)
g
Density
Log Peak Variance (Distribution overlap)
h
Density difference
Peak Sparsity (Density difference vs Real)
i
Density difference
Log Peak Variance (Density difference vs Real)
Real
simPIC gamma
simPIC weibull
simPIC ln-gamma
simPIC pareto
scDesign3
simCAS

PBMC10k_CD4_Naive

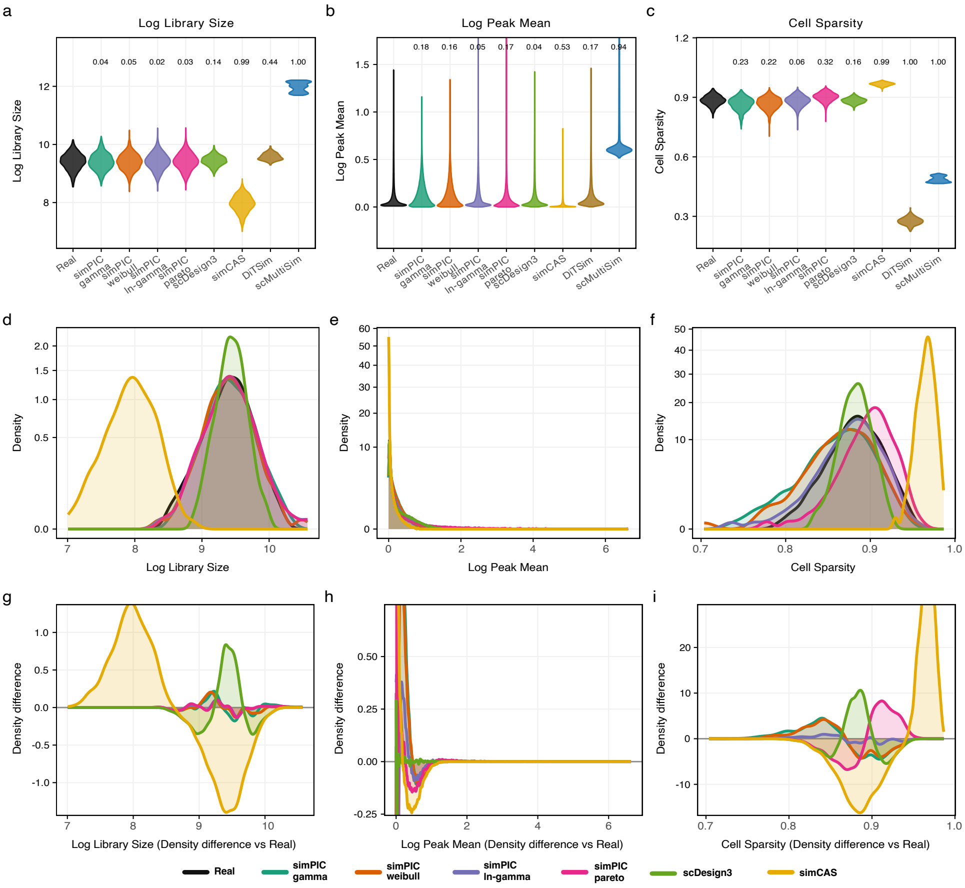

PBMC10k_CD4_Naive

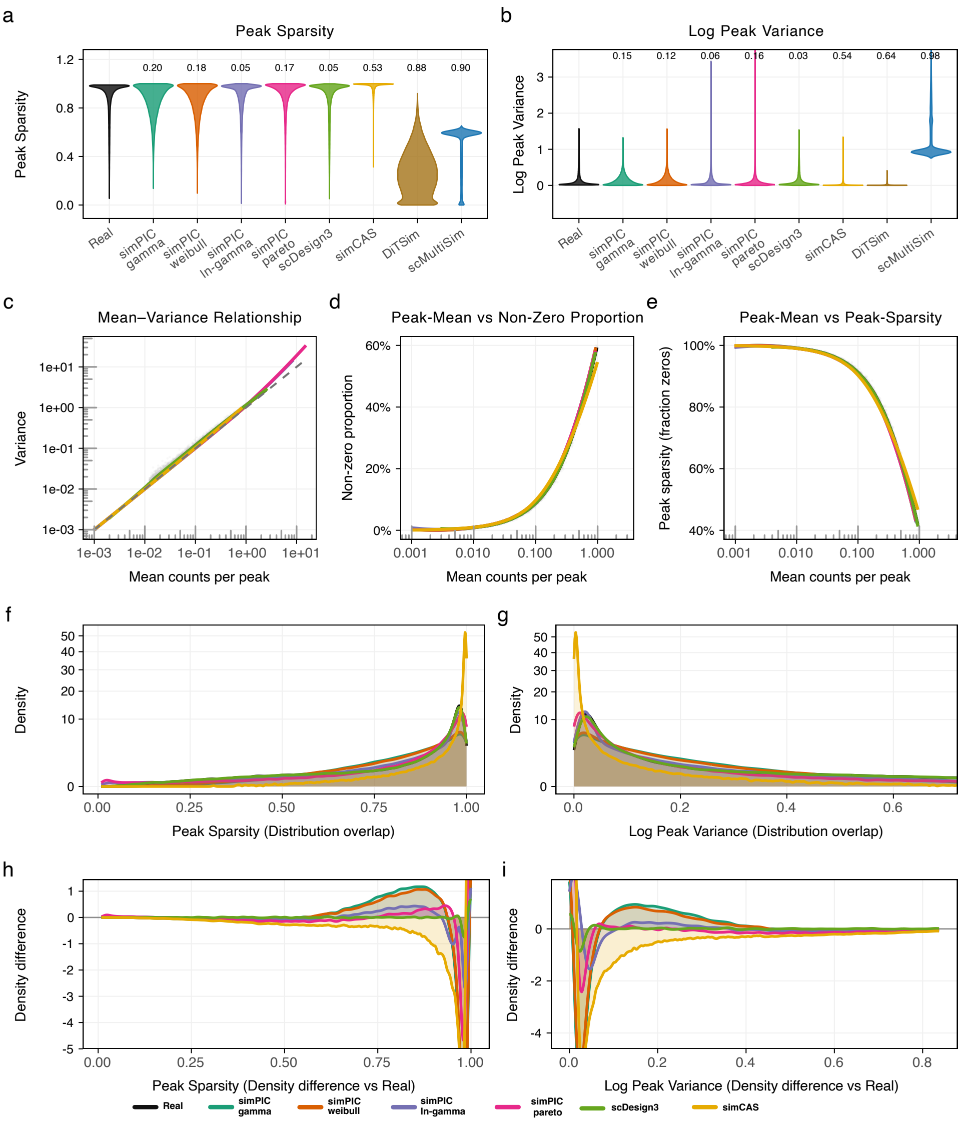

PBMC10k_CD8_naive

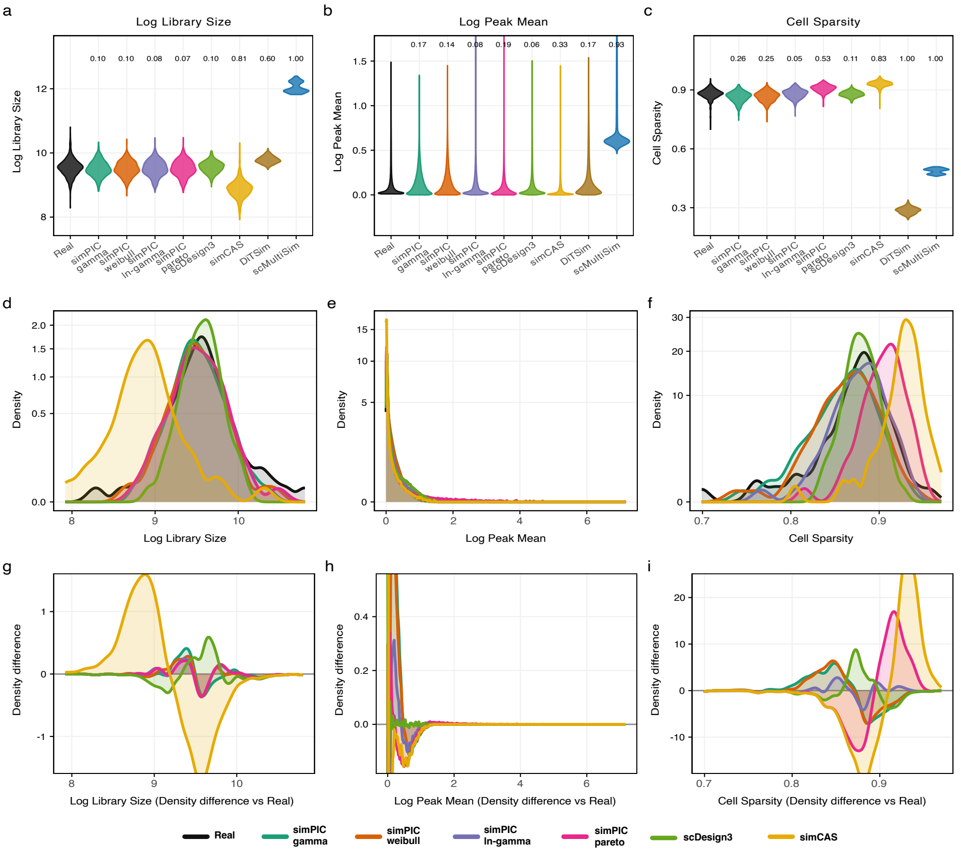

PBMC10k_CD8_naive

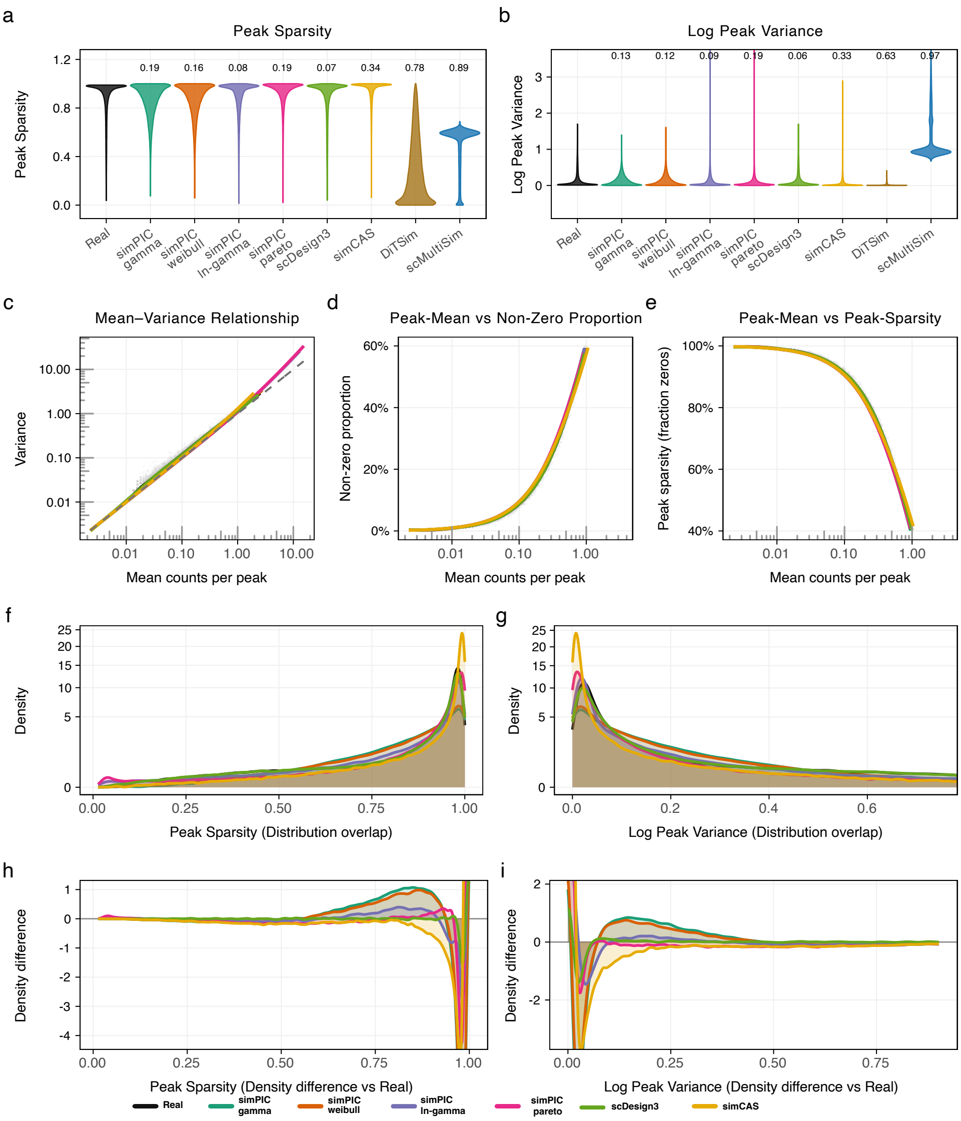

PBMC10k_CD8_effector

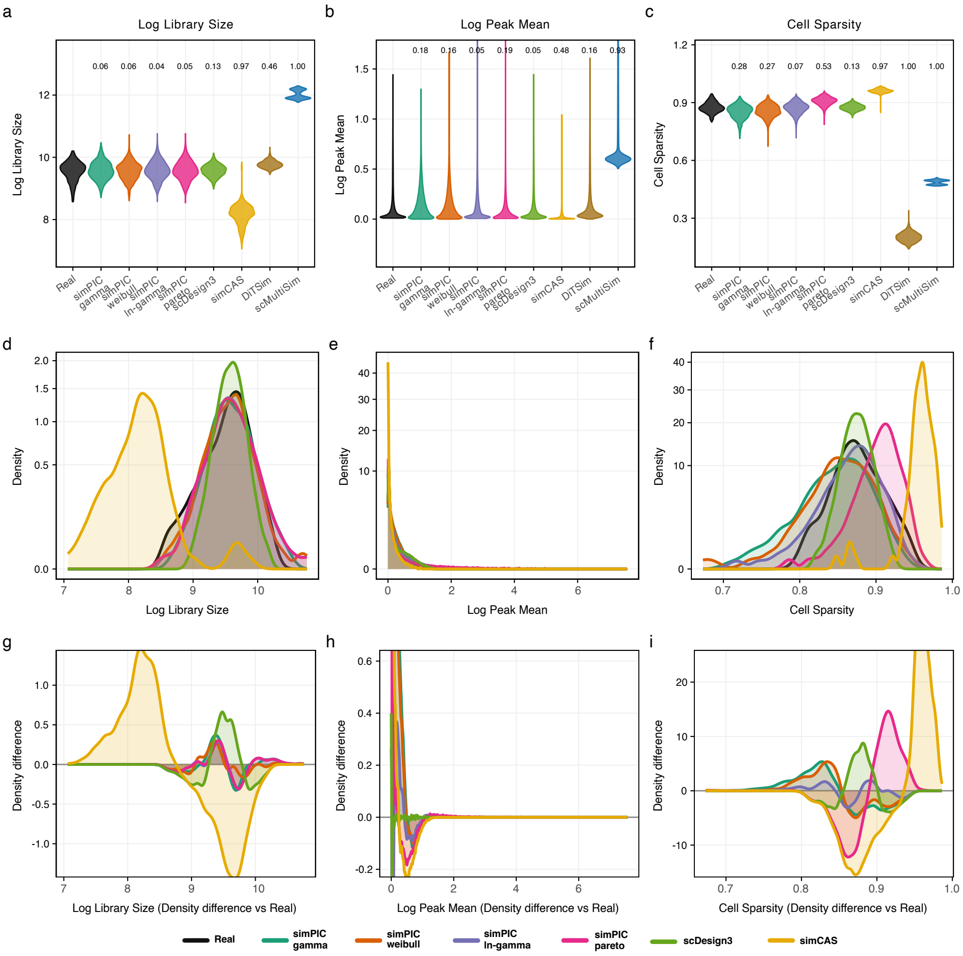

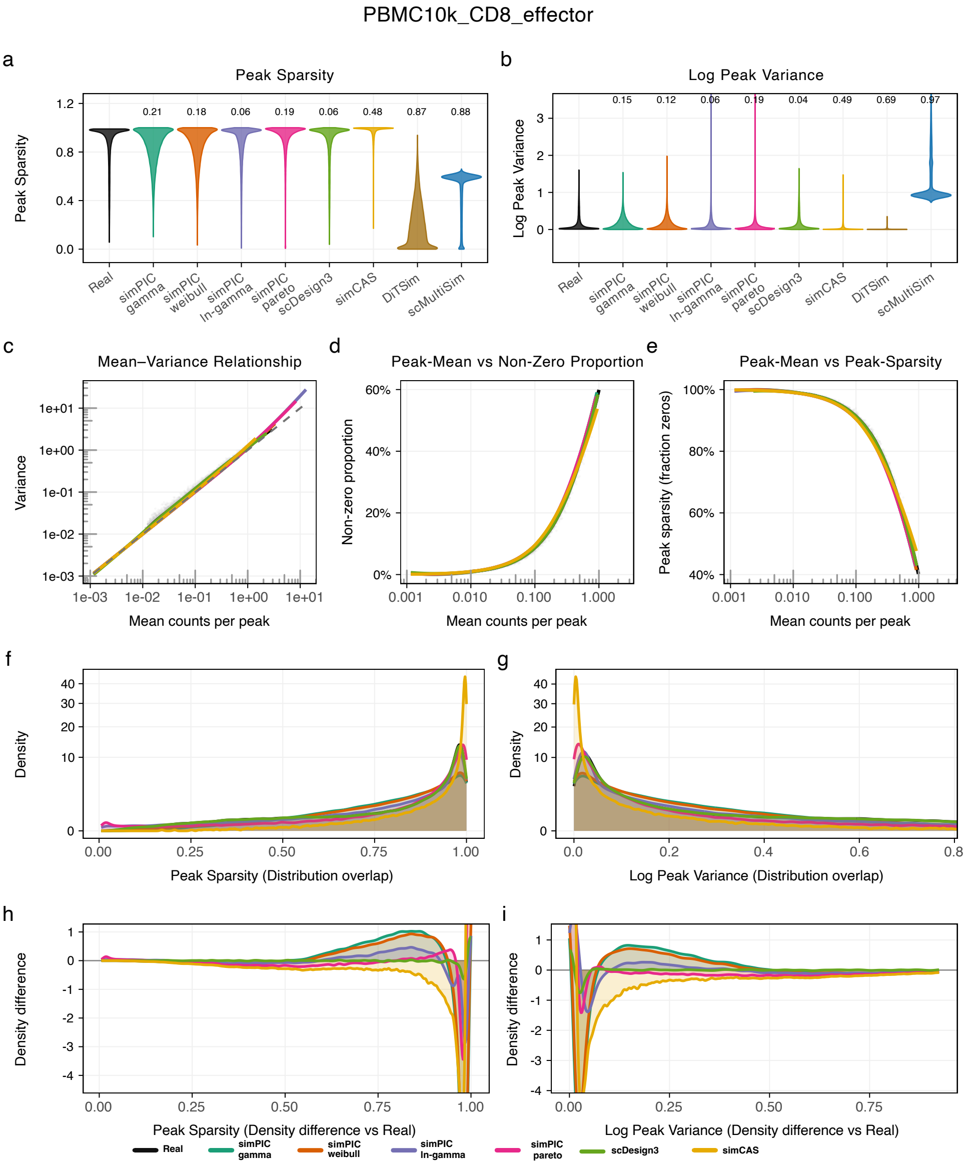
PBMC10k_CD8_effector
a
Peak Sparsity
Peak Sparsity
0.21
0.18
0.06
0.19
0.06
0.48
0.87
0.88
Real
simPIC gamma
simPIC weibull
simPIC ln-gamma
simPIC pareto
scDesign3
simCAS
DiTSim
scMultiSim
b
Log Peak Variance
Log Peak Variance
0.15
0.12
0.06
0.19
0.04
0.49
0.69
0.97
c
Mean–Variance Relationship
Variance
Mean counts per peak
d
Peak-Mean vs Non-Zero Proportion
Non-zero proportion
Mean counts per peak
e
Peak-Mean vs Peak-Sparsity
Peak sparsity (fraction zeros)
Mean counts per peak
f
Density
Peak Sparsity (Distribution overlap)
g
Density
Log Peak Variance (Distribution overlap)
h
Density difference
Peak Sparsity (Density difference vs Real)
i
Density difference
Log Peak Variance (Density difference vs Real)
Real
simPIC gamma
simPIC weibull
simPIC ln-gamma
simPIC pareto
scDesign3
simCAS

PBMC10k_B_cell_progenitor

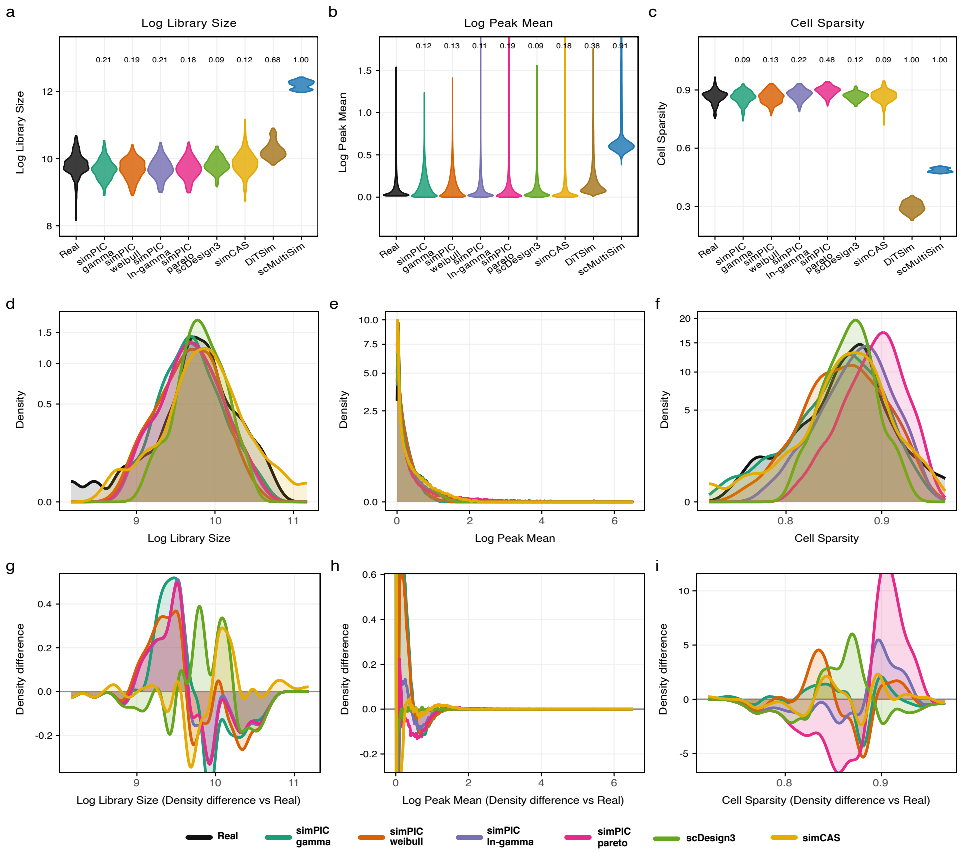

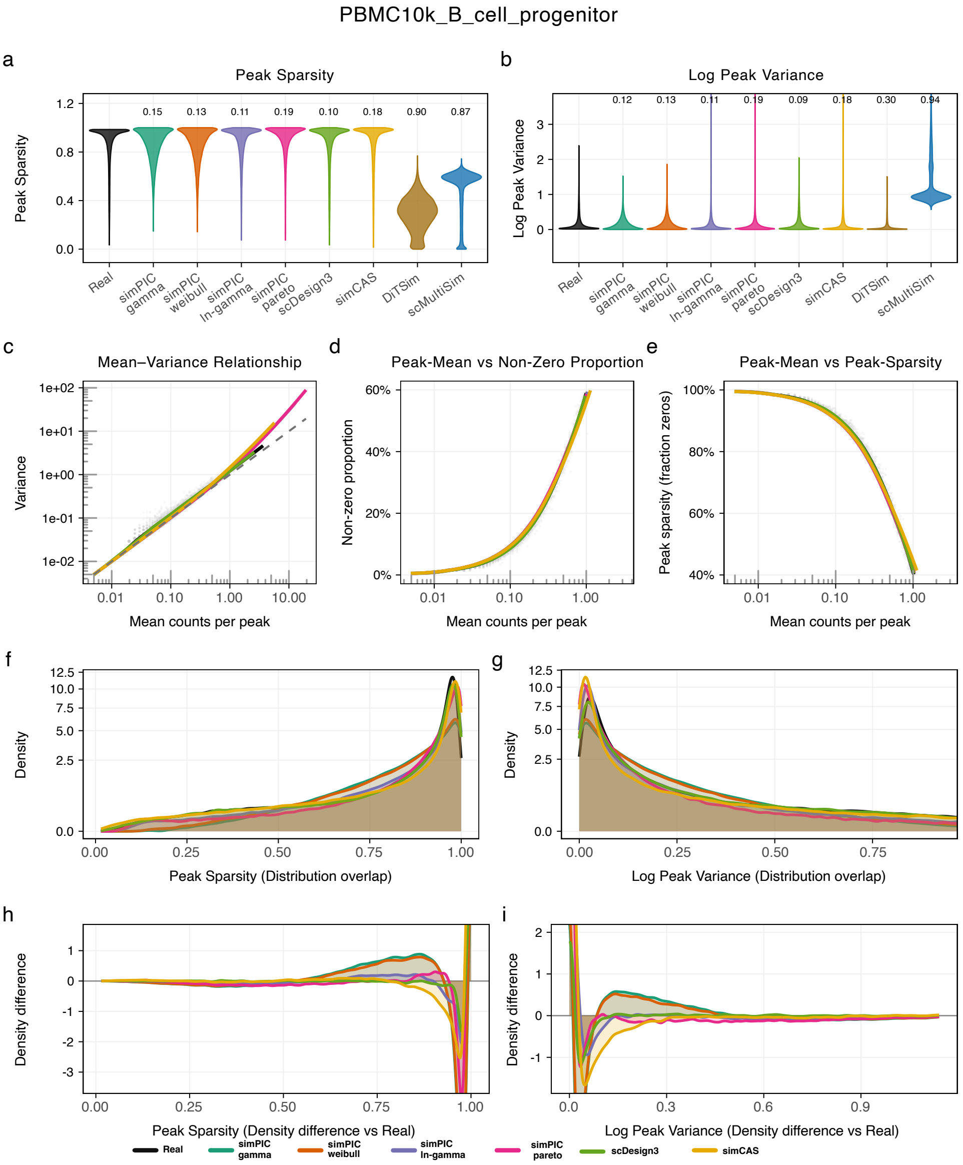

PBMC10k_B_cell_progenitor
a
Peak Sparsity
Peak Sparsity
0.15 0.13 0.11 0.19 0.10 0.18 0.90 0.87
Real
simPIC gamma
simPIC weibull
simPIC ln-gamma
simPIC pareto
scDesign3
simCAS
DiTSim
scMultiSim
b
Log Peak Variance
Log Peak Variance
0.12 0.13 0.11 0.19 0.09 0.18 0.30 0.94
c
Mean–Variance Relationship
Variance
Mean counts per peak
d
Peak-Mean vs Non-Zero Proportion
Non-zero proportion
Mean counts per peak
e
Peak-Mean vs Peak-Sparsity
Peak sparsity (fraction zeros)
Mean counts per peak
f
Density
Peak Sparsity (Distribution overlap)
g
Density
Log Peak Variance (Distribution overlap)
h
Density difference
Peak Sparsity (Density difference vs Real)
i
Density difference
Log Peak Variance (Density difference vs Real)
Real
simPIC gamma
simPIC weibull
simPIC ln-gamma
simPIC pareto
scDesign3
simCAS

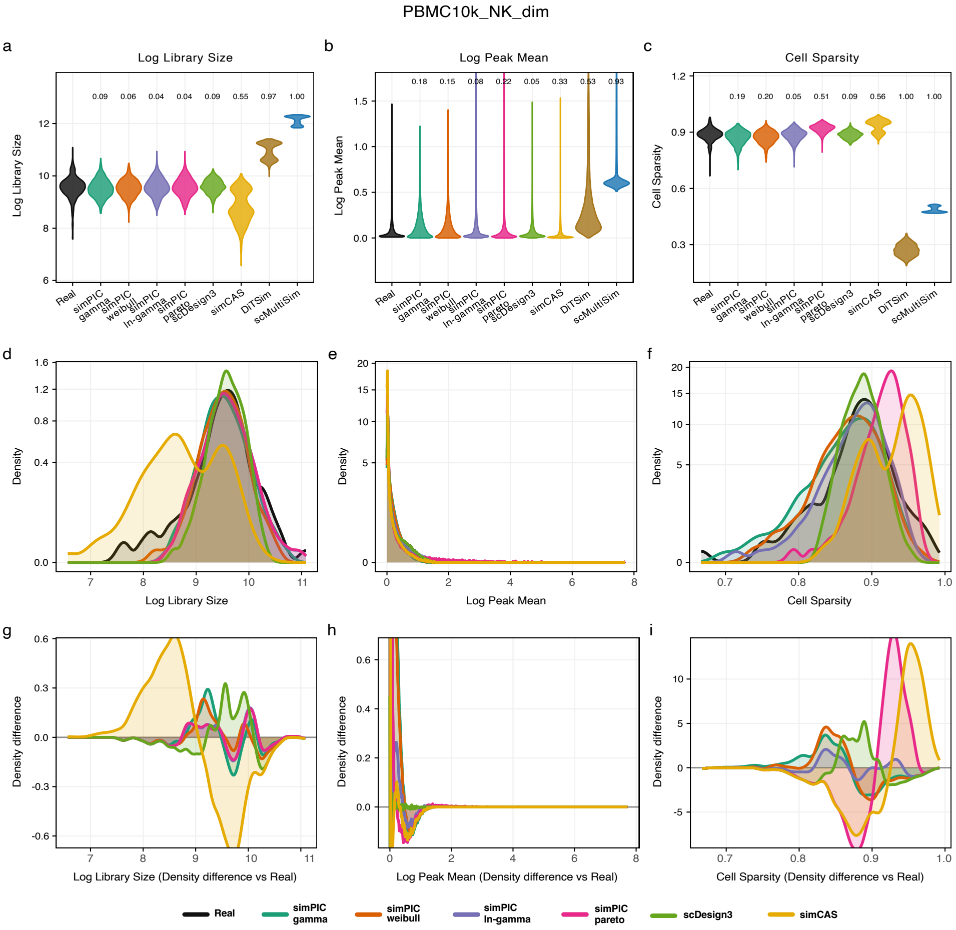
PBMC10k_NK_dim
a
Log Library Size
b
Log Peak Mean
c
Cell Sparsity
d
e
f
g
h
i
Log Library Size
Log Peak Mean
Cell Sparsity
Density
Density difference
Log Library Size (Density difference vs Real)
Log Peak Mean (Density difference vs Real)
Cell Sparsity (Density difference vs Real)
Real
simPIC gamma
simPIC weibull
simPIC ln-gamma
simPIC pareto
scDesign3
simCAS
DiTSim
scMultiSim

PBMC10k_NK_dim

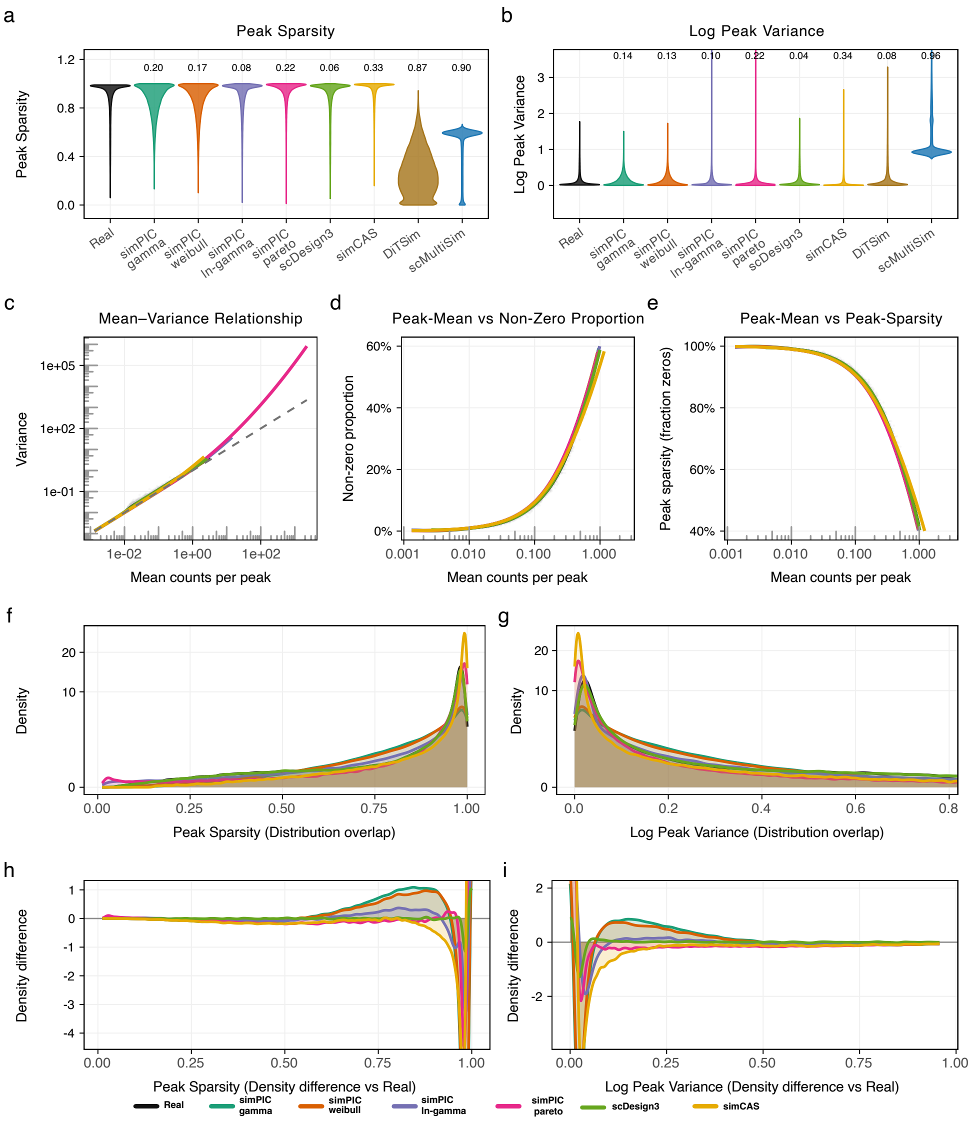

Satpathy_CD8_Naive

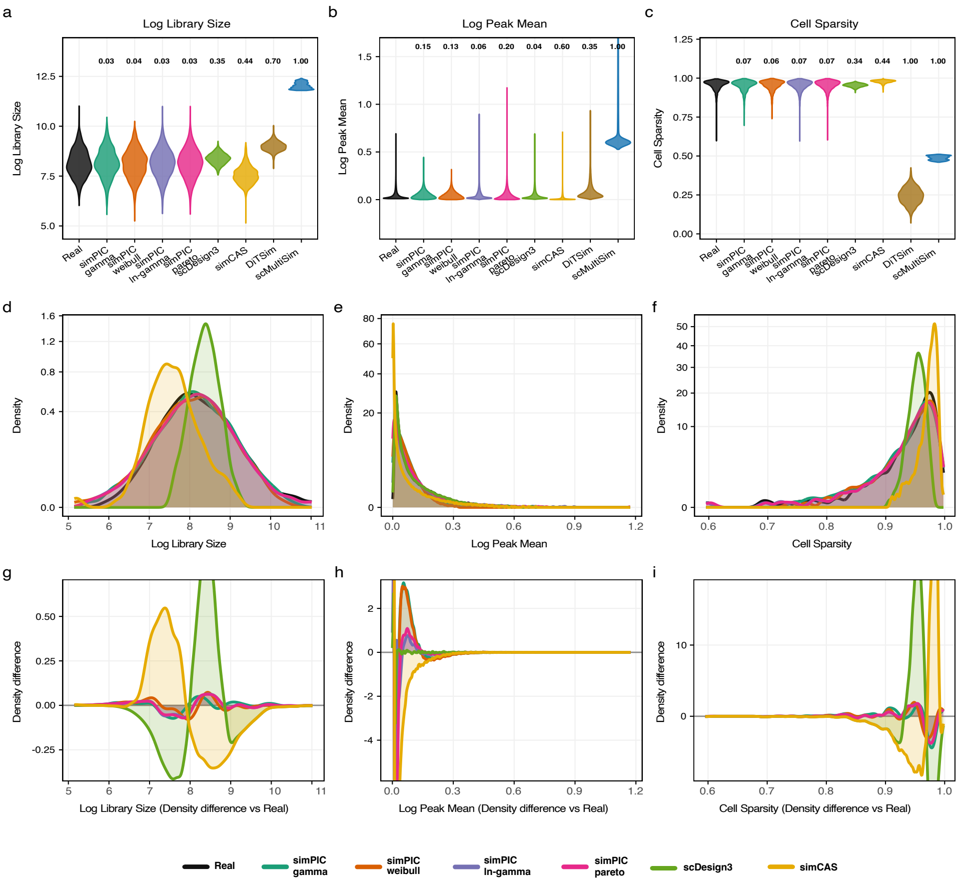

Satpathy_CD8_Naive

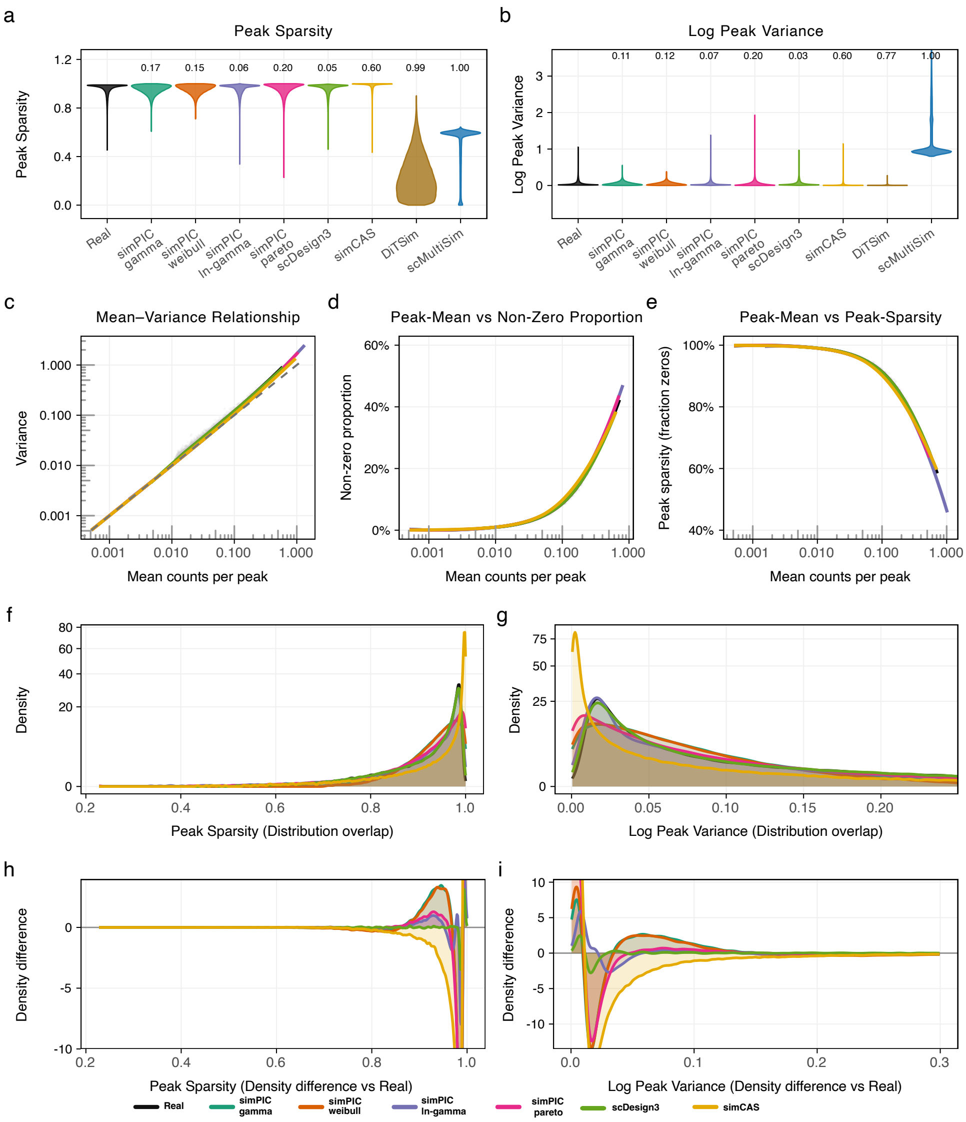

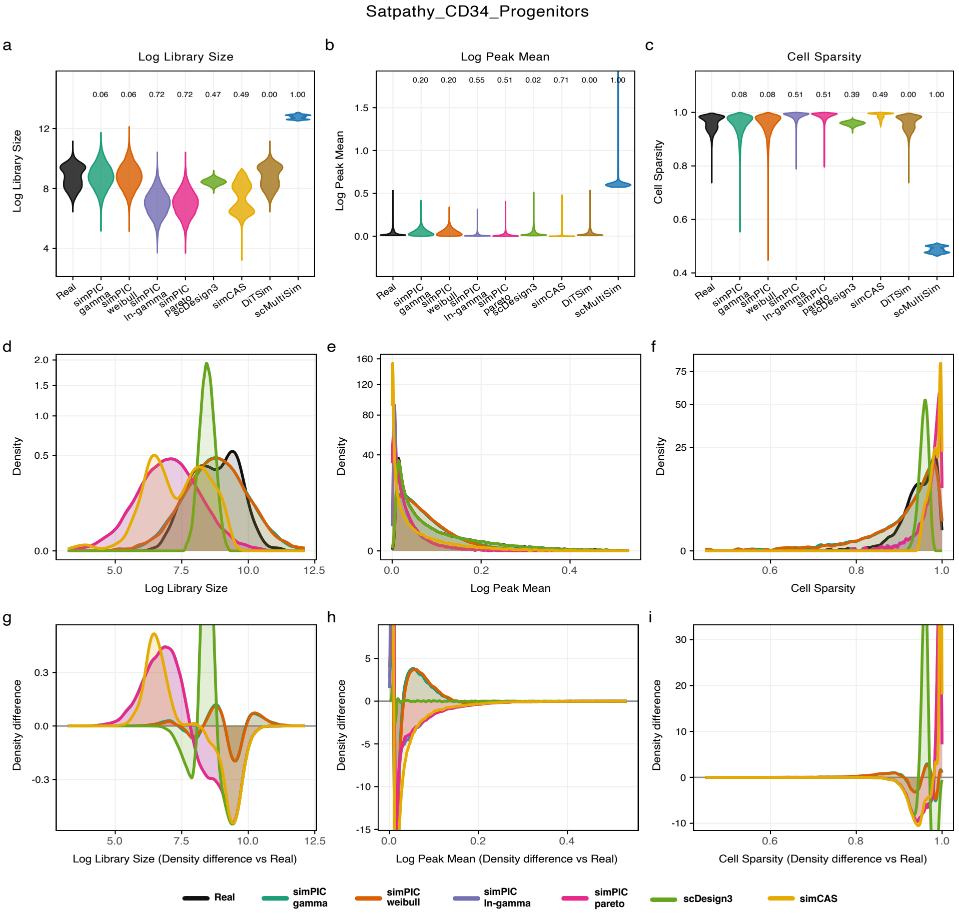

Satpathy_CD34_Progenitors
a
Log Library Size
b
Log Peak Mean
c
Cell Sparsity
d
e
f
g
h
i
Real
simPIC gamma
simPIC weibull
simPIC ln-gamma
simPIC pareto
scDesign3
simCAS
DiTSim
scMultiSim
Log Library Size (Density difference vs Real)
Log Peak Mean (Density difference vs Real)
Cell Sparsity (Density difference vs Real)

### Satpathy_CD34_Progenitors

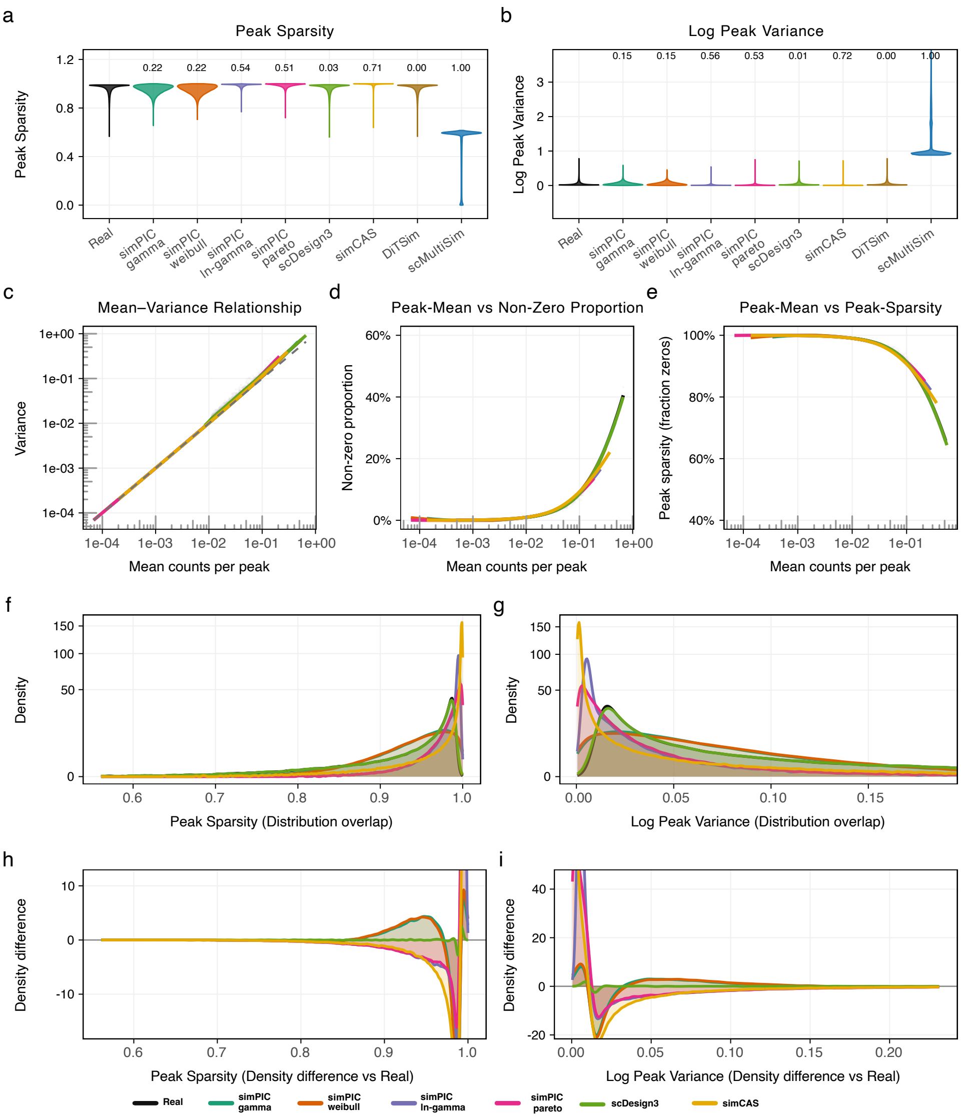

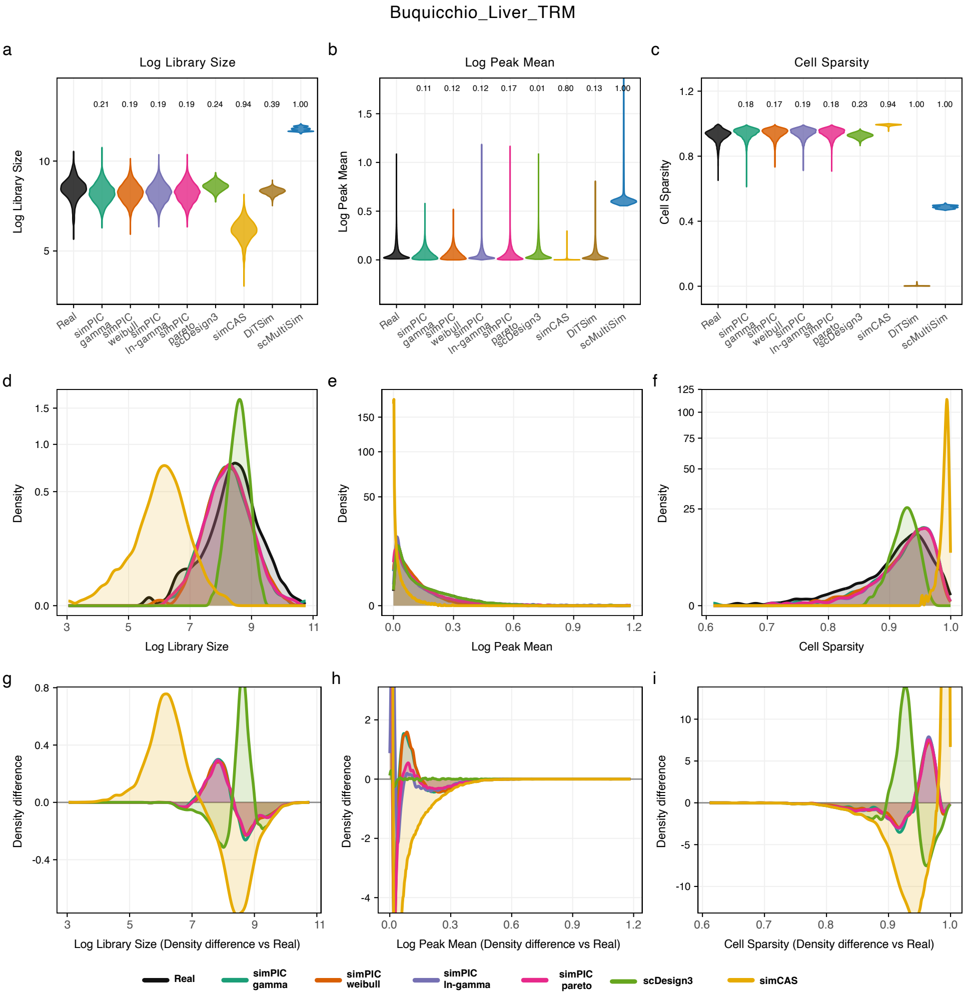
Buquicchio_Liver_TRM
a
Log Library Size
b
Log Peak Mean
c
Cell Sparsity
d
e
f
g
h
i
Real
simPIC gamma
simPIC weibull
simPIC ln-gamma
simPIC pareto
scDesign3
simCAS
DiTSim
scMultiSim
Log Library Size (Density difference vs Real)
Log Peak Mean (Density difference vs Real)
Cell Sparsity (Density difference vs Real)

Buquicchio_Liver_TRM

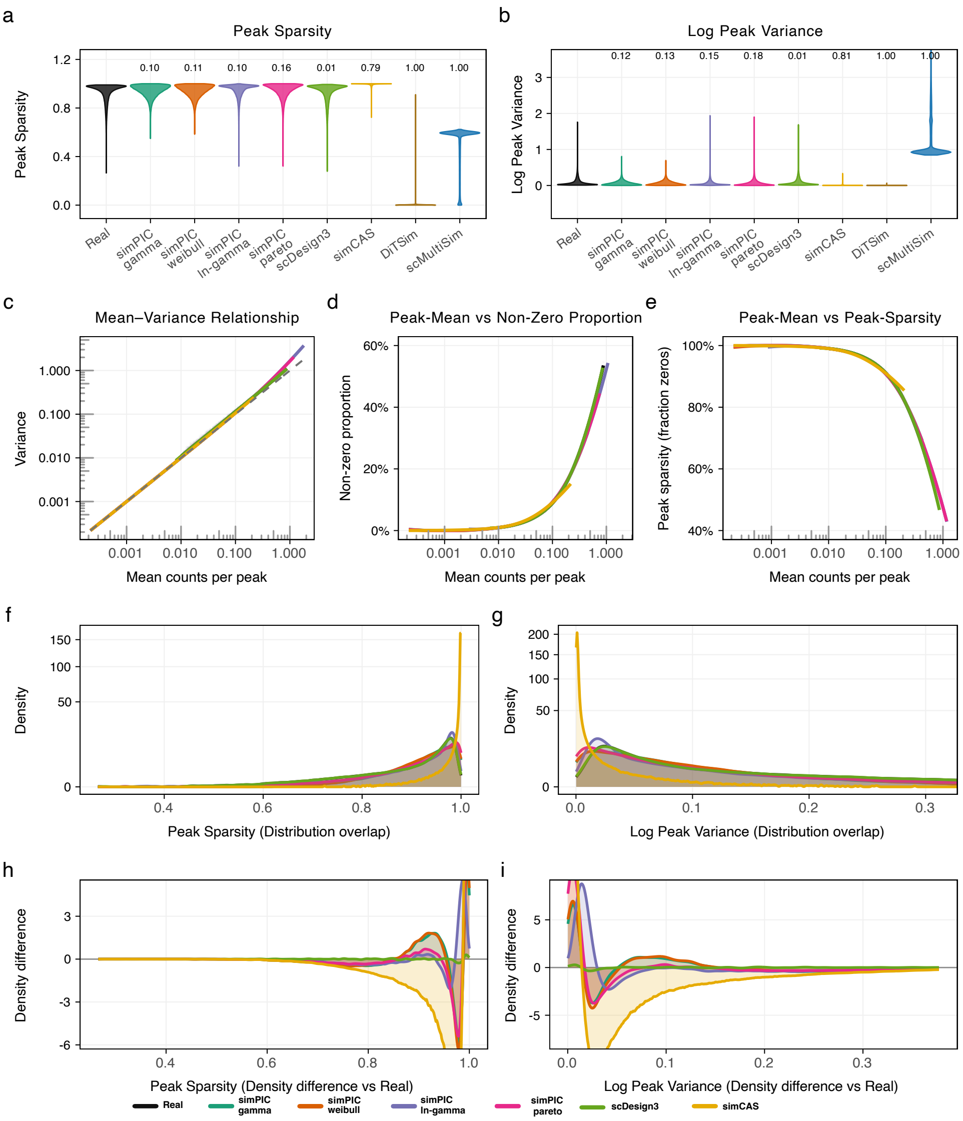

### Evaluation metrics for additional datasets

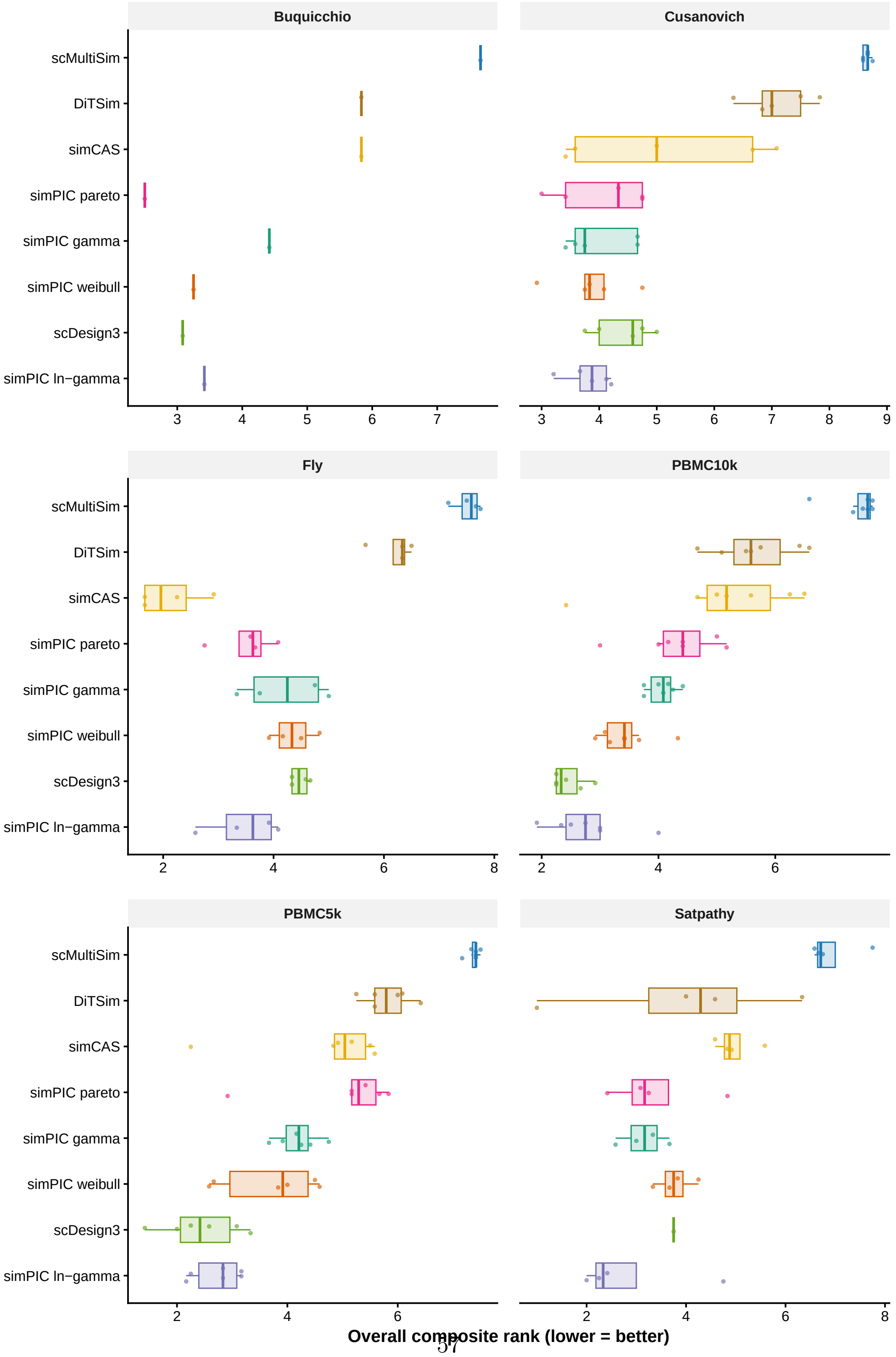




**Fig. S4** Comparative boxplots for evaluation metrics MAD, MAE, RMSE and 1-PCC across library size, peak means and cell sparsity in PBMC5k, Cusanovich and Fly brain dataset for SIMPIC(g) in blue, SIMPIC(w) in green and simCAS in purple. Each point represents a cell-type.

### simPIC simulates multiple cell types and batch effects with configurable group structure

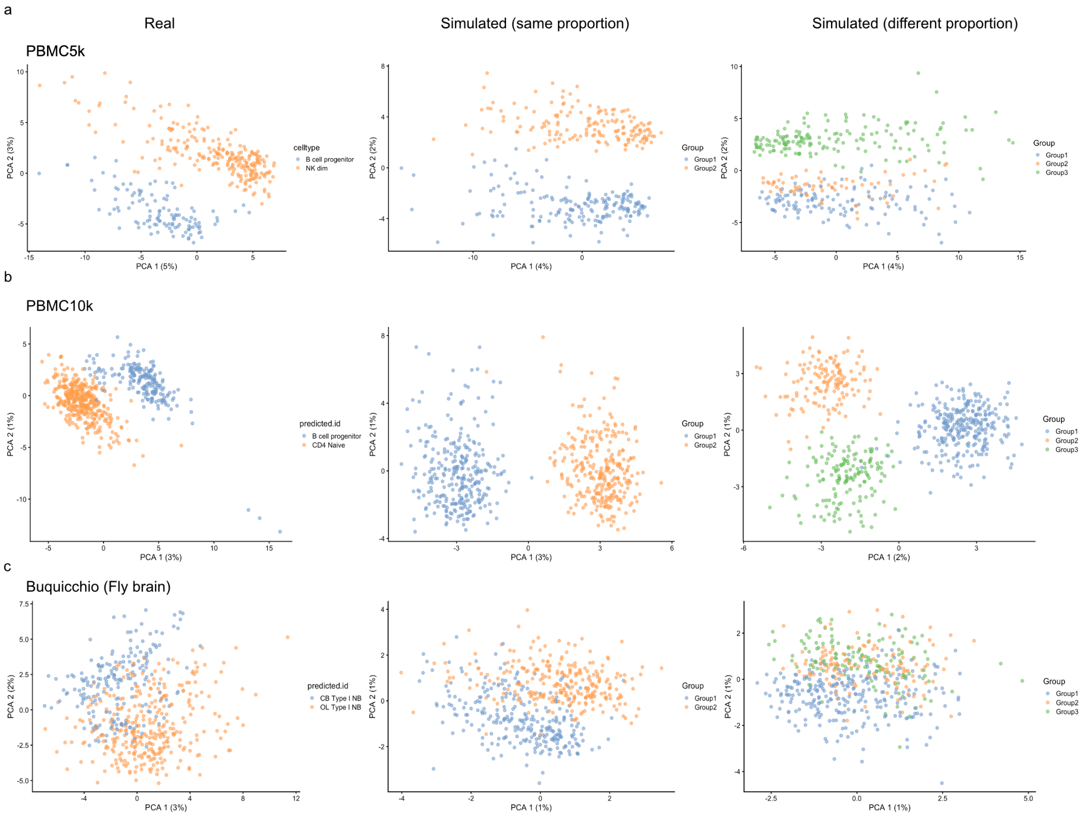


**Fig. S5** Principal Component Analysis (PCA) plot illustrating the separation of two simulated cell type groups generated using simPIC. Group 1 is represented in blue, and Group 2 in orange, demonstrating that simPIC can simulate multiple cell types while maintaining the same or different proportions across groups.

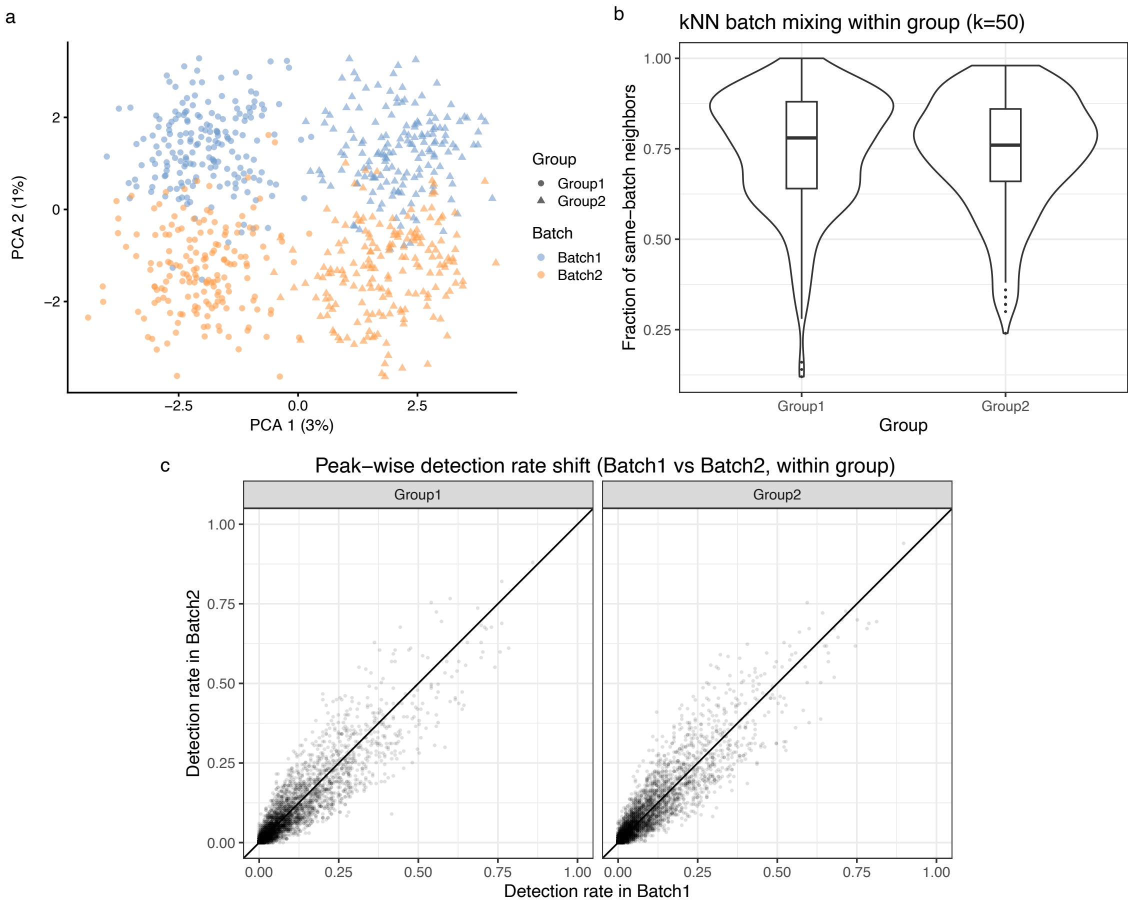


**Fig. S6** a)PCA of simulated cells showing preservation of biological group structure (shapes) while introducing technical batch variation (colours). Batch effects are generated using peak-wise, batch-specific multiplicative factors applied to all cells within each batch.b) kNN batch-mixing (k = 50) quantifies within-group batch-associated structure as the fraction of same-batch neighbours (higher values indicate stronger batch separation).c) Peak-wise detection-rate shift shows per-peak detection rates in Batch1 versus Batch2 within each group, with the diagonal indicating no batch effect; systematic deviations reflect feature-level batch shifts consistent with the multiplicative batch model

#### Simulating population scale data with genetic effects

Clustering diagnostics were done for multiple libraries across the Alzheimer's disease (AD) and non Alzheimer's category. The following figures show all the different examples with varying no of cells for each library

**The below figures are for earlyAD samples**

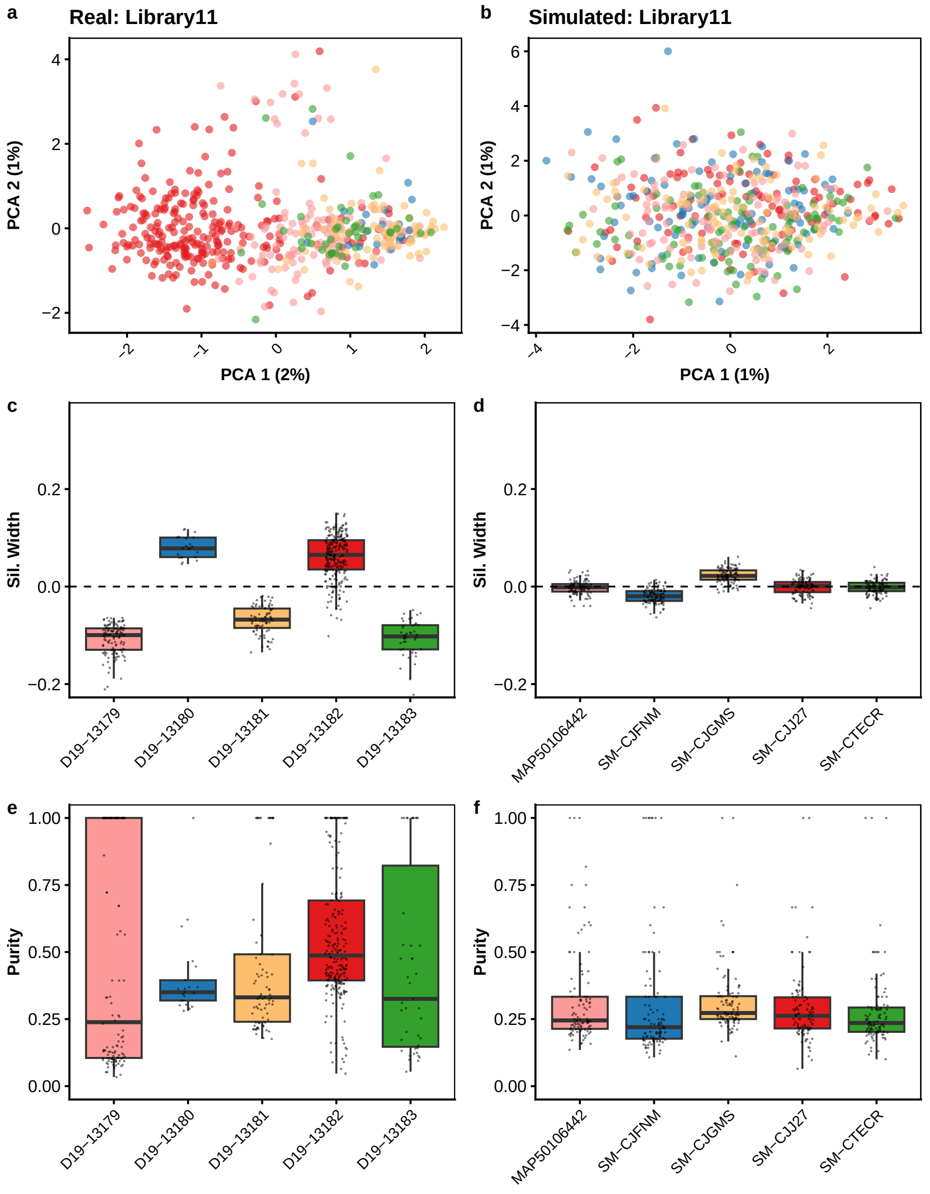


**Fig. S7** Quantifying clustering in real and simPIC-simulated data for early AD library 11 with 1693 peaks and 507 cells. (Top row) PCA plot of real and simulated data, coloured by individual, showing the preservation of individual-specific structure. (Middle row) Silhouette width comparison between real and simulated data, assessing cluster compactness. (Bottom row) Neighbourhood purity analysis, evaluating local structure consistency across real and simulated datasets.

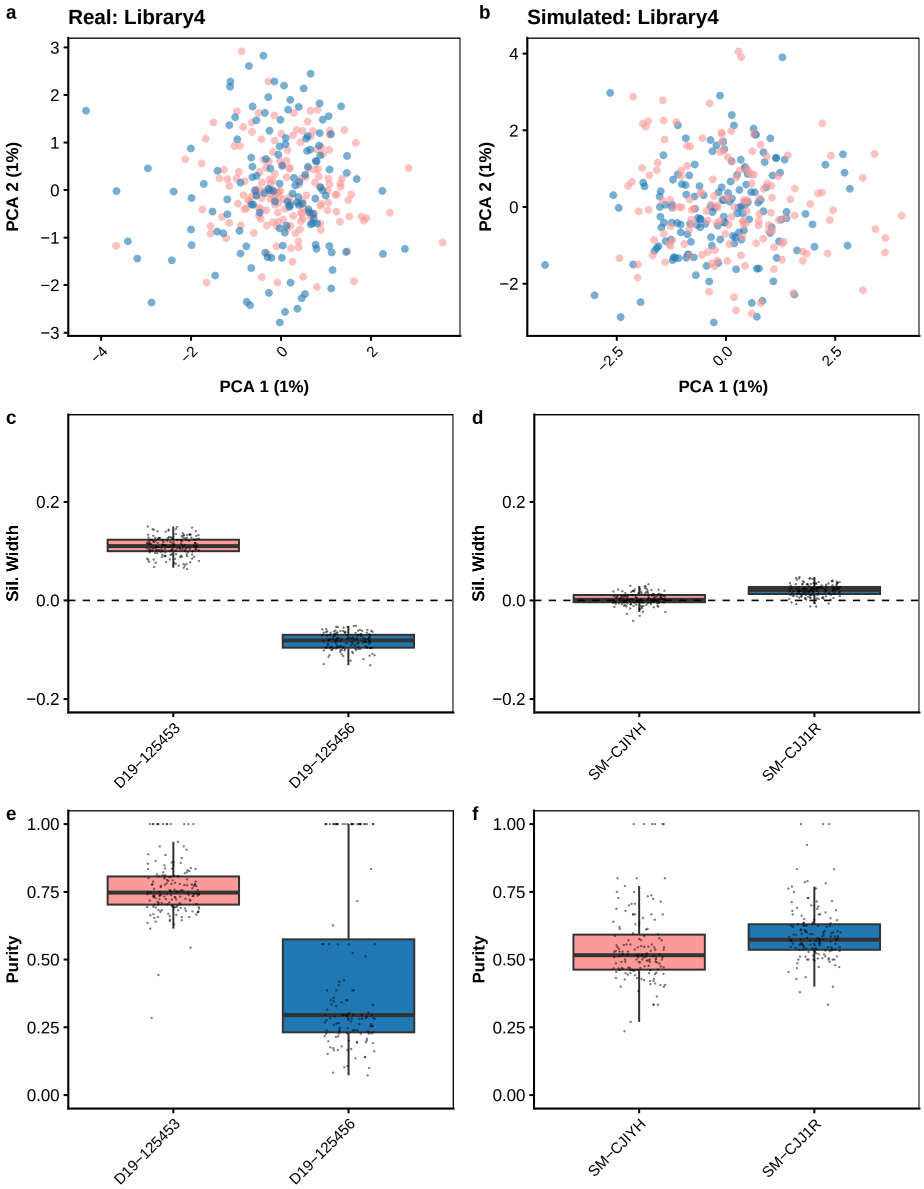


**Fig. S8** Quantifying clustering in real and simPIC-simulated data for early AD library 4 with 1693 peaks and 307 cells. (Top row) PCA plot of real and simulated data, coloured by individual, showing the preservation of individual-specific structure. (Middle row) Silhouette width comparison between real and simulated data, assessing cluster compactness. (Bottom row) Neighbourhood purity analysis, evaluating local structure consistency across real and simulated datasets.

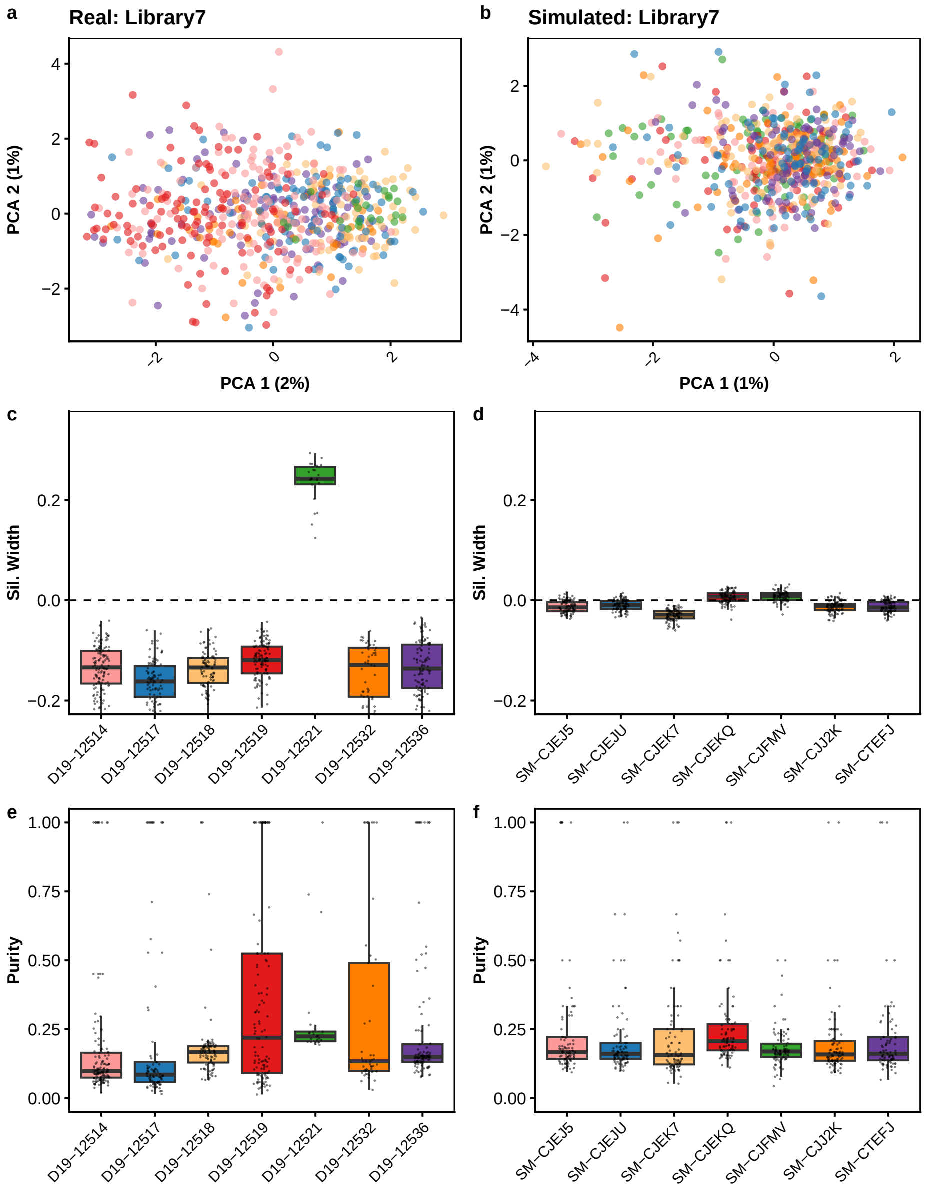


**Fig. S9** Quantifying clustering in real and simPIC-simulated data for early AD library 7 with 1693 peaks and 604 cells. (Top row) PCA plot of real and simulated data, coloured by individual, showing the preservation of individual-specific structure. (Middle row) Silhouette width comparison between real and simulated data, assessing cluster compactness. (Bottom row) Neighbourhood purity analysis, evaluating local structure consistency across real and simulated datasets.

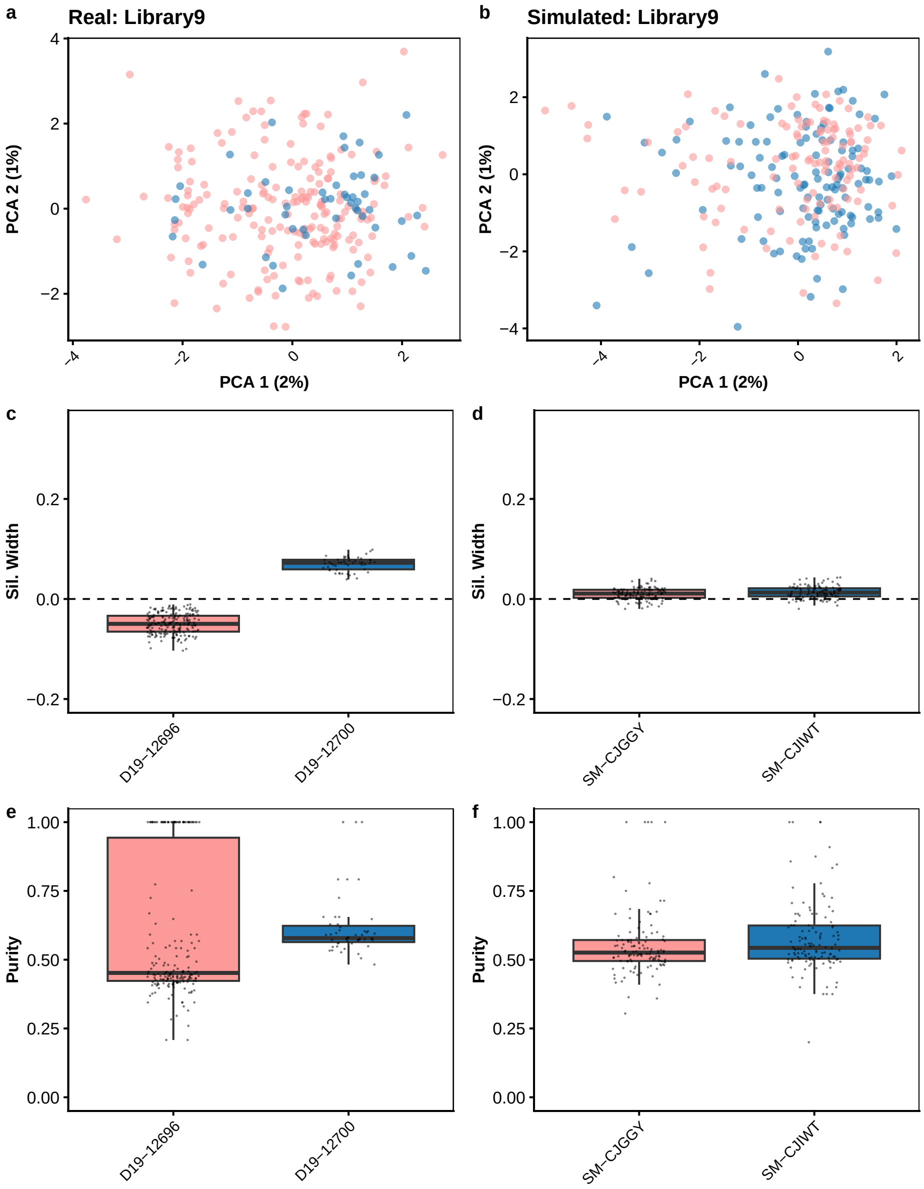


**Fig. S10** Quantifying clustering in real and simPIC-simulated data for early AD library 9 with 1693 peaks and 244 cells. (Top row) PCA plot of real and simulated data, coloured by individual, showing the preservation of individual-specific structure. (Middle row) Silhouette width comparison between real and simulated data, assessing cluster compactness. (Bottom row) Neighbourhood purity analysis, evaluating local structure consistency across real and simulated datasets.

The below figures are for nonAD samples

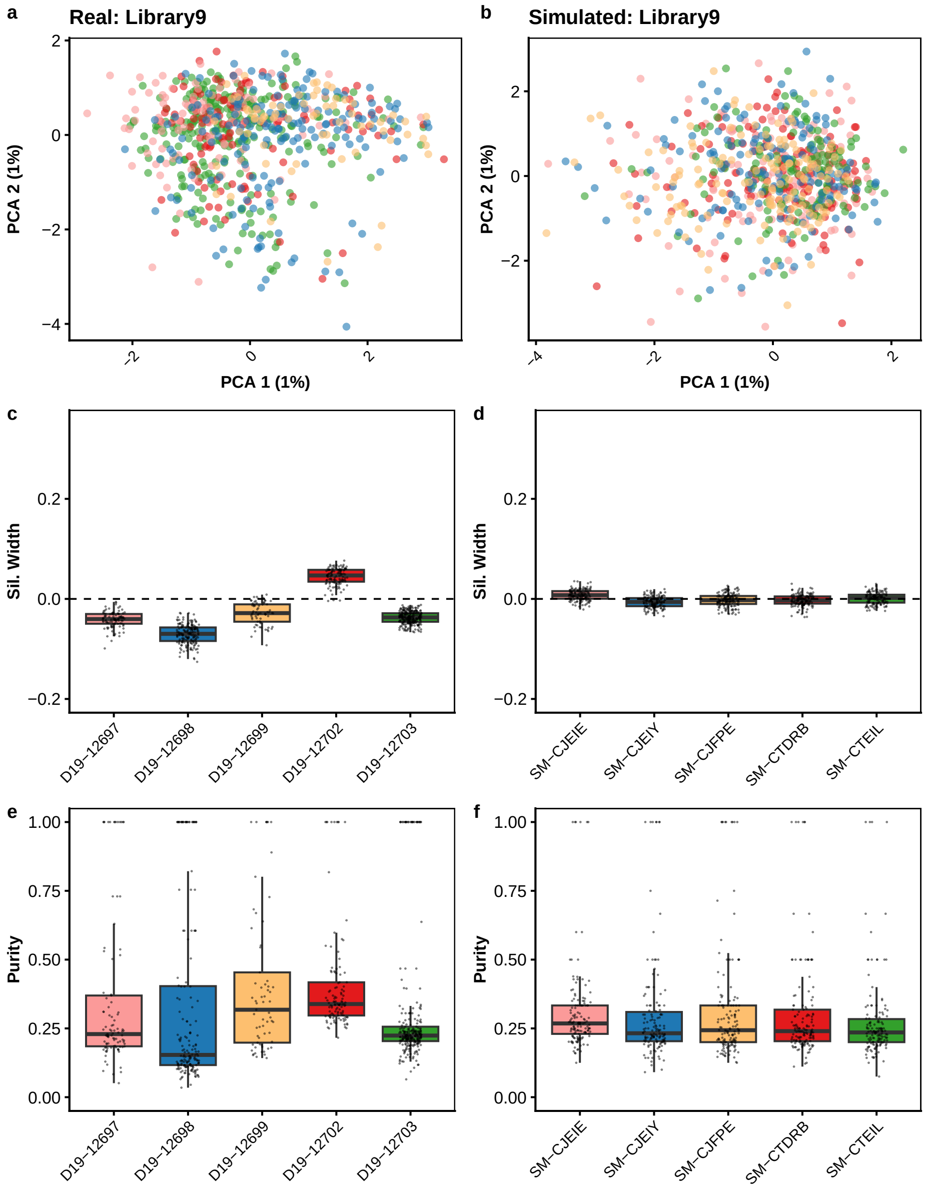


**Fig. S11** Quantifying clustering in real and simPIC-simulated data for non AD library 9 with 1693 peaks and 710 cells. (Top row) PCA plot of real and simulated data, coloured by individual, showing the preservation of individual-specific structure. (Middle row) Silhouette width comparison between real and simulated data, assessing cluster compactness. (Bottom row) Neighbourhood purity analysis, evaluating local structure consistency across real and simulated datasets.

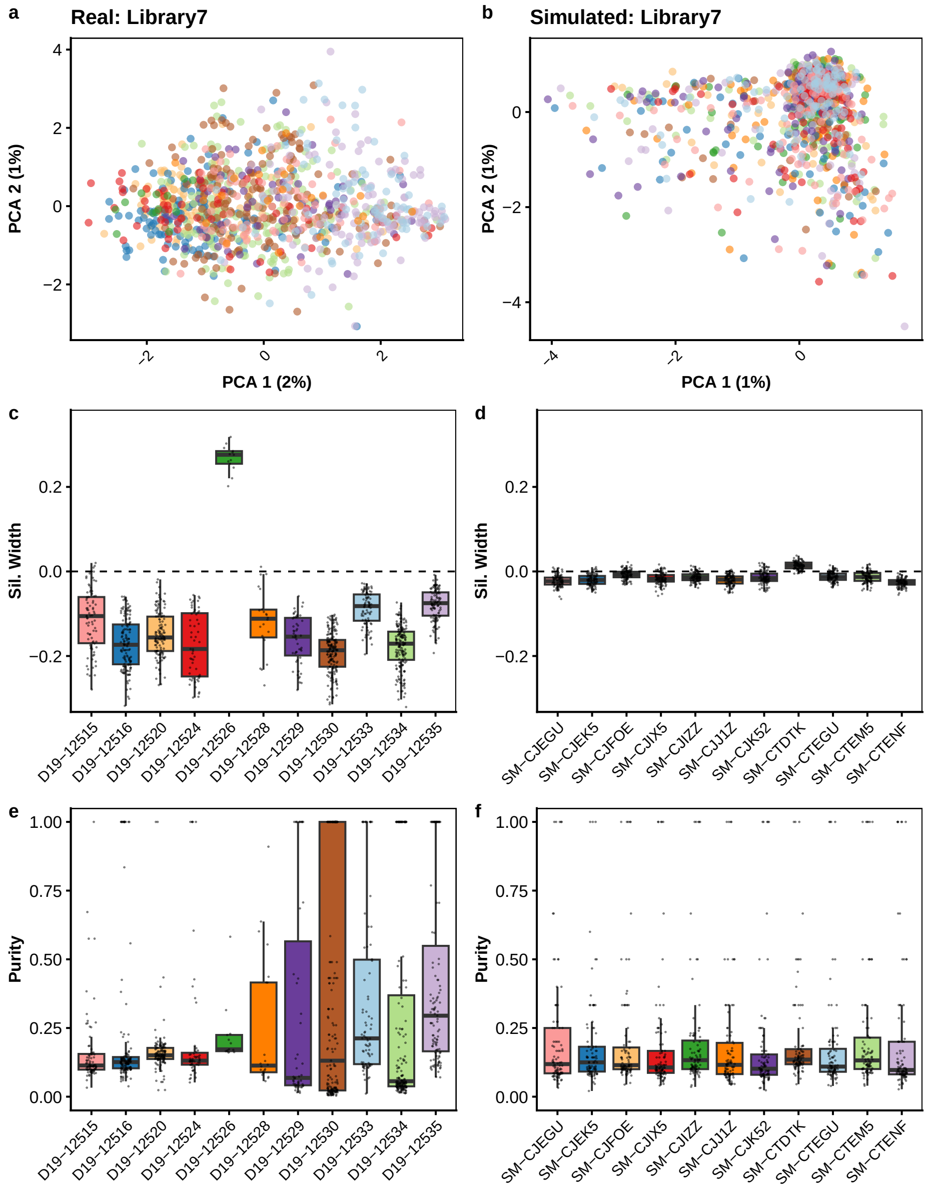


**Fig. S12** Quantifying clustering in real and simPIC-simulated data for non AD library 7 with 1693 peaks and 1012 cells. (Top row) PCA plot of real and simulated data, coloured by individual, showing the preservation of individual-specific structure. (Middle row) Silhouette width comparison between real and simulated data, assessing cluster compactness. (Bottom row) Neighbourhood purity analysis, evaluating local structure consistency across real and simulated datasets.

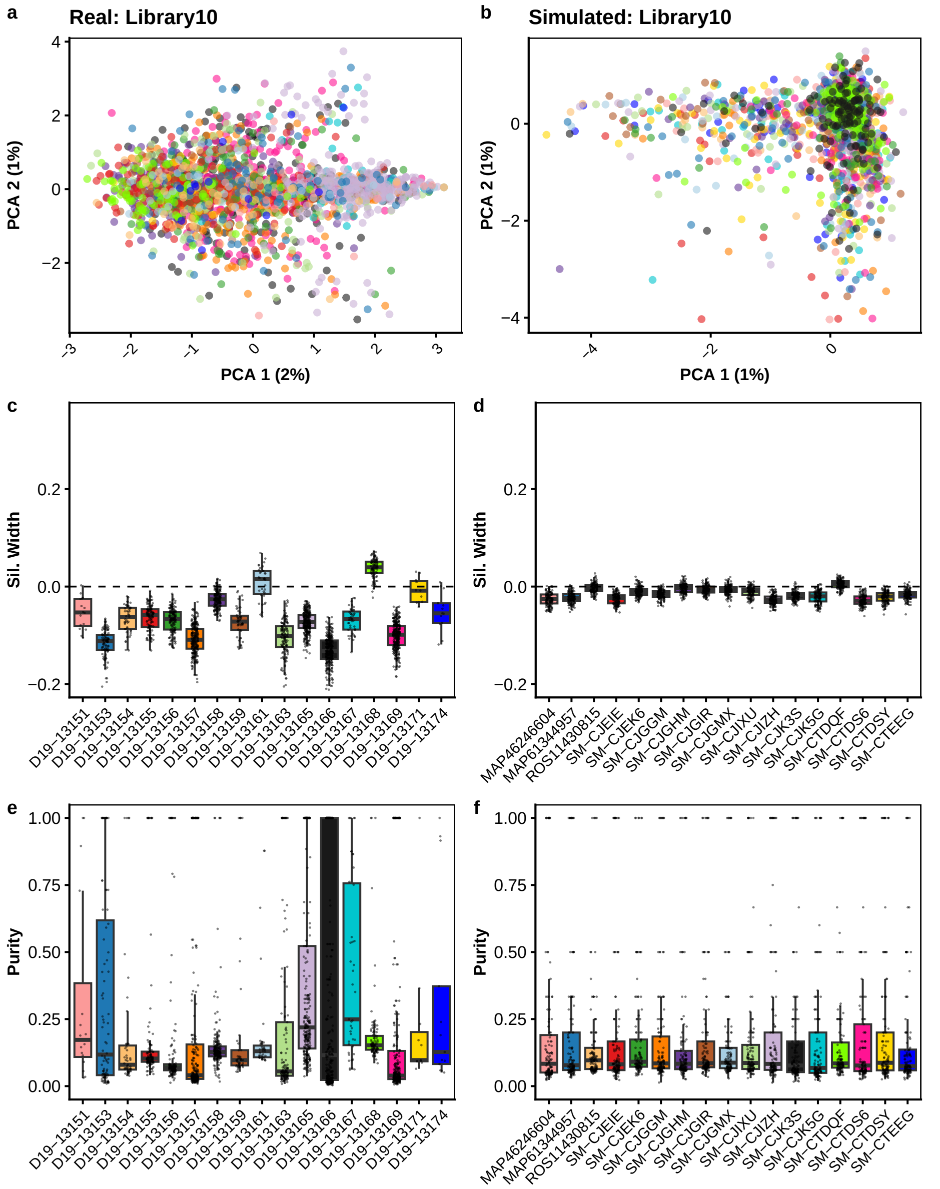


**Fig. S13** Quantifying clustering in real and simPIC-simulated data for non AD library 10 with 1693 peaks and 2057 cells. (Top row) PCA plot of real and simulated data, coloured by individual, showing the preservation of individual-specific structure. (Middle row) Silhouette width comparison between real and simulated data, assessing cluster compactness. (Bottom row) Neighbourhood purity analysis, evaluating local structure consistency across real and simulated datasets.

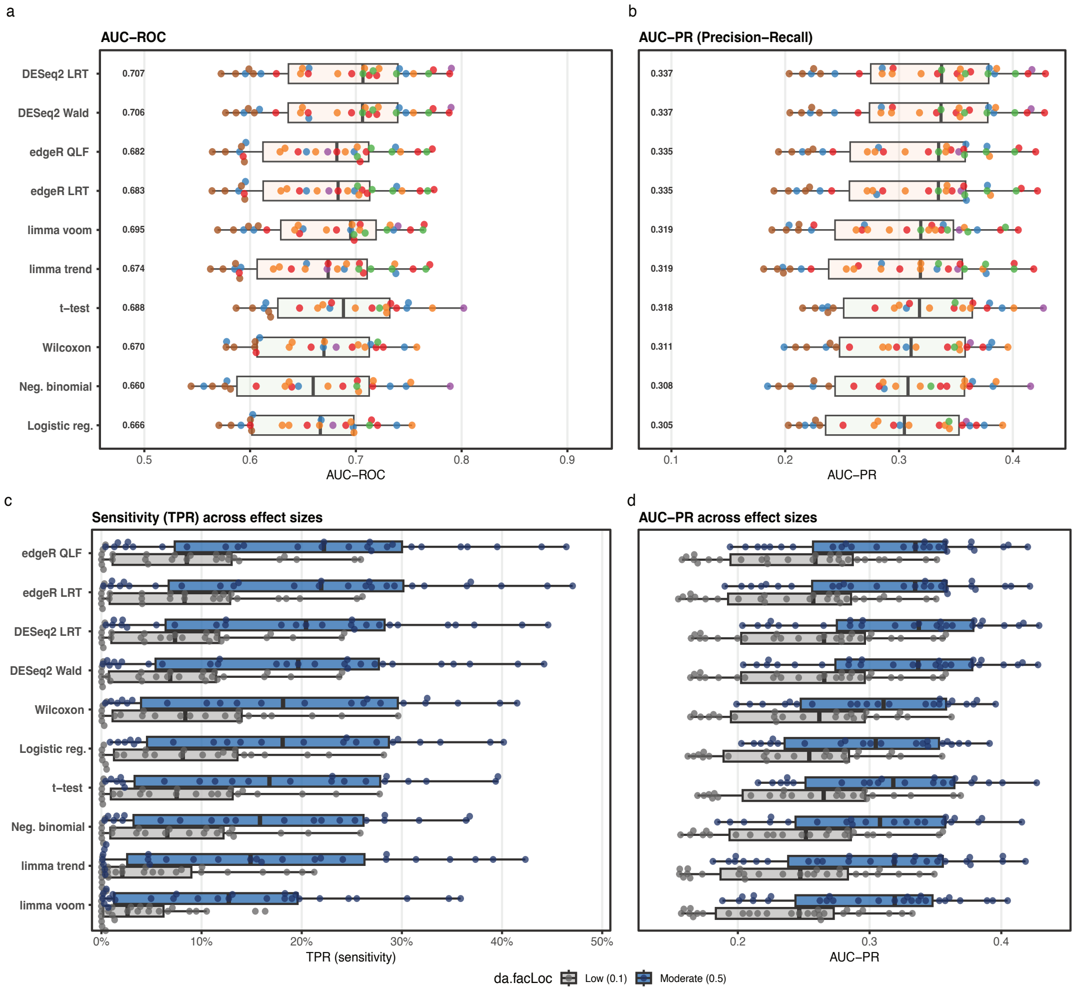


**Fig. S14** Performance of differential accessibility methods across simPIC simulated scATAC-seq datasets. (a) Area under the receiver operating characteristic curve (AUC–ROC) for each method across simulated datasets, summarising overall discrimination performance. (b) Area under the precision–recall curve (AUC–PR), highlighting performance under class imbalance typical of differential accessibility analyses. (c) Sensitivity (true positive rate, TPR) across effect sizes, comparing detection power under low (0.1) and moderate (0.5) effect sizes.(d) AUC–PR stratified by effect size, illustrating method robustness to increasing signal strength.

**Table S1**: Input parameters for SIMPIC simulation model.

| **Category** | **Parameter** | **Symbol** | **Description** | **Source** |
|---|---|---|---|---|
| Single-cell Parameters | Mean scale and shape | $\eta, \kappa$ | Scale and shape parameters for the Weibull distribution used to model peak means ($\lambda_i$). | estimated from input data |
| Single-cell Parameters | Library size location and scale | $\mu, \sigma$ | Parameters for the log-normal distribution characterizing the library size ($L_j$). | estimated from input data |
| Single-cell Parameters | Sparsity | $\pi_j$ | The proportion of zeros in the peak-by-cell matrix, modeled via a Bernoulli distribution ($Z_{i,j}$). | estimated from input data |
| Single-cell Parameters | Dispersion & Degrees of Freedom | $\phi, df_0$ | Common dispersion and degrees of freedom used to calculate the Biological Coefficient of Variation (BCV). | estimated from input data |
| Population Parameters | Population mean shape and rate | $\eta_p, \kappa_p$ | Weibull distribution parameters for generating peak means across a population. | estimated from input data |
| Population Parameters | Variance shape and rate | $\alpha_v, \beta_v$ | Gamma distribution parameters governing the coefficient of variation ($\sigma_i$) for sample-specific means. | estimated from input data |
| Population Parameters | caQTL effect shape and rate | $\alpha_c, \beta_c$ | Gamma distribution parameters for the chromatin accessibility QTL effect size ($\omega_i$). | estimated from input data |
| Manual & Batch Parameters | Batch factor location and scale | $\mu_b, \sigma_b$ | Log-normal parameters defining the batch effect ($\omega_j^b$) applied to cell means. | user defined; default provided |
| Manual & Batch Parameters | Group DA effect | $\pi_{da}, \mu_{da}, \sigma_{da}$ | Probability, location, and scale parameters for defining group-specific differentially accessible (DA) effects. | user-defined; default provided |
| Manual & Batch Parameters | Similarity scale | $S_s$ | A manual scaling factor influencing the population-wide variance. | user-defined; default provided |
| Manual & Batch Parameters | caQTL group specific | $caQTL_g$ | The percentage of caQTLs that are assigned as specific to a particular group. | user-defined; default provided |

**Table S2** Usability and practical comparison of scATAC-seq simulators.

| Dimension | simPIC | scDesign3 | scMultiSim | scReadSim | DiTSim |
|---|---|---|---|---|---|
| Required input | **Count matrix only** | Count matrix + cell annotations | Count matrix + GRN specification | BAM file + reference genome | Count matrix + cell annotations |
| Parameter estimation | **Automatic** `simPICestimate()` | Automatic | Manual / guided | Automatic from BAM | Deep-model training |
| Setup complexity | **Low** | Medium | High | High; BAM-processing pipeline | Medium–high; GPU recommended |
| Computational cost | **Low** | Moderate | High | High | High; deep learning |
| Bioconductor / CRAN | **Bioconductor** | GitHub / CRAN | Bioconductor | PyPI; Python | GitHub; Python |
| Interpretable parameters | ✓ | ✓ | Partial | ✗ | ✗; black box |
| Diagnostic / QC tools | ✓ `simPICcompare()` | ✓ | ✗ | ✗ | ✗ |
| Primary benchmarking use case | **Count-level analysis; population genetics / caQTL studies** | Multi-modal benchmarking | GRN, CCI and RNA velocity benchmarking | Read-level benchmarking; peak callers and UMI deduplication | Clustering and annotation |
| Population-scale capability | ✓ `splatPop` integration | ✗ | ✗ | ✗ | ✗ |
| Accessible to R-only users | ✓ | ✓ | ✓ | ✗ | ✗ |

**Key:** ✓ = supported; ✗ = not supported. Green ticks indicate supported capabilities; red crosses indicate absent capabilities. GRN, gene regulatory network; CCI, cell–cell interaction; caQTL, chromatin accessibility quantitative trait locus; UMI, unique molecular identifier.

**Table S3** Feature and scope comparison of scATAC-seq simulators.

| **Feature** | **simPIC** | **scDesign3** | **scMultiSim** | **scReadSim** | **DiTSim** |
|---|---|---|---|---|---|
| Modality focus | **scATAC-seq only** | Multi-modal; RNA, ATAC, spatial, CITE | Multi-modal; RNA, ATAC, velocity, spatial | scRNA-seq + scATAC-seq; read-level | scATAC-seq only |
| Output type | **Count matrix; peak × cell** | Count matrix | Count matrix | Raw reads; FASTQ / BAM | Count matrix; peak × cell |
| Statistical/modelling framework | **Gamma–Poisson / Weibull** | GAMLSS / copula | Kinetic ODE / GRN | scDesign2/3 internally | Diffusion transformer; deep learning |
| PIC-based quantification | ✓ | ✗ | ✗ | ✗ | ✗ |
| Multiple cell types | ✓ | ✓ | ✓ | ✓ | ✓ |
| Continuous trajectories | ✗ | ✓ | ✓ | ✓; via scDesign3 | ✗ |
| Batch effects | ✓ | ✓ | ✓ | ✓ | ✗ |
| Population-scale; multi-individual | **✓; via `splatPop`** | ✗ | ✗ | ✗ | ✗ |
| caQTL / genetic effects | ✓ | ✗ | ✗ | ✗ | ✗ |
| GRN-guided simulation | ✗ | ✗ | ✓ | ✗ | ✗ |
| Requires cell type annotations | ✗ | ✓ | ✓; cell states | ✗ | ✓ |
| Requires bulk ATAC data | ✗ | ✗ | ✗ | ✗ | ✗ |
| Sparsity modelling | **✓; Bernoulli** | ✓ | Partial | ✓ | ✓ |
| Read-level output; FASTQ/BAM | ✗ | ✗ | ✗ | ✓ | ✗ |
| Diagnostic / QC tools built in | **✓; `simPICcompare`** | ✓ | ✗ | ✗ | ✗ |
| Language | **R** | R | R | Python | Python |
| Bioconductor native | ✓ | ✗ | ✓ | ✗ | ✗ |

**Abbreviations:** PIC, Paired Insertion Counting; GRN, gene regulatory network; caQTL, chromatin accessibility quantitative trait locus; CCI, cell–cell interaction; ODE, ordinary differential equation; CITE, cellular indexing of transcriptomes and epitopes by sequencing.

**Table S4**: Cell types included in the simulator benchmark. For each dataset, the table reports the annotated cell type, number of cells, total number of peaks before filtering, and number of peaks retained after applying the 1% peak-detection filter.

| Dataset | Celltype | nCells | nPeaks | nPeaks (1%filter) |
|---|---|---|---|---|
| PBMC5k | CD14 Monocytes | 1768 | 69523 | 56279 |
| | CD4 Naive | 432 | 69523 | 45140 |
| | CD8 Effector | 505 | 69523 | 51493 |
| | Double negative T cell | 286 | 69523 | 46534 |
| | CD4 Memory | 641 | 69523 | 47538 |
| | CD8 Naive | 278 | 69523 | 43410 |
| PBMC10k | CD14 Monocytes | 2201 | 127689 | 106967 |
| | CD4 Memory | 2419 | 127689 | 83919 |
| | CD4 Naive | 1033 | 127689 | 78962 |
| | CD8 Effector | 866 | 127689 | 81813 |
| | B cell progenitor | 203 | 165434 | 97785 |
| | CD8 naïve | 440 | 165434 | 84291 |
| | NK dim | 760 | 165434 | 91594 |
| Satpathy | CD 34 Progenitors | 14633 | 571400 | 175151 |
| | Monocytes | 1868 | 571400 | 144533 |
| Buquicchio | Liver TRM | 4618 | 67364 | 64428 |
| Fly Brain | OL Type I NB | 323 | 51135 | 42926 |
| | OL Neuroepithelium | 274 | 51135 | 49296 |
| | Capa | 216 | 51135 | 42196 |
| | CB Type I NB | 205 | 51135 | 43759 |
| Cusanovich (Spleen) | B cells | 2197 | 73177 | 62219 |
| Cusanovich (Cerebellum) | Cerebellar granule cells | 1091 | 67260 | 37376 |
| | Astrocytes | 363 | 67260 | 36496 |
| Cusanovich (Kidney) | Podocyte | 447 | 101900 | 40426 |
| | Proximal Tubule | 2565 | 101900 | 85796 |

**Table S5** Summary statistics for real and simulated data
**(a) Peak usage and absolute value percentiles**

| Cell Type | Transform | Peaks Used | | P95 Abs | | P99 Abs | |
|---|---|---|---|---|---|---|---|
| | | Real | Sim | Real | Sim | Real | Sim |
| PBMC10k_CD8_effector | TFIDF | 2000 | 2000 | 0.080 | 0.070 | 0.112 | 0.092 |
| PBMC5k_CD4_memory | TFIDF | 2000 | 2000 | 0.312 | 0.187 | 0.737 | 0.455 |
| Cusanovich_Podocytes | TFIDF | 2000 | 2000 | 0.204 | 0.438 | 0.426 | 0.780 |
| Satpathy_Monocytes | TFIDF | 2000 | 2000 | 0.163 | 0.050 | 0.384 | 0.070 |
| FlyBrain_OL_neuroepi | TFIDF | 2000 | 2000 | 0.162 | 0.632 | 0.273 | 0.829 |
| Liver_TRM | TFIDF | 2000 | 2000 | 0.078 | 0.034 | 0.391 | 0.049 |

**(b) Pair counts and correlation thresholds**

| Cell Type | Transform | N Pairs | | Abs $r > 0.3$ | | Abs $r > 0.5$ | |
|---|---|---|---|---|---|---|---|
| | | Real | Sim | Real | Sim | Real | Sim |
| PBMC10k_CD8_effector | TFIDF | 1999000 | 1999000 | 63 | 0 | 6 | 0 |
| PBMC5k_CD4_memory | TFIDF | 1999000 | 1999000 | 104445 | 46953 | 56421 | 15093 |
| Cusanovich_Podocytes | TFIDF | 1999000 | 1999000 | 50800 | 120320 | 11225 | 87219 |
| Satpathy_Monocytes | TFIDF | 1999000 | 1999000 | 36053 | 38 | 9187 | 1 |
| FlyBrain_OL_neuroepi | TFIDF | 1999000 | 1999000 | 14322 | 199140 | 2012 | 142129 |
| Liver_TRM | TFIDF | 1999000 | 1999000 | 35606 | 26 | 9718 | 2 |

**Table S6**: Donor-associated separation metrics for real and simulated data across earlyAD and nonAD libraries.

| Condition | Library | Type | Mean silhouette | Mean purity |
|---|---|---|---|---|
| earlyAD | Library11 | real | -0.0120 | 0.5063 |
| | | simulated | -0.0002 | 0.3074 |
| | Library7 | real | -0.1261 | 0.2562 |
| | | simulated | -0.0092 | 0.2229 |
| | Library9 | real | -0.0246 | 0.5943 |
| | | simulated | 0.0118 | 0.5653 |
| | Library5 | real | 0.0051 | 0.5393 |
| | | simulated | 0.0171 | 0.4731 |
| | Library10 | real | -0.0581 | 0.3333 |
| | | simulated | -0.0070 | 0.2640 |
| | Library4 | real | 0.0247 | 0.6249 |
| | | simulated | 0.0118 | 0.5702 |
| nonAD | Library10 | real | -0.0816 | 0.2570 |
| | | simulated | -0.0153 | 0.1912 |
| | Library7 | real | -0.1448 | 0.2843 |
| | | simulated | -0.0139 | 0.1974 |
| | Library9 | real | -0.0320 | 0.3475 |
| | | simulated | -0.0004 | 0.3018 |
| | Library5 | real | 0.0728 | 0.5300 |
| | | simulated | 0.0114 | 0.3947 |
